# Multimodal alignments of *in vivo* imaging and spatial biology datasets at cellular resolution

**DOI:** 10.64898/2026.04.28.719500

**Authors:** Lun Wang, Xiqian Jiang, Xiaochen Sun, Gaurav Chattree, Ali Cetin, Xiaochun Cai, Elijah Paul, Radosław Chrapkiewicz, Oscar Hernandez, Yuxi Ke, Tomohisa Yoda, Fatih Dinc, Bariscan Kurtkaya, Yanping Zhang, Zhengji Zhang, Mark J. Schnitzer

**Affiliations:** CNC Program, Stanford University, Stanford, CA 94305, USA; Department of Biology, Stanford University, Stanford, CA 94305, USA; Department of Neurology & Neurological Sciences, Stanford University, Stanford, CA 94305, USA; Department of Applied Physics, Stanford University, Stanford, CA 94305, USA; Department of Neurosurgery, Stanford University, Stanford, CA 94305, USA; Department of Bioengineering, Stanford University, Stanford, CA 94305, USA; Koc University, Istanbul, Turkey; Howard Hughes Medical Institute, Stanford University, Stanford, CA 94305, USA

## Abstract

Parallel revolutions in intravital microscopy and spatial biology techniques have respectively enabled large-scale recordings of cellular dynamics in live animals and multi-dimensional molecular profiling at single-cell resolution. However, due to the challenges of aligning data from different modalities at cellular resolution, these two transformational approaches have generally been applied on separate biological samples, stymying the ability to link activity patterns and molecular attributes in the same exact cells. To enable routine, multimodal investigations of cells’ *in vivo* dynamics and molecular content, we created TRU-FACT (Total Registration Under Functional Activity, Connectivity, and Transcriptomics), a broadly applicable experimental and computational pipeline for registering large populations of individual cells across intravital imaging and spatial biology datasets. The pipeline combines three key innovations: an optomechanical tissue handling and alignment method to parallelize specimen planes, a graph-theoretic method to register individual cells based on their geometric relationships to neighboring cells, and a statistical framework that provides for each cell an *a posteriori* probability of correct registration. We validated TRU-FACT with several preparations for imaging neural Ca^2+^ activity in cortical and deep brain areas in head-fixed and freely behaving mice, RNA-barcode-expressing viruses for labeling neural projections, and low- and high-plex spatial transcriptomic methods. In mice performing a skilled reaching task, TRU-FACT alignments revealed the movement-related signaling patterns of intratelencephalic, extratelencephalic, and striatum-, superior colliculus-, and thalamus-projecting motor cortical neurons. Overall, TRU-FACT constitutes a scalable, multimodal discovery platform that is applicable to diverse tissue-types and spatial biology techniques, thereby enabling multiscale analyses of many complex biological systems.

## Introduction

Structure-function relationships, across length scales spanning molecules to organs, are central to much of biology research. Thus, in diverse fields such as developmental biology, physiology, immunology, cancer biology, and neuroscience, researchers have long sought to relate cells’ *in vivo* dynamics to their molecular architecture and organization in tissue^1–4^. To this end, powerful new techniques for spatial transcriptomic, epigenomic, metabolomic, or proteomic investigations have emerged over the past decade for revealing cells’ macromolecular content and the cytoarchitecture of tissue^5–7^.

For example, spatial transcriptomic methods based on RNA fluorescence *in situ* hybridization (FISH), such as single-molecule FISH^8,9^ and low-plex RNAscope^10^ or hybridization chain reaction (HCR) FISH^11,12^, have become commonplace for highly sensitive imaging of gene expression patterns at micron-scale resolution. More recent high-plex spatial transcriptomic techniques map the expression of up to tens of thousands of genes in cut tissue sections^13–17^. Spatial proteomic techniques are also advancing rapidly and use mass spectrometry and/or antibody panels to localize tens to hundreds of protein species in tissue specimens^18–20^. In turn, these innovations have sparked large-scale collaborations^21^ to build comprehensive atlases at cellular resolution of molecular expression patterns in mouse^22–25^, monkey^26,27^, and human brains^28–31^, as well as other organs^32,33^. Altogether, spatial biology studies are yielding important data about tissue development and architecture^4,34^, cell-type-specific functions^35,36^, cellular abnormalities under diseased conditions^37–40^, and new avenues for therapeutic development^30,41^.

Notwithstanding these exciting capabilities for revealing cellular and tissue architecture, key challenges remain for uncovering structure-function relationships, as no current approach allows biologists to reliably link the dynamics of large sets of individual cells in live plants or animals to the molecular attributes of these very same cells as revealed with spatial biology methods. A main hurdle is the technical difficulty of aligning dense populations of individual cells across intravital microscopy and postmortem image datasets. Thus, a reliable and broadly applicable way to integrate and register distinct categories of data across large sets of the same individual cells is sorely needed^42^. Without such an approach, many undiscovered structure-function relationships will likely stay hidden as cellular dynamics and structure remain imperfectly linked.

In neuroscience, the field in which we focused our experiments, the need to link the dynamics of single cells to their molecular attributes and organization in tissue is especially acute^43^. Individual neurons have wide-ranging activity patterns and computational roles in how the brain processes information and shapes animal behavior^44,45^. There are also hundreds of transcriptomic subclasses of neurons, with diverse connectivity and functional dynamics that are influenced but not solely determined by cells’ transcriptomic identities^22,23,46–49^. A major goal in the field is to determine how neurons’ genetic and connectomic attributes relate to their activity patterns and computational functions across a wide range of sensory, cognitive, and motor behaviors in health and disease.

Common ways to identify genetically- or projection-defined neuron-types in live animals include the use of transgenic animals and/or viruses to express fluorescent proteins in cells with specific genetic properties or axonal projections^50–52^. These approaches are usually limited by the modest number of fluorescent labels that can be expressed at once and reliably distinguished during imaging in live animals^53^. Thus, most *in vivo* imaging studies of neural activity track only one or a few fluorescently labeled cell classes; this limits efforts to study interactions between cell classes and how the array of neural subtypes with distinctive molecular or connectivity features sculpt brain dynamics.

In the absence of a way to identify many distinct cell-types in live animals, an emerging alternative involves cutting postmortem tissue slices from the regions imaged *in vivo* and then combining spatial biology and cell registration methods to classify at least a subset of the cells monitored while the brain was alive. Early work of this type relied on small numbers of postmortem markers and fluorescent labeling using immunostaining^54–57^, fluorescent protein^54,56–58^, or RNA FISH^59–63^ methods. More recent studies using cell registration employed greater numbers of genetic markers and FISH methods that allow more extensive multiplexing^60–64^.

Despite the major scientific advances that are achievable in principle by combining spatial biology and cell registration methods, cell alignments can be extremely challenging in practice. Substantial tissue deformations often occur during postmortem fixation and the cutting and mounting of tissue sections. Further, many optical systems used to image the live brain, such as microendoscopes for imaging deep regions or mesoscopes for imaging millimeters-wide areas, suffer from aberrations and field curvature that vary across the image field. Thus, to align images taken in the intact brain and postmortem sections, some studies confined cell registrations to sparsely labeled subsets of cells^59,61^, which can be aligned more easily than dense sets of labeled cells. Others used fluorescent markers^54,59,61^ or tissue landmarks such as blood vessels to guide image alignments^54,56,65^. To overcome misalignments, some studies relied on image processing or computational image reconstructions to correct for tissue deformations and optical aberrations^56,60,61^, which can be especially prominent when using microendoscopes for deep brain imaging^62,65^.

Notably, no study published to date has aligned neural activity at cellular resolution to high-plex spatial biology data with hundreds of markers. This is not just due to the difficulty of working with thin tissue sections (*e.g.*, ∼10 μm thick), as required for most high-plex spatial methods^10,13,14,23,66,67^. An equally important barrier is that high-plex image sets typically comprise large numbers of punctate labeled molecules and have very different visual appearances than images taken *in vivo*, greatly complicating approaches based on image alignment. Further, no study has provided a formal statistical framework to quantify cell registration fidelity and the likelihood any particular cell match was performed correctly.

Having statistics of this kind is crucial, owing to the challenges of optical and tissue distortions and the very different appearances, visual landmarks, and contrast levels of low- or high-plex spatial biology images as compared to images taken with intravital microscopy. Given these hurdles, the use of image-to-image alignments as a way to match cells across postmortem and *in vivo* datasets is a precarious endeavor, especially for densely labeled cell populations or large fields-of-view with spatially varying aberrations. To overcome the technical obstacles, it is vital to have a rigorous metric of the likelihood any individual cell was correctly registered. As our results show, the formulation and use of such a metric greatly facilitates registrations and affords statistical confidence in the results.

Beyond statistical issues, the use of specialized protocols and alignment algorithms tailored to the work of a small number of labs has hindered the field as a whole from routinely registering cells across *in vivo* and postmortem data. Bespoke methods have also made it hard for independent evaluators to assess registration fidelity and conclusions based on cell alignments. For instance, workflows with specialized tissue handling^60,62^, chemical labeling^54,60^, or fluorescence amplification^60,61,63^ steps can be difficult to standardize and transfer across labs, biological preparations, and imaging systems, leading to operator-dependent results. Collectively, overreliance on image alignment to match cells, lack of metrics to assess cell matching fidelity, and narrowly applicable lab protocols have stymied establishment of a general-purpose, scalable framework for high-fidelity cell matching across a wide range of *in vivo* microscopy and spatial biology platforms.

To integrate functional activity and spatial biology data at cellular resolution, we sought a widely applicable way to align individual cells across *in vivo* and postmortem spatial biology datasets. To this end, we created the TRU-FACT (Total Registration Under Functional Activity, Connectivity, and Transcriptomics) pipeline, which enables routine alignments at cellular resolution of neural activity imaging in live animals, mapping of long-range neural connections, and spatial biology data with low-or high-plex marker sets (**Fig. 1A**). To achieve this, TRU-FACT combines three innovations.

**Figure 1.**
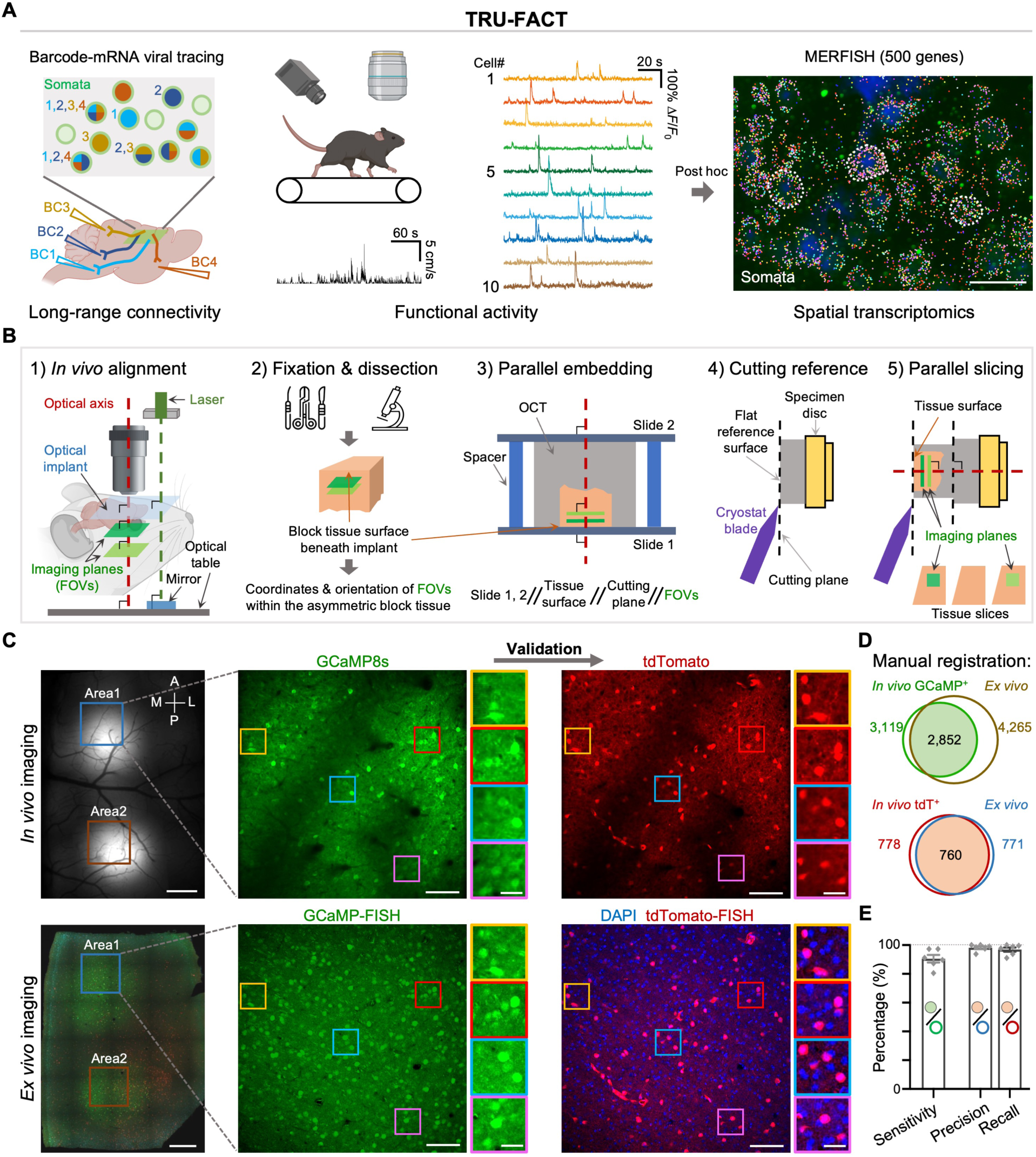
TRU-FACT accurately aligns large-scale movies of cellular activity acquired in live animals to *ex vivo* spatial biology datasets at a precision of individual cells. **A)** Schematic of the TRU-FACT pipeline to align multimodal image datasets, illustrated for an experiment combining two-photon neural Ca^2+^ imaging and spatial transcriptomics, as the *in vivo* and *ex vivo* imaging modalities, respectively. Other combinations of *in vivo* activity and spatial biology imaging methods are feasible. *Far left*: A mouse that expresses a fluorescent indicator in a brain area in which neural activity will be imaged *in vivo* is injected with retroAAV viruses in downstream target areas innervated by axons from neurons in the *in vivo* imaging field-of-view. Each target area receives a retroAAV expressing a unique barcode (illustrated here for 4 barcodes, BC1–BC4); this allows neurons in the *in vivo* imaging field to be labeled according to their axonal projections. Individual cells may express none or even more than one barcode. *Middle left*: The mouse is head-fixed under the objective lens of a two-photon microscope and performs a behavioral assay, *e.g.*, running in place or pellet reaching. *Middle right*: During active mouse behavior, neural Ca^2+^ dynamics are monitored at cellular resolution by two-photon microscopy. *Far right*: After *in vivo* imaging, transcriptomic and projectomic cell-types are determined for the same individual cells that were monitored by Ca^2+^ imaging. Such cell classifications can be determined with a variety of methods; shown here is an illustrative MERFISH image of a primary motor cortical tissue slice. This slice was labeled using a 500-gene Mouse PanNeuro Cell-Type panel; each dot marks a single RNA transcript. White dashed lines enclose 3 example neuronal cell bodies. Scale bar: 20 μm. **B)** Key steps for aligning brain tissue planes imaged in a live animal to the tissue slices used for low-or high-plex postmortem spatial biology analyses. **Methods** and **Fig. S1** detail three variations of our alignment strategy; the third and most general version used in our work is schematized here. **(1)** During *in vivo* imaging, we aligned the optical axis to be orthogonal to the outward-facing surface of a glass implant, *e.g.*, a cranial window, microendoscope, or microprism, thereby establishing a volumetric set of imaging planes in tissue that all lie parallel to each other. The cartoon illustrates this for the case of a cranial window. **(2)** After *in vivo* imaging, we perfused the animal to fix the brain tissue while the glass implant remained intact. After perfusion, we dissected a tissue block containing the region of *in vivo* imaging and removed the glass implant. Throughout this process, with each step we progressively updated records of the *in vivo* imaging field-of-view using prominent landmarks such as blood vessels and edges of the tissue images. **(3)** We embedded the tissue in OCT (optimal cutting temperature) compound and sandwiched the OCT block between two glass slides. To ensure that both faces of the OCT block and the set of planes imaged *in vivo* all were parallel to each other, we placed the tissue surface that had abutted the glass implant *in vivo* onto the bottom slide and used two sets of spacers of identical net height to ensure the second slide was parallel to the first. **(4)** We created a flat reference surface in another (empty) OCT block by cutting it in a cryostat parallel to the blade’s cutting plane. **(5)** We attached the OCT block containing the tissue block to this reference surface of the empty OCT block, ensuring that the plane of the cutting, the tissue block surfaces, and the planes imaged *in vivo* were all mutually parallel. We then cut tissue sections with the same blade. **C)** To evaluate the alignment accuracy of TRU-FACT, we used it to align pairs of *in vivo* and *ex vivo* images of dense sets of cortical pyramidal cells expressing a green fluorescent Ca^2+^ indicator (GCaMP8s). We quantified alignment accuracy using a subset of these cells that co-expressed a red fluorescent marker (tdTomato) and that was sufficiently sparse to allow nearly unambiguous visual alignments of the *in vivo* and *ex vivo* datasets. For these experiments, we expressed GCaMP8s in the motor (Area 1) and somatosensory (Area 2) cortices via local injections of AAV2/PHP.eB-CaMKII-jGCaMP8s virus. Mice (Drd1a-Cre ✕ Ai14) also expressed a Cre-dependent tdTomato label in a sparse subset of cells expressing the D1-dopamine-receptor. Panels show example image pairs taken in an awake resting mouse (*top row*) and with *ex vivo* HCR-FISH and confocal fluorescence imaging (*bottom row*), as aligned by TRU-FACT. *Left*: Tiled image mosaics of GCaMP8s fluorescence (*top*), acquired by epi-fluorescence microscopy through the mouse’s cranial window, and HCR-FISH in a 100-μm-thick slice (*bottom*). The two areas are enclosed in colored squares; magnified views of Area 1 are shown in the middle and right columns. **Fig. S2E** has images from Area 2, acquired at 3 tissue depths spanning cortical layer 2/3 to layer 5. Scale bars: 500 μm. *Middle*: Aligned pair of *in vivo* two-photon (average of 3,000 image frames of GCaMP fluorescence taken at 30 fps) and *ex vivo* images of cells 290 μm beneath the brain surface. The confocal image shows Alexa-488 dye labeling of GCaMP mRNA. Colored squares enclose areas shown at higher digital magnification in the color-corresponding insets at right. Scale bars: 100 μm. *Insets*: 4 aligned pairs of magnified images. Scale bars: 30 μm. *Right*: Aligned pair of *in vivo* two-photon (average of 3,000 frames of tdTomato fluorescence taken at 30 fps) and *ex vivo* images, over the same tissue areas as in the middle column. The *ex vivo* image shows Alexa-594 dye labeling of tdTomato mRNA, plus DAPI staining (blue) used for cell segmentations. *Insets*: Magnified views of the areas enclosed by the color-corresponding squares in the middle and right columns. **D, E)** Venn diagrams, **D**, showing results of a validation study involving manual cell segmentation and registration across 6 different planes imaged *in vivo* in 2 mice, including the example images of **C**. We initially performed all cell registrations using only green fluorescence. We then validated the alignment results for the green channel by performing a separate set of alignments on the sparse set of cells that expressed red tdTomato, for which human visual alignments could be trusted due to the sparseness of the red-labeled cells, providing a set of ground truth data. We detected a total of 3,119 green fluorescent neurons *in vivo*, of which we found 2,852 again among the 4,265 green fluorescent cells found *ex vivo*. For the sparse set of 778 red fluorescent cells observed *in vivo*, we found 760 of them among the 771 red fluorescent cells observed *ex vivo*. Bar plots, **E**, showing quantifications of the results, plotted as mean ± s.e.m. over n=6 pairs of corresponding image planes. Gray diamonds denote results from individual image planes. The sensitivity, defined as the percentage of green cells found *in vivo* that were also found *ex vivo,* was 91%. The precision, defined as the percentage of red cells found *ex vivo* that were also found *in vivo,* was 98 ± 1%. The recall, defined as the percentage of red cells found *in vivo* that were also found *ex vivo,* was 97 ± 2%. The high values of both the precision and recall verify our abilities to visually align the densely labeled green fluorescent cells.

First, it uses an optomechanical strategy to parallelize imaging and tissue-cutting planes across all tissue handling steps and the *in vivo* and spatial biology microscopy setups. To align *in vivo* and postmortem datasets, it is advantageous to cut tissue slices as parallel as one can to the optical planes imaged *in vivo*; this eases cell registrations by reducing the need for computational image corrections. The advantage is especially notable when using thin (∼10 μm) tissue sections, but even for transcriptomic^11,16,68–70^ or proteomic^71–74^ studies of thick sections it remains beneficial to align tissue sections to the *in vivo* imaging planes, due to tissue distortions, optical aberrations, and the challenge of aligning image stacks from different microscopy modalities. TRU-FACT starts with an optomechanical workflow to track and transfer parallel sets of imaging planes across different microscopes, facilitating trustworthy alignments even when cells are densely labeled and in thin tissue sections. We present three variants of this workflow (**Fig. S1**), all based on the core idea of maintaining a physical record of the planes imaged *in vivo*, either by making a mold of the brain shape or by retaining the shape of the tissue surface contacting the glass implant used for *in vivo* imaging. All three variants are straightforward to execute in practice, facilitating routine, accurate alignments.

Second, TRU-FACT uses a graph-theoretic algorithm to register cells based on their geometric relationships to neighboring cells. Even if tissue sections are cut in an ideal manner, reliable cell registration across datasets taken with different microscopes, contrast modalities, and distortions remains a substantial computational challenge. Unlike predecessor algorithms^60–62,75,76^, the cell alignment algorithm in TRU-FACT does not rely on overall image similarities or the spatial distribution of image contrast. Instead, we create a Euclidean graph representation of each cell’s relative locations to its neighbors, termed a ‘Soma-print’, and align cells by identifying candidate cell pairs with matching Soma-prints, *i.e.*, highly similar graph representations. The rationale is that the Soma-print is designed to capture a robust, distinctive signature of each cell that is impervious to visual differences in how cells appear in images taken with different contrast mechanisms^77–79^. If one considers a sufficient number of neighboring cells, typically 10 or more neighbors per cell, the Soma-print algorithm generally provides trustworthy cell matches across postmortem and *in vivo* datasets. The algorithm works well with either 2D or 3D image data, is insensitive to moderate tissue distortions or cutting misalignments, scales well to mesoscope images with large numbers of cells, and is applicable to a wide variety of *in vivo* and postmortem imaging approaches. Further, the Soma-print algorithm typically takes minutes to run, far quicker than the hours often required for the painstaking process of matching hundreds of cells through human visual inspection.

Third, TRU-FACT includes a statistical framework that provides for each cell match an *a posteriori* probability that the match is correct. Each match comes with a statistical p-value, estimated directly from the data, and the relative likelihoods of an incorrect *vs.* a correct match, enabling rigorous assessments of cell registration fidelity and any resulting biological conclusions.

We validated TRU-FACT with several rodent preparations and microscopes commonly used for two-photon Ca^2+^ imaging of neural activity. These included a conventional two-photon microscope, combined with a cranial window or an implanted microendoscope or microprism for cortical or deep brain imaging, respectively, as well as a two-photon mesoscope for Ca^2+^ imaging over a ∼4 mm^2^ tissue area and a miniature two-photon microscope for Ca^2+^ imaging in freely behaving mice. We further verified that TRU-FACT works well with low- and high-plex spatial biology approaches, using HCR-FISH^11,12^ and MERFISH^13^, respectively. (See **Supplementary Video 1**, which shows 5 example alignments across a range of experiments). Testifying to TRU-FACT’s versatility and ease of use, the data here are from 13 mice and 5 different *in vivo* imaging preparations, allowing matching of 10,522 individual cells across *in vivo* and postmortem datasets. These values constitute more mice and more optically matched cells than reported in prior studies of the mammalian brain^54,56–63,80^. Overall, TRU-FACT constitutes the first broadly applicable way to register densely labeled cells across *in vivo* and postmortem image datasets using thin or thick tissue sections.

In addition to Ca^2+^ activity, we sought to relate neurons’ axonal targets to the RNA expression profiles of the same individual cells. For this, we created 8 retrograde adeno-associated viruses (retroAAVs), each expressing a unique RNA barcode. After injecting one or more of these viruses in brain areas innervated by axon terminals of neural cell bodies within the *in vivo* imaging field-of-view, spatial transcriptomic methods can be used to identify the barcodes postmortem and reveal individual cells’ long-range projection targets. We applied these joint alignments of Ca^2+^ activity, connectivity, and RNA profiles to study projectomic and transcriptomic classes of motor cortical neurons in mice doing a skilled reaching task. In this way, we characterized movement-evoked Ca^2+^ dynamics for intratelencephalic, extratelencephalic, and striatum-, superior colliculus-, and thalamus-projecting motor cortical neurons, showing how TRU-FACT can forge links between animal behavior and the underlying cell-types, neural activity patterns, and circuit attributes.

Although our experiments here focused on the mouse brain, the broadly applicable principles of the TRU-FACT pipeline should make it readily adaptable to other cell-types, tissues, and species across multiple length scales and diverse biological fields. A Soma-print’s insensitivity to the image contrast mechanism makes our alignment pipeline suitable for use with other means of monitoring cellular dynamics beyond Ca^2+^ imaging. The pipeline is compatible with many different spatial biology platforms, including a range of transcriptomic, proteomic, metabolomic, or epigenomic imaging modalities. Overall, TRU-FACT provides a unified framework for integrating molecular, anatomical, and functional data from the same individual cells. This in turn enables a scalable, multimodal approach to biomedical discovery with diverse tissue-types, spatial biology techniques, and organism species in health and disease.

## Results

### Optomechanical alignment of *in vivo* imaging planes with postmortem tissue sections

The first key innovation of TRU-FACT is an optomechanical process to ensure that the *ex vivo* imaging planes, which are usually defined by the mechanical planes for cutting postmortem tissue sections, are parallel to the optical focal planes examined by intravital microscopy in the same animal (**Figs. 1B; S1A–C**). This alignment process is general-purpose and should be applicable to different organs and animal preparations for *in vivo* imaging. As neuroscientists, we focused on three preparations commonly used in brain research, the cranial window preparation for imaging the neocortex^81^, and deep tissue imaging preparations in which either a microprism^82,83^ or a GRIN microlens^84,85^ is implanted in the brain.

Past studies that sought to register individual cells across *in vivo* and *ex vivo* imaging datasets used a variety of mechanical or optical procedures to align the two sets of planes^55,56,59–62,65,80^. What distinguishes our approach from past efforts is a complete workflow that starts with a laser-based alignment of the optical axis of the microscope used with live animals, employs an analogous procedure to align the animal’s glass implant (*viz.*, a cranial window, microprism, or GRIN lens) at the start of *in vivo* imaging, and then involves a postmortem, optomechanical parallelization procedure to transfer the precise orientation of the glass implant to a parallel plane in the tissue embedding medium (*e.g.*, OCT or agarose) used to cut tissue sections. Once this parallel plane is created in the embedding medium, a final mechanical step transfers this plane’s orientation to yet another plane, namely the cutting plane of the blade used to slice tissue sections.

We present three variations of the optomechanical strategy, two that involve casting a mold capturing the shape of the brain’s exterior surface (**Fig. S1A, B**), and one that relies solely on the flatness and integrity of the fixed tissue touching the glass window, GRIN microlens, or microprism (**Figs. 1B** and **S1C**). The common idea underlying each of these is that the form of the tissue surface that had touched the glass implant provides a physical record of the pitch and roll coordinates of the *in vivo* imaging planes, provided these planes were optomechanically parallelized to the implant prior to imaging. The choice of which of the three optomechanical variations to select is dictated by the user’s level of expertise, the thickness of and other requirements regarding the cut tissue sections, and the details of the *in vivo* imaging preparation. The first variation is recommended for beginners, the second for applications in which the tissue should not be frozen postmortem, and the third for most other use cases (see **Methods** for details and further guidance). Across the data shown in the rest of this paper, citations to **Fig. S1A, B**, or **C** indicate which variation we used in each case. In addition to the procedures designed to ensure parallelization of the imaging planes, photographs and cellular-level images taken at each step allow users to record the tissue appearance, anatomic landmarks, and locations and orientations of the cell arrangements imaged *in vivo* and postmortem.

Altogether, our workflow allows reliable tracking and alignment of tissue across 6 mechanical degrees-of-freedom (3 rotational, 3 translational) throughout the acquisition of *in vivo* and *ex vivo* imaging datasets. As illustrated below, each of the variations allowed us to consistently relocate individual cells across *in vivo* and *ex vivo* imaging datasets that differed substantially in appearance. This, in turn, enabled our goal of integrating large-scale neural recordings with *ex vivo* projectomic and transcriptomic data for the same individual cells.

### Quantitative benchmarking of optomechanical alignment fidelity

We first sought to evaluate the fidelity with which densely labeled neurons could be registered across *in vivo* Ca^2+^ imaging and *ex vivo* transcriptomic datasets using our optomechanical alignment process. In general, most past studies aligning Ca^2+^ imaging and spatial transcriptomic datasets did not perform rigorous benchmarking of alignment fidelity and instead relied on computational image registrations and post hoc visual assessments of cellular and image appearances^55,56,59–62,65^. In many cases, these visual verifications were greatly eased by focusing selectively on sparsely labeled subsets of cells, such as sparse subsets of interneurons^54,56,57,59,61^.

To benchmark our alignment fidelity with densely labeled cells, we performed a statistical test involving dual-color fluorescent mice, in which a dense set of neocortical neurons expressed a green fluorescent Ca^2+^ indicator, and a very sparse subset of these cells co-expressed a red fluorescent marker. Due to the sparsity of the red neurons, we reasoned they could be faithfully registered across *in vivo* and *ex vivo* datasets through human visual alignments, and that these cross-modality assignments for the red cells could be used to compute quality control metrics for a far more challenging set of visual alignments performed separately for the dense set of green cells.

To execute this test, we used Drd1a-Cre × Ai14 double transgenic mice in which a red fluorescent tdTomato marker is confined to neurons expressing the D1-dopamine-receptor^86^. In these mice, we also expressed green fluorescent GCaMP8s in neocortical pyramidal cells by locally injecting AAV2/PHP.eB-CaMKII-jGCaMP8s virus; this yielded a dense set of green-labeled pyramidal cells plus an overlapping sparse subset of cells that expressed the red marker. Throughout, we used the optomechanical alignment procedure of **Fig. S1B** and microscope objective lenses with minimal field curvature and thus flat focal planes that can be parallelized to each other with micron-scale precision (**Fig. S2A, B**). After one or more sessions of two-photon Ca^2+^ imaging in awake mice over 700 × 700 μm^2^ fields-of-view, fixation of the brain, and cutting of 100-μm-thick tissue sections, we performed postmortem, low-plex spatial transcriptomic analyses using hybridization chain reaction fluorescence *in situ* hybridization (HCR-FISH^11,12^) (**Fig. S2C**). Due to the precision of the optomechanical steps, after taking confocal images of the tissue sections labeled with HCR-FISH probes of GCaMP and tdTomato, we were able to visually align corresponding pairs of *in vivo* and postmortem images using the densely labeled GCaMP green channel (**Figs. 1C** and **S2D, E**). We then visually registered the green-labeled cells across the *in vivo* and HCR-FISH datasets and evaluated the quality of cell registration by checking the alignments against those obtained independently for the cells expressing tdTomato.

Of the 3,119 GCaMP-labeled cells detected *in vivo* across 6 different planes imaged at one of two different brain areas in each of two mice, we re-identified 2,852 cells in postmortem tissue sections, for an overall sensitivity of >91% (**Fig. 1D**). To evaluate the fidelity of alignments performed with the green cells, we checked the alignment results against those obtained separately for the sparsely labeled tdTomato-positive cells. In the cut tissue sections, we re-identified 760 of the 778 red cells observed *in vivo* (**Fig. 1D**), yielding precision and recall values of 98 ± 1% and 97 ± 2% (mean ± s.e.m.; n=6 planes; **Fig. 1E**), respectively defined as the percentages of red cells in the postmortem and *in vivo* datasets that were re-identified in images of the other modality. With these results in hand, we then re-used the datasets as a source of ground-truth results when creating the Soma-print algorithm for automated cell alignments.

### Soma-print: A cellular fingerprint for registering cells across imaging modalities

The above evaluations of our optomechanical alignments relied on human visual image registration, as in many past studies^56,59–63^. However, this time-consuming approach plainly does not scale well to large datasets. Further, even with our optomechanical procedures, cell registrations by human annotators are prone to perceptual errors and inter-human variability and also suffer from the more fundamental problem that they lack a rigorous statistical metric characterizing the likelihoods the cell assignments are correct. Past studies that used computational methods as part of the cell registration process also lacked statistical metrics of cell alignment fidelity^60–63^. For these reasons, we sought an automated algorithm that could not only align individual cells across datasets but also provide for each proposed cell match a p-value, an *a posteriori* probability, and a likelihood ratio characterizing the relative odds that the match was incorrect *vs.* correct.

As we designed such an algorithm, we wanted to ensure it could accurately align cells across images taken with different microscopy modalities, without relying on the fine details of image appearances. Notably, registration methods that rely on hallmark image features are vulnerable to errors when the appearances of cells and surrounding tissue are not preserved well across *in vivo* and postmortem images^87^. More broadly, cell registration methods that use metrics of spatial overlap^88^ are prone to false-positive matches, due to spatial distortions that often occur in postmortem tissue-processing (**Fig. 2A** shows an example) or simply due to optical aberrations^59,62^.

**Figure 2.**
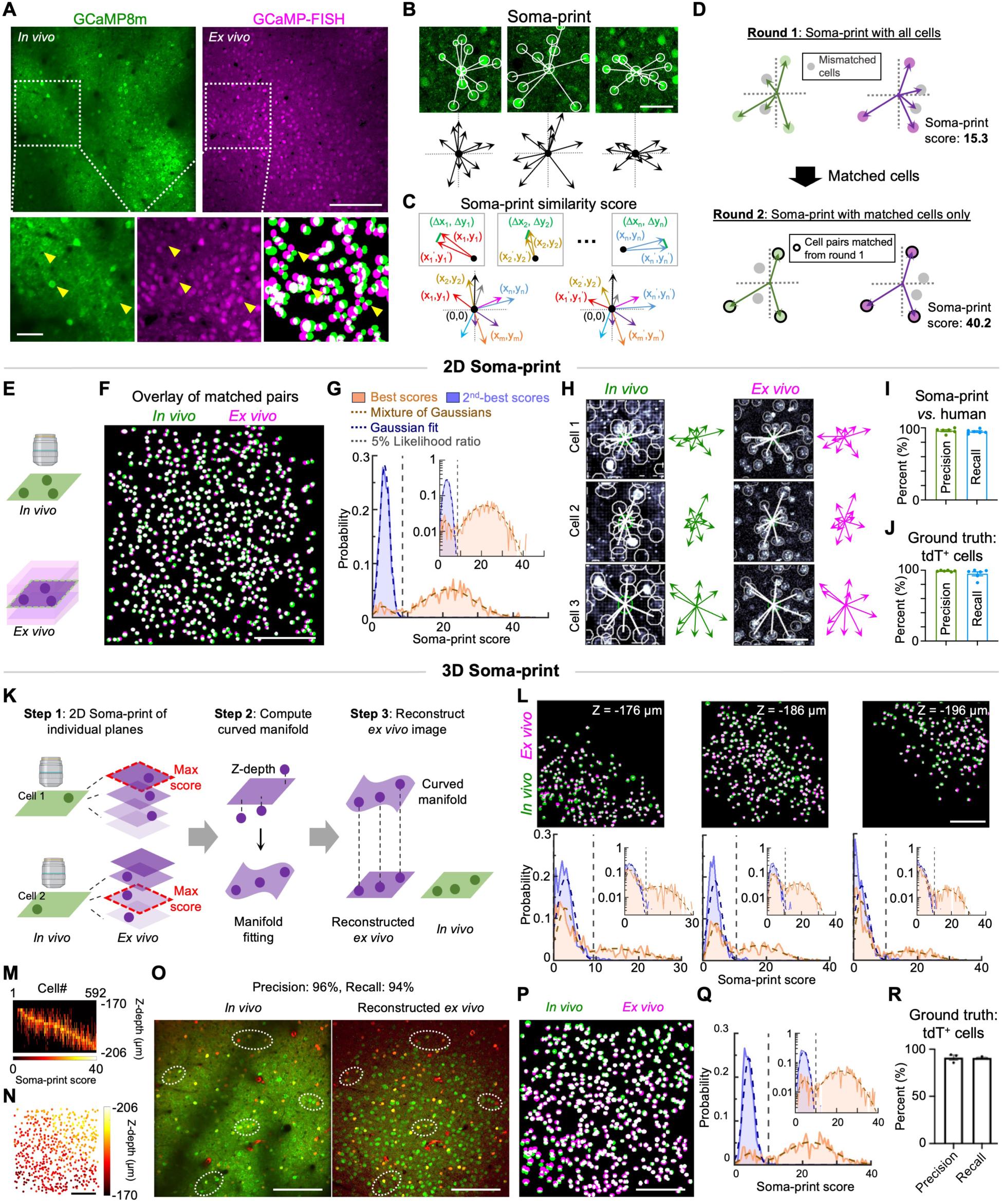
The Soma-print algorithm is a general method to register cells’ identities across images obtained with different microscopy modalities. Registration of individual cells can be challenging when the images involved come from different forms of microscopy yielding images of very different appearances. To overcome this hurdle, the Soma-print algorithm relies on a set of characteristic vectors for each cell, extending from the centroid of a cell’s soma to the centroids of neighboring cells. In this way, the Soma-print is a high-dimensional metric that captures the distinctive geometric relationships between an individual cell and its neighbors. **A)** *Left*: Example *in vivo* two-photon image of pyramidal cells expressing the GCaMP8m Ca^2+^ indicator in the motor cortex (a time-averaged image of 3,000 frames taken at 30 fps at a depth of 250 μm from the pia surface). *Right*: *Ex vivo* image of the same tissue plane, labeled with a fluorescent Alexa-488-conjugated HCR probe to GCaMP mRNA. White squares enclose areas shown at higher magnification in the insets below. Scale bar: 200 μm. *Insets*: A pair of magnified images (*left* and *center*), plus an overlay (*right*) of the cell maps determined from the *in vivo* and *ex vivo* images. Yellow arrowheads mark matched cells for which spatially non-uniform distortions in the postmortem tissue (*e.g.*, due to shrinkage and shearing during tissue processing) lead to imperfect overlaps in the overlaid cell maps. Scale bar: 50 μm. **B)** *Top row*: *In vivo* images of 3 example GCaMP-expressing cells, each shown with n=10 neighboring cells that are encircled with a white perimeter and connected to the cell of interest by a vector (white lines). *Lower row*: Soma-prints for the 3 corresponding cells above. Each of the 10 vectors for each cell is defined by the centroid coordinates of a neighboring cell body, (x_k_, y_k_), relative to the centroid of the cell of interest. Scale bar: 50 μm. **C)** Schematic showing how Soma-print similarity scores are computed (*top row*) for a pair of cells to assess the concordance of their Soma-prints (*bottom row*). The score is computed by finding the mean Euclidean distance (green lines, *top row*) of the vector differences between the *n* pairs of matched vectors across the two Soma-prints (*i.e.*, the pairs of red-, tan-, or light blue-labeled vectors). Given the *n* vectors in each Soma-print, the algorithm sets the *n* pairings in a greedy manner from larger sets of *m* vectors (*m* > *n*), via an iterative process in which the two best-matched vectors are determined and then removed from consideration in further iterations. The mean Euclidean distance, computed across all the matched pairs of vectors, is then converted into a Soma-print similarity score ranging from 0–100, with a higher score indicating greater similarity (**Methods**). **D)** Schematic showing the Soma-print algorithm’s multi-round design for matching *in vivo* (schematized in green) and *ex vivo* (magenta) datasets. *Upper*: In the first round, each cell’s Soma-print is computed based on the positions of *n* nearest neighbors in the field-of-view. Cells that are well-matched across the two imaging modalities are shown in color, whereas cells lacking good matches are shown in gray. *Lower*: In subsequent rounds of computation, each cell’s Soma-print is recomputed by restricting the selection of neighboring cells to only those that were successfully matched across imaging modalities in prior computational rounds (schematized as cells with bold perimeters). This restriction progressively increases the Soma-print similarity scores of cells that had mismatched neighbors in earlier rounds (**Fig. S3F**). **E)** In the 2D Soma-print algorithm, cells from an *in vivo* image (*top,* green) are matched with cells found in a single image slice (*bottom*, purple) within the stack of images acquired *ex vivo*. **F)** Cell map from one of the planes used to compute the results of Fig. 1E, showing the overlay of pairs of cells matched together using the Soma-print algorithm from *in vivo* (green) and *ex vivo* (magenta) cell maps. Overlapping pixels in both maps are shown in white. Scale bar: 200 μm. **G)** Distributions of Soma-print scores for pairs of best-matched cell pairs (orange) or 2^nd^-best-matched cell pairs (blue), computed for the image of **F**. Parametric fits (dashed curves) used a Gaussian function to fit the distribution of scores for 2^nd^-best matches, or a mixture of two Gaussians to fit the distribution of scores for best-matched cell pairs. Using these fits, we computed the likelihood ratio that a given Soma-print score reflects a 2^nd^-best match *vs.* a best-match. In practice, we only accepted matches for which this likelihood ratio was <0.05 (vertical dashed line marks this 0.05 cutoff value). For the data of **F**, 621 of the 704 cells found *in vivo* had corresponding matches among the 979 cells found in *ex vivo*. *Inset*: The same distributions and fits are shown on a semi-logarithmic plot. **H)** 3 example pairs of cells from **F** that were matched across the *in vivo* and *ex vivo* images, shown with their cell soma boundaries enclosed in white and corresponding Soma-prints (n=10 vectors per Soma-print). Scale bar: 40 μm. **I)** We compared the alignment precision (96 ± 1%; mean ± s.e.m. over the n=6 corresponding pairs of images used in Fig. 1C**–E**) and recall (95 ± 1%) values for cell pairs matched using the Soma-print algorithm, against the precision and recall values for cell pairs matched by human visual alignments of the densely labeled green fluorescent images. Notably, the Soma-print algorithm took minutes to run for each image pair, whereas the process of matching the same sets of cells by human visual inspection took many hours. **J)** To evaluate the quality of cell matching with the Soma-print algorithm, we first ran the algorithm on the dense sets of green-labeled cells in the dataset of Fig. 1C**–E**. We then performed a quality check on the algorithm’s results using the sparsely labeled subset of red (tdT^+^) fluorescent cells, which were sparse enough for us to reliably match cells through visual inspection. Taking the matches from the sparsely labeled subset as the ground truth, the precision and recall values for the matches determined by the algorithm were (99% ± 0.4%; mean ± s.e.m. over the n=6 image pairs) and (95 ± 3%), respectively. (See Fig. 1E for definitions of alignment precision and recall). **K)** Schematic showing the workflow of the 3D Soma-print algorithm. We used this algorithm when the *in vivo* imaging plane was not flat, *e.g.*, due to optical curvature of the imaging field, or when the mechanical slicing of tissue sections was not perfectly parallel to the *in vivo* imaging plane. This version of the algorithm allows cells imaged *in vivo* (shown as green circles) to be matched with corresponding cells (purple circles) distributed across a stack of images acquired *ex vivo*. Individual cells observed *in vivo* are first matched to cells in the individual *ex vivo* image planes using the 2D Soma-print algorithm. Each cell from the *in vivo* data with a match in one or more planes is then registered to a single plane within the stack of *ex vivo* images by determining the plane that yields the cell match with the highest 2D Soma-print score. This information is then used to compute a 2D curved manifold that is an estimate of the shape of the *in vivo* imaging field-of-view. Finally, a flat 2D *ex vivo* image is reconstructed for this estimated field-of-view by sampling the *ex vivo* image stack, for each (*x,y*) lateral coordinate pair, at the axial (*z*) location specified by the curved manifold. **L)** *Top*: Maps of cells obtained from *in vivo* (green, 190 μm below the pia surface, from Area 1 of Fig. 1C) and *ex vivo* (magenta) datasets, showing that different subsets of cells observed *in vivo* are matched with counterparts in several different axial planes in the *ex vivo* image stack (image planes were sampled with an axial spacing of 2 μm; the figure shows planes 10 μm apart). Overlapping pixels from cells present in both maps are shown in white. Scale bar: 200 μm. *Bottom*: Distributions of Soma-print scores for pairs of best-matched (orange) or 2^nd^-best-matched cell pairs (blue), along with parametric fits and semi-logarithmic versions (*insets*) of the plots, shown in the same format as in **G** for each of the image slices in the top row. **M** and **N** show further analyses of the Soma-print results. **M)** Color plot showing the best Soma-print scores for all 592 cells found *in vivo,* when matched using the 2D Soma-print algorithm to the data from individual axial planes in the *ex vivo* image stack. The 3D algorithm identifies for each matched cell the axial plane with the maximum 2D Soma-print score. **N)** Map of 560 cells matched with the 3D Soma-print algorithm out of a total of 592 cells from one *in vivo* imaging plane (95% sensitivity), color-coded by the cells’ registered axial locations. Scale bar: 200 μm. **O)** *Left*: Example *in vivo* two-photon image of pyramidal cells in the mouse motor cortex (Area 1 of Fig. 1C) that densely expressed the green GCaMP8m Ca^2+^ indicator and sparsely expressed the red tdTomato marker. The image shown is a time-average over 3,000 image frames taken at 30 fps, 190 μm beneath the pia surface. *Right*: Reconstructed *ex vivo* image determined using the 3D Soma-print algorithm. White ellipses enclose corresponding sub-regions of the two images. Scale bars: 200 μm. **P)** Example image showing an overlay of paired cells, matched using the 3D Soma-print algorithm, from the *in vivo* (green) and *ex vivo* (magenta) cell maps created for the images of **O**. Overlapping pixels in both maps are shown in white. Scale bar: 200 μm. **Q)** Distributions of Soma-print scores for pairs of best-matched cell pairs (orange) or 2^nd^-best-matched cell pairs (blue), computed for the image of **O**. Data are presented in the same format as in **G**. **R)** We used the sparsely labeled set of red (tdT^+^) fluorescent cells to evaluate the quality of cell pair matching with the 3D Soma-print algorithm for the dataset of **Q**. Using the visually determined matches for this sparse subset of cells as the ground truth, the precision and recall values for matches determined with the algorithm were, respectively, 91% ± 3% and 90 ± 1% (mean ± s.e.m. over n=3 image pairs). See Fig. 1E for definitions of alignment precision and recall.

Therefore, instead of relying on cells’ appearances, we sought a way to assign each cell a distinctive computational signature that would be (i) unique to individual cells, even within large, densely labeled cell populations, (ii) robust to distortions from tissue processing and optical aberrations, (iii) generalizable across a wide variety of imaging modalities used with live animals or for postmortem spatial biology studies, and (iv) readily allow statistical evaluations of cell alignment fidelity. We reasoned that, if we could create such a unique computational fingerprint of individual cells, then these fingerprints could be used to reliably register individual cells across image sets of very different appearances.

Inspired by how humans identify individual stars and constellations in the nighttime sky via their relative locations, we developed a graph-based representation that captures each cell’s geometric relationships to its nearby neighbors. This serves as a unique fingerprint for each cell, which we termed a ‘Soma-print’. The accompanying Soma-print alignment algorithm uses each cell’s Soma-print as the basis for cell registration, reducing the challenge of aligning cells across imaging modalities to that of matching cells with similar vectorial relationships to their neighbors. The algorithm presupposes that, prior to its application, the *in vivo* and *ex vivo* images have been approximately aligned and individual cells have been segmented in each dataset.

We defined a cell’s Soma-print as the set of two-dimensional vectors directed from the cell’s centroid to the centroids of *n* neighboring cells within an individual image plane. This set of *n* vectors, with *n* = 10–15 in typical cases, provides a cell with a geometric signature that is distinct from that of its neighbors (**Figs. 2B** and **S3A**). To register cells, the Soma-print algorithm evaluates candidate cell pairs by assessing the similarity of their vector sets. To do this, the algorithm computes the mean Euclidean distance between pairs of vectors taken from the two cells’ respective Soma-prints and converts this mean distance into a ‘Soma-print similarity score’ (**Fig. 2C**; **Methods**). Finally, this similarity score is adjusted using a multiplicative penalty factor based on the displacement between the two cells in the candidate match. The final scores range from 0–100 (with 100 denoting a perfect match) and have high values for pairs of images of the same cell but low values for images of different cells (**Fig. S3A–C**). Crucially, since it is based on cells’ centroid locations, the similarity score is designed to be robust to differences in image appearance and modest tissue distortions, as the local geometric patterns of cells’ locations should generally be well preserved across datasets.

When implementing an algorithm to match cells based on their Soma-prints, we had to address the issue that *ex vivo* images often contain a greater number of visible cells than the corresponding *in vivo* images. This numeric discrepancy can arise from selective *in vivo* imaging of chosen subclasses of cells, the use of postmortem stains such as DAPI that label all cells, or the difficulty of detecting all cells under the more challenging optical conditions associated with imaging in a live animal. The presence of additional cells in the *ex vivo* data might lead in principle to higher rates of mismatch registrations. To address this issue, the Soma-print algorithm has an iterative design, in which the *n* neighboring cells used to construct each cell’s Soma-print are first selected from a larger pool of *m* candidate neighbors (**Figs. 2D** and **S3D**; **Methods**).

In the initial round of computation, the algorithm first creates a Soma-print for every cell in each of the two datasets by identifying an *m-*fold set of vectors directed from each cell’s centroid to those of *m* of its neighbors. We typically chose *m* to be 15 for *in vivo* and 15–30 for *ex vivo* images, using a greater number for the latter due to the larger number of visible cells postmortem. To evaluate candidate cell matches, in the first round of computation, the algorithm selects *n* pairs of vectors drawn from the larger set of *m* vectors chosen for each cell (**Fig. 2C**). We typically used *n=*10 and selected the *n* vector pairs with the greatest similarity, using the Euclidean distance as a similarity metric. (The final sets of matched cells were insensitive to the value of *n*, as only 0.2% ± 0.05% of cells altered their matches as we varied *n* between 5–20 (**Fig. S3E, F**)). The algorithm converts the mean value of this distance over all *n* vector pairs into a score that is normalized by a factor depending on the two cells’ separation, reflecting the fact that correctly matched cells are more likely to lie nearby in the field-of-view. This yields a Soma-print score for each candidate cell match.

Given the Soma-print scores of all candidate matches, we computed the distributions of scores for all best-matched and all 2^nd^-best-matched cell pairs, as determined across all cells found *in vivo*. Since a Soma-print is a relatively high-dimensional object (*i.e.*, with 20 dimensions in 2D) and each similarity score is based on a sum of similarly distributed distances, we expected the distributions of scores to approximate normal distributions, facilitating statistical evaluations. Specifically, we expected the distributions of 2^nd^-best scores to be approximately Gaussian. Whereas, we reasoned that the distributions of best scores should approximate a sum of two Gaussian functions, one representing the distribution of similarity scores for correctly registered cell pairs and the other representing the scores of misregistered pairs that were incorrectly assigned to the best-matched category.

These intuitions about the distributions’ shapes were borne out in practice, as demonstrated with parametric fits (**Figs. 2E–G**). Bolstering the interpretation that the similarity score distributions for best-matched cell pairs reflected a mixture of correctly and incorrectly registered pairs, one of the two Gaussians in the mixture fit was usually similar in shape to the distribution of similarity scores for 2^nd^-best-matched cell pairs. Overall, these findings led naturally to an empirical Bayes framework^89,90^ for evaluating the quality of cell matching, since, for any best-matched cell pair, the fit parameters allowed us to estimate a p-value, the *a posteriori* probability that the cell pair was correctly registered, and the relative likelihood of an incorrect *vs.* a correct match (**Methods**). We identified the set of matched cell pairs from the first round of computation as those for which the likelihood ratio of an incorrect *vs.* a correct match was <0.05 (**Fig. 2G, H**). We then used these well-matched cells to improve other cells’ Soma-prints (**Figs. 2D**, **S3G,** and **S3H**).

In the algorithm’s second and further rounds, we recomputed the Soma-print for each cell in the *in vivo* map, this time using the *n* nearest cells found in the prior round to have correct matches in the *ex vivo* map. This was based on the idea that the restriction to well-matched cells would yield more trustworthy Soma-prints. Just as humans rely on faithful landmarks to improve recognition of complex spatial scenes, we reasoned that by having our algorithm focus on Soma-prints built from trusted cell matches, it would be better equipped to recognize additional pairs of matched cells. Thus, to identify new matches, for each cell found *in vivo*, we re-computed the Soma-print of each candidate matching cell in the *ex vivo* data using the same *n* well-matched cells comprising the Soma-print of the cell found *in vivo.* Using these Soma-prints based solely on well-matched cells, we then recomputed Soma-print scores across all possible pairs of cells and re-estimated the likelihoods of correct and incorrect cell matches using the distributions of best and 2^nd^-best similarity scores. We retained any cell pairing for which the likelihood ratio of an incorrect match was <0.05. The algorithm stopped when the number of matched cells hardly increased over the prior round, which typically occurred after three rounds (**Fig. S3G, H**). The algorithm then curated a final list of matched cells using a slightly more conservative means of estimating the likelihood of an incorrect match than that used for the intermediate calculations (**Methods**). Runtimes for each round of computation on a conventional (3.7 GHz) desktop computer typically ranged from a few to up to tens of minutes, depending on the total number of cells.

Using the data of **Fig. 1C–E**, in which both image sets were taken with flat focal planes and the red cells served as fiducials, we evaluated cell matches from the Soma-print algorithm against those obtained visually. Despite differences in image appearance, the algorithm yielded large numbers of convincing matches across the *in vivo* and *postmortem* image sets (**Fig. 2F–I**). For example, for the example image pair of **Fig. 2F** with 704 cells detected *in vivo* and 979 postmortem, the algorithm aligned >88% (621/704) of the former, versus 87% (615/704) from manual alignments (**Fig. 2G, H**). For the 6 image pairs from **Fig. 1C–E**, applying the Soma-print algorithm to the images of the densely labeled green cells yielded results very similar to those from human registration (precision: 96 ± 1%, recall: 95 ± 1%, mean ± s.e.m.; **Fig. 2I**). As before, we used the sparsely labeled red cells as fiducials to independently check the registration results obtained with the green cells. Notably, when we used the Soma-print algorithm to align the green cells, the results for the red cells were highly accurate (precision: 99 ± 0.4%, recall: 95 ± 3%, mean ± s.e.m.; **Fig. 2J**).

In many applications of interest, cells from an individual image taken *in vivo* cannot all be localized to the same image plane in a postmortem dataset. This can be due to mechanical misalignment during tissue cutting or field curvature across the *in vivo* imaging field-of-view. To address these scenarios and align cells across multiple planes, we created a three-dimensional (3D) version of the Soma-print algorithm that aligns cells in individual *in vivo* images to those in postmortem, volumetric image stacks (**Fig. 2K–M**). This 3D Soma-print algorithm uses the basic, 2D version of the algorithm described above to match cells across different pairs of planes and then combines this information to determine the best cell alignments in 3D.

Specifically, the 3D Soma-print algorithm first aligns cells observed in the *in vivo* image with those found in the individual image slices of the 3D image stack acquired postmortem. Then, for each cell from the *in vivo* image that was matched to a cell in one or more postmortem image slices, the algorithm identifies the match with the highest Soma-print score (**Fig. 2L–N** and **Fig. S3I, J**). This in turn allows an estimation of the 2D curved manifold within the postmortem image stack that best represents the field-of-view seen *in vivo* (**Fig. 2K**). It is useful to identify this manifold, as its projection to a 2D representation allows direct visual comparisons between the field-of-view observed *in vivo* and a best estimate of how this same field of cells appears in the postmortem data (**Fig. 2O–Q**).

Similar to results obtained with the 2D Soma-print algorithm, the 3D algorithm achieved a high sensitivity, matching 560 of 592 cells found *in vivo* in an initial test dataset. In this particular test, we knew beforehand that the tissue slicing had been slightly misaligned with the *in vivo* imaging plane, so it was heartening that the 3D algorithm could reveal and correct for this cutting error without any special instructions or inputs (**Fig. 2M, N**). For more rigorous benchmarking, we applied the 3D algorithm to *in vivo* and postmortem image sets acquired from the same Drd1a-Cre × Ai14 mice used earlier to assess alignment fidelities by comparing cell matches determined separately for green and red cell pairs. Precision and recall values for cell matches found across the two datasets with the 3D algorithm were 91% ± 3% and 90 ± 1%, respectively (mean ± s.e.m.; n=3 image pairs; **Fig. 2R**).

In sum, the Soma-print algorithm uses the unique geometric relationships between individual cells and their neighbors and is designed to be robust to differences in image appearances and modest levels of tissue distortion. The algorithm has a typical runtime of a handful of minutes and should scale well to large datasets of many different kinds.

### Integrated viral tracing, transcriptomic profiling, and Ca²⁺ imaging

Deciphering how different cell-types contribute to the brain’s circuit dynamics requires knowledge not just of neurons’ transcriptomic classifications but also their connectivity. Thus, we sought to include data about neural projections within the types of information provided by TRU-FACT. To this end, we created a suite of 8 retrograde adeno-associated viruses (retroAAVs), each expressing a unique mRNA barcode, which allows one to label up to 8 neural output pathways from the site of *in vivo* Ca^2+^ imaging (**Fig. 3A**; **Methods**). By injecting one of these retroAAVs in a downstream brain area innervated by axonal projections from neural cell bodies in the Ca^2+^-imaging field-of-view, the cell bodies of neurons with such axons can be monitored *in vivo* and marked via barcode expression. This use of mRNA barcodes to label cells with specific projections follows past viral tracing studies that employed this strategy, albeit without alignment to individual cells’ activity traces^91–96^. By including RNA probes against each barcode in postmortem FISH studies, one can use TRU-FACT to gather combined datasets about functional Ca^2+^ activity, transcriptomic profiles, and axonal projections of the same individual neurons, tracked from *in vivo* imaging sessions to postmortem spatial biology studies. We first designed, synthesized, and packaged a set of 8 distinct barcodes with random sequences, 100–200 bp in length, into retroAAV vectors (**Fig. 3A**). We then sought to verify individual neurons could express multiple barcodes at once. For this test, we co-injected all 8 retroAAVs at a single site in the mouse cortex (n=3 mice with co-injections in the motor cortex). Postmortem multi-round HCR-FISH studies revealed that 88% of 735 labeled neurons located at the center of the injection site expressed ≥ 7 of the barcodes, and 76% of these cells expressed all 8 barcodes (**Fig. 3B,C**). These results illustrate that cortical neurons can express multiple barcodes at once, lending confidence that our retroAAV suite can be used to identify multiple projections from individual neurons. To test whether viral tracing with mRNA barcodes and transcriptomic profiling could be combined within TRU-FACT, we injected 3 barcode-expressing retroAAVs respectively into the mouse primary somatosensory cortex, ventral medial thalamus, and substantia nigra. These three areas receive inputs from the secondary motor cortex (M2), where we tracked Ca^2+^ activity in layer 2/3 and layer 5 pyramidal neurons in the same mice as they were awake (**Fig. 3D**). To track cells from *in vivo* Ca^2+^ imaging to postmortem multi-round HCR-FISH, we optomechanically aligned (**Figs. S1A**) the imaging planes for the two datasets (**Figs. 3E–G, S4A**). After identifying neurons’ spatial profiles in each dataset (**Methods**), we matched cells with the 2D Soma-print algorithm, only accepting matches for which the estimated likelihood ratio of an incorrect registration was <0.05. With this approach, 278 of 353 layer 2/3 cells (79%) and 187 of 262 layer 5 cells (71%) tracked *in vivo* were matched to one of the cells (**Figs. 3H–J** and **S4B–D**) found by HCR-FISH, allowing us to identify axonal projections for neurons for which we had recorded Ca^2+^ activity traces (**Fig. S4E**). Among motor cortical pyramidal neurons, 9% of layer 2/3 cells and 34% of layer 5 cells expressed two or more barcodes, revealing their projections to more than one brain area.

**Figure 3.**
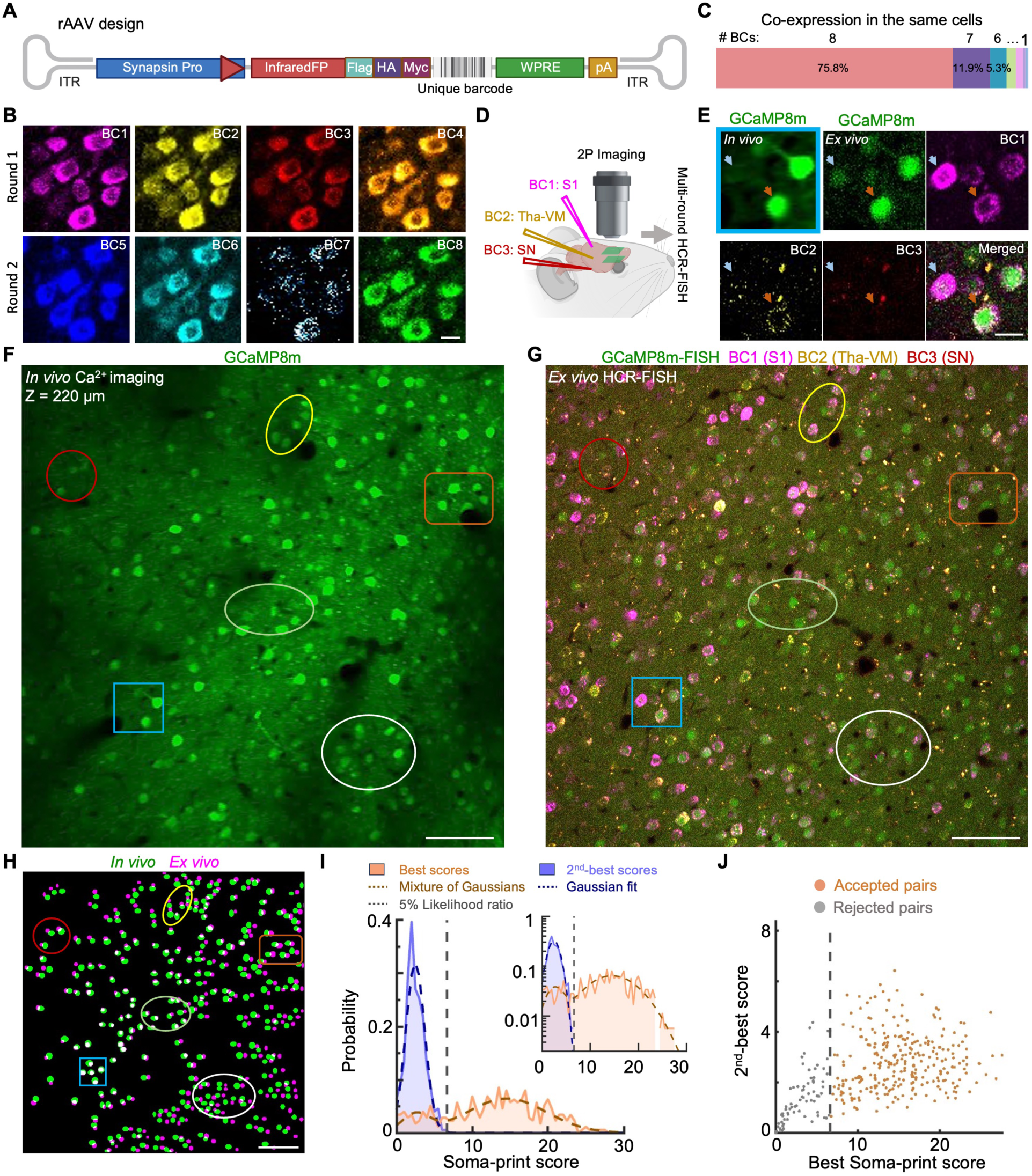
Alignment of neural Ca^2+^ imaging data to RNA FISH images of barcode probes labeling axonal projections. **A)** Schematic of the genetic design of retrograde AAVs used to express RNA barcodes in neurons with axons nearby the site of retrograde virus injection. The DNA vector includes the sequence encoding an infrared fluorescent protein (InfraredFP) fused to a set of 3 epitope tags (Flag, HA, and Myc), which are jointly expressed with the barcode via a synapsin promoter for identifying labeled cells *in vivo* and *ex vivo*, respectively. The barcode sequence was 100 or 200 bp in length (**Methods**) and was followed by the woodchuck hepatitis virus post-transcriptional regulatory element (WPRE) and a polyadenine (pA) sequence. **B)** Example confocal fluorescence images showing mouse primary motor cortical (M1) neurons labeled in either the first (*top row*) or second (*bottom row*) rounds of HCR-FISH imaging using RNA probes for 8 different barcodes (BC1–BC8). We co-injected the mouse with a mixture of the 8 different retroAAVs in area M1 and harvested the tissue 3 weeks later. In addition to imaging DAPI-labeled cell nuclei (405 nm excitation) and an Alexa-488-labeled probe for GCaMP RNA, we imaged Alexa-546 probes for BC1 and BC5, Alexa-594-labeled probes for BC2 and BC6, Alexa-647-labeled probes for BC4 and BC8, and Cy7-labeled probes (750 nm excitation) for BC3 and BC7. Scale bar: 10 μm. **C)** We quantified the percentages of cells that co-expressed different numbers of barcodes, following co-injection of the 8 mixed different viruses at the same tissue site in area M1 (n=3 injections in 3 mice). The percentages (mean ± s.e.m.) of cells co-expressing from 8 to 1 different barcodes were 76 ± 4%, 12 ± 2%, 5 ± 2%, 3.0 ± 1%, 2.3 ± 0.7%, 0.8 ± 0.2%, 0.6 ± 0.4%, and 0.3 ± 0.3%. Results were computed over 208, 327 and 200 labeled cells located at the center of the injections for the three different mice, respectively. **D)** Schematic of an example study in which neurons observed during *in vivo* two-photon Ca^2+^ imaging in the motor cortex (M2) of awake mice were matched to barcode-expressing neurons found by postmortem HCR-FISH. Retrograde AAVs expressing distinct barcodes (BC1–BC3) were injected in somatosensory cortex (S1), ventral medial thalamus (Tha-VM), and substantia nigra (SN). **E)** Magnified views of the image subregion in area M2 enclosed by the cyan square in **F** and **G,** showing results from *in vivo* two-photon Ca^2+^ imaging (*upper left*), and HCR-FISH imaging of RNA probes for GCaMP8m (*upper middle*), BC1–BC3 (*upper right, lower left* and *lower middle*), as well as a merger of all the HCR images (*lower right*). Sites of barcode injection are those in **D**. Red and blue arrows point to two example cells across all 6 images. Scale bar: 20 μm. **F, G)** An example pair of corresponding images, acquired by *in vivo* two-photon Ca^2+^ imaging (**F**, an average of 3,000 frames captured at 30 fps at a depth of 220 μm below the pial surface in cortical area M2, showing cells expressing GCaMP8m), and HCR-FISH (**G**, fluorescence confocal image of labeled barcodes revealing neural projection targets (BC1, area S1; BC2, Tha-VM; BC3, SN red), and Alexa-488-labeled GCaMP RNA (green)). Example subregions are enclosed in colored perimeters to highlight readily visible sets of matched cells. Scale bar: 100 μm. **H)** Overlay of two cell maps for neurons matched across the *in vivo* Ca^2+^ imaging (green) and *ex vivo* HCR-FISH (magenta) datasets. Colored ellipses and rectangles enclose the same cells and subregions as in **F** and **G**. Scale bar: 100 μm. **I)** Distributions of Soma-print similarity scores and corresponding parametric fits for pairs of best-matched and 2^nd^-best-matched cells, plotted in the same format as Fig. 2G for the images of **F** and **G**. Using the likelihood ratio cutoff of 0.05 (vertical dashed line), we successfully matched 278 of 353 cells found *in vivo* to one of the 690 cells found in the *ex vivo* data. **J)** Scatter plot showing the best (*x-*axis) versus the 2^nd^-best Soma-print (*y*-axis) scores, in which each data point denotes the results from an individual neuron. Vertical dashed line divides the cells into two subsets, those for which the likelihood ratio of an incorrect match *vs.* a correct match is greater (gray points) or less than (orange points) 0.05.

These results show TRU-FACT can reveal projectomic and transcriptomic attributes of the same cells monitored with *in vivo* Ca^2+^ imaging. Combined acquisition of the three types of data for individual cells will enable unprecedented, multimodal analyses of neural circuits.

### Alignment of large field-of-view Ca²⁺ imaging and spatial transcriptomic datasets

We next tested whether TRU-FACT allows reliable alignments of cells imaged *in vivo* over a large field-of-view using a two-photon mesoscope. Use of mesoscopes for Ca^2+^ imaging across large cell populations has become increasingly commonplace^97–104^, but the objective lenses used for such studies typically have substantial field curvature and spatially non-uniform resolution across the focal plane, posing challenges for multimodal alignments. We applied TRU-FACT to Ca^2+^ imaging data taken with a two-photon mesoscope^104^ with a field curvature of ∼28 µm axial displacement of the focal plane from the center to the edge of the 4-mm^2^-field-of-view (**Fig. S5A**). This is much greater than the ∼3 μm axial displacement of the focal plane of the conventional objective lens (0.49-mm^2^-field-of-view) we had used earlier for our initial Ca^2+^ imaging studies (**Fig. S2A**).

We monitored *in vivo* Ca^2+^ dynamics from the mouse motor cortex with the mesoscope and then prepared 100-μm-thick tissue sections that fully included the cell population imaged *in vivo* (**Figs. 4A**, **S1B**). In these sections, we used multi-round HCR-FISH to identify mRNA markers of specific neuron-types within 3D confocal image stacks. The 3D Soma-print algorithm matched cells expressing GCaMP *in vivo* with those containing GCaMP mRNA postmortem (**Fig. 4B,C**). Despite the curvature of the *in vivo* imaging field, our optomechanical workflow (**Fig. S1B**) and Soma-print alignments allowed cells to be reliably matched across the two datasets (**Figs. 4D** and **S5B–F**).

**Figure 4.**
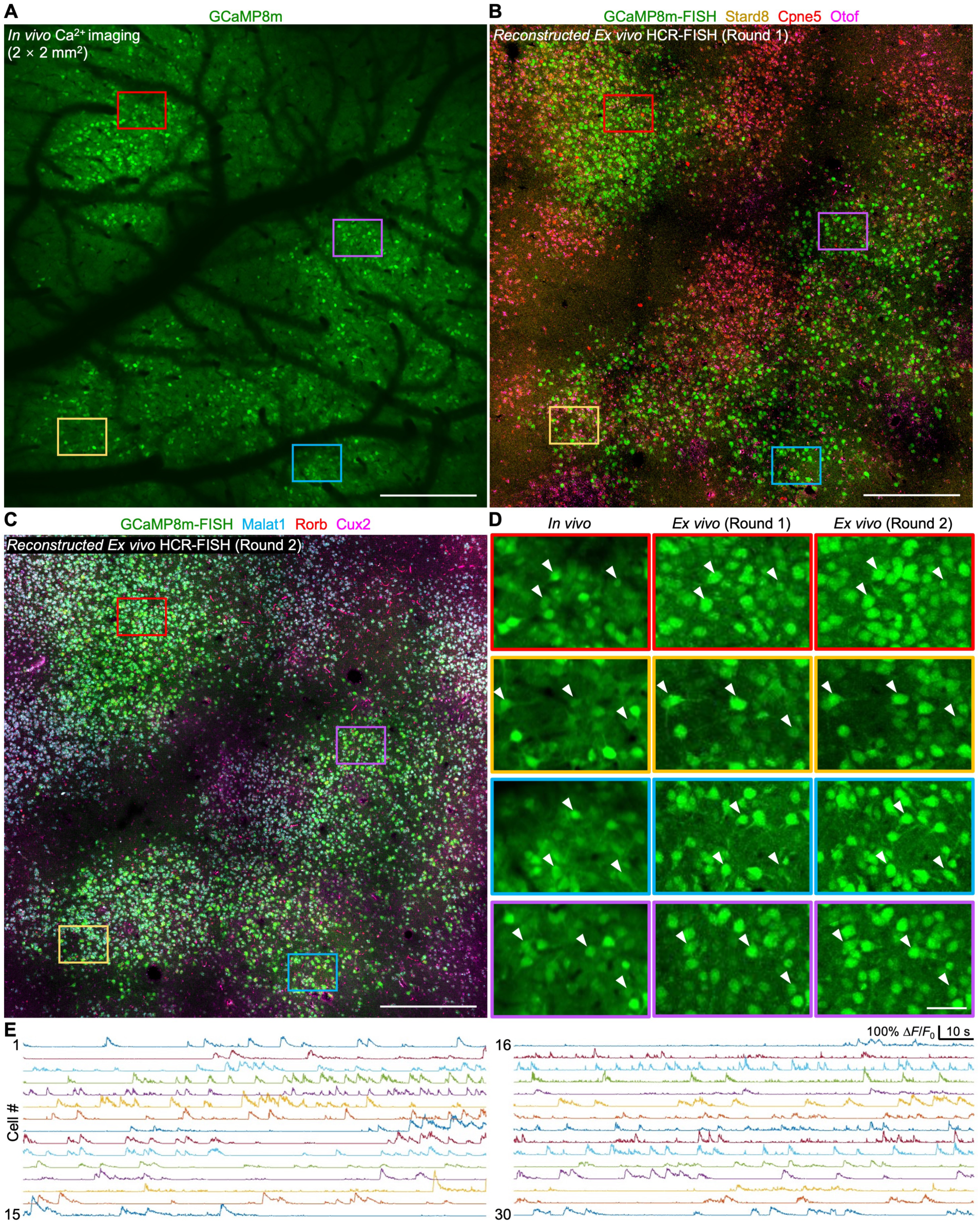
Alignment of Ca^2+^ images acquired in a live mouse with a two-photon mesoscope to HCR-FISH images. **A–C)** Example image, **A**, of GCaMP8m-expressing pyramidal neurons in the motor cortex of an awake resting mouse (mean of 2,000 image frames taken at 7.5 fps, 150 μm beneath pia surface) observed by *in vivo* Ca^2+^ imaging with a two-photon mesoscope^104^. Corresponding confocal fluorescence images (reconstructed by 3D Soma-print) from Round 1 (**B**) and Round 2 (**C**) of HCR-FISH imaging with 100-μm-thick tissue sections (Round 1 (**B**) fluorescent RNA probes were to Otof (magenta), Stard8 (yellow), Cpne5 (red) and GCaMP8m (green); Round 2 (**C**) probes were to Malat1 (blue), Rorb (red), Cux2 (magenta), and GCaMP8m (green)). Colored rectangles highlight regions with sets of cells that are readily matched by eye and shown at greater magnification in **D**. Scale bars: 400 μm. **D)** Magnified views of GCaMP8m expression across the regions enclosed by color-corresponding rectangles in **A–C**, as observed *in vivo* and in both rounds of HCR-FISH. Arrowheads point to individual matched neurons across the different images. Scale bar: 50 μm. **E)** Example traces of Ca^2+^ activity for 30 cells among the 1,615 neurons tracked in the live mouse with the two-photon mesoscope during quiet wakefulness.

Among the 1,615 cells tracked by *in vivo* Ca^2+^ imaging (**Fig. 4E**), the 3D Soma-print algorithm re-identified 1,359 cells among the 41,945 total cells segmented across 33 confocal image slices in the HCR-FISH data, yielding a registration sensitivity of ∼84% (**Fig. S5B–F**). Overall, by illustrating the alignment of large, curved *in vivo* imaging fields to spatial transcriptomic data, these results open the door to diverse studies combining the use of two-photon mesoscopes for *in vivo* imaging and spatial biology methods for in-depth cell-type characterizations.

### Aligning RNA and Ca²⁺ activity datasets acquired in deep brain areas and freely moving mice

Many important applications of *in vivo* Ca²⁺ imaging involve monitoring the activity of deep brain areas with a GRIN microendoscope^62,105–111^. However, GRIN lenses and microendoscopes generally suffer from substantial optical aberrations^62^ and curvature of the imaging field, making it highly challenging to align cells to those in postmortem datasets. For instance, the field curvature of a typical GRIN microlens used for *in vivo* imaging can lead to a ∼100 μm axial displacement of the focal plane across the field-of-view (**Fig. S6A**). A recent seminal study registered hypothalamic neurons observed in two-photon Ca²⁺ videos taken with a microendoscope to multi-round spatial transcriptomic datasets acquired in thin (∼14 μm) tissue sections^62^. However, the alignment approach in this prior study relied on a detailed optical characterization of the aberrations in one particular microendoscope and thus may not generalize to other optical systems and experiments^62^. The study also relied on manual cell alignments and lacked a framework of the type provided here for quantifying the statistical likelihood of correct cell matching.

To provide a fully generalizable alignment approach that can be applied without the need to characterize and account for microendoscope aberrations, we explored the use of TRU-FACT to align neurons across deep brain Ca²⁺ imaging and spatial transcriptomic datasets. We acquired Ca²⁺ videos in the dorsal striatum of mice allowed to walk or run in place by imaging GCaMP-labeled neurons via a GRIN microlens (**Figs. 5A–F** and **S6A–C**) or a microprism (**Fig. S6D–H**).

**Figure 5.**
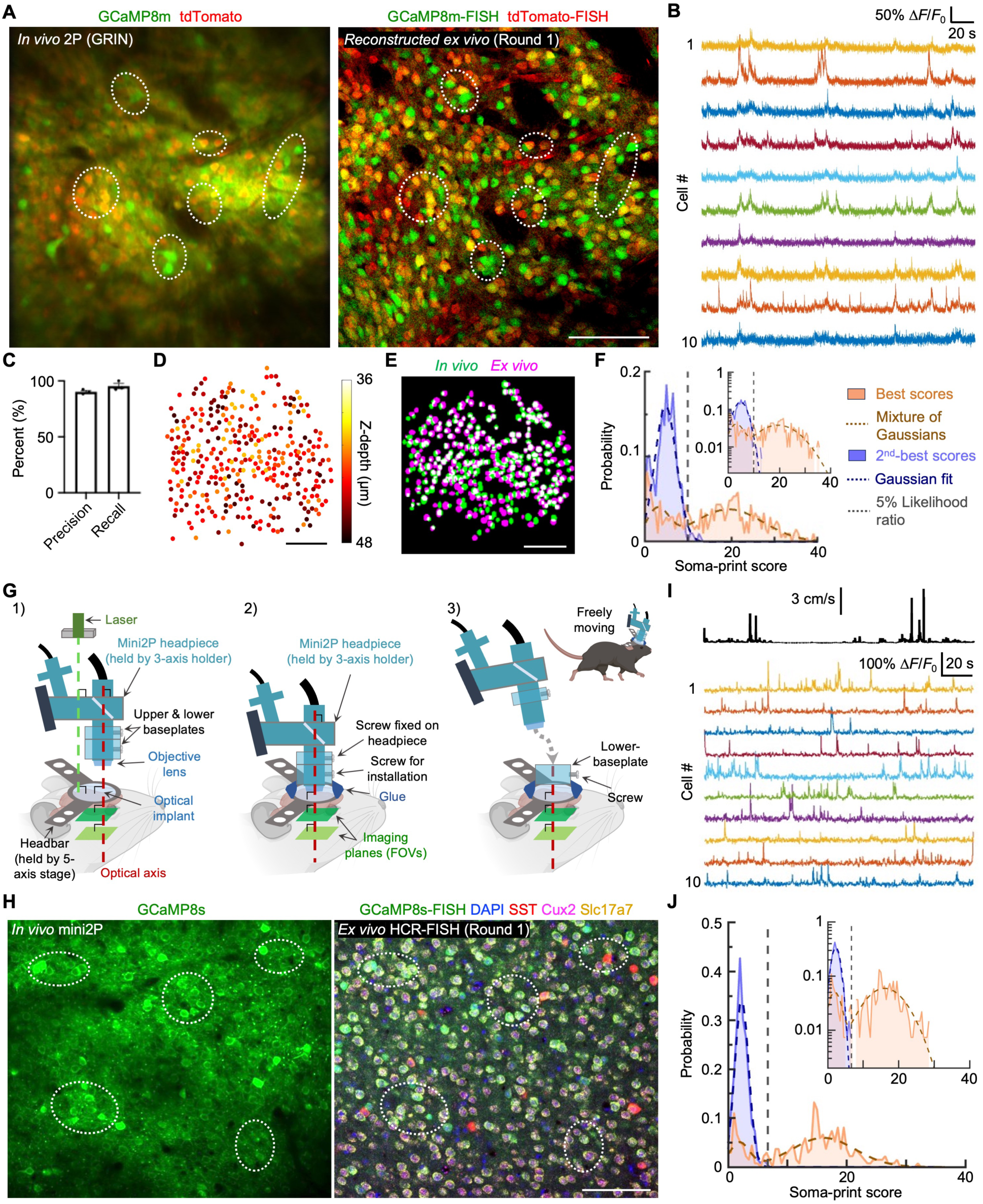
Alignment of Ca^2+^ videos, obtained in deep brain tissue via a GRIN microlens or during naturalistic mouse behavior using a miniscope, to postmortem RNA FISH images. **A)** Corresponding images from the dorsal medial striatum of a Drd1a-Cre ✕ Ai14 mouse, in which spiny projection neurons (SPNs) expressing the D1-dopamine receptor are labeled with red fluorescent tdTomato. Local injection of AAV2/PHP.eB-CaMKII-jGCaMP8m in the striatum allowed viral expression of GCaMP8m and Ca^2+^ imaging in both major classes of SPNs, irrespective of whether they expressed D1- or D2-dopamine receptors. *Left*: Example image taken by two-photon microscopy via a GRIN microlens in a mouse at liberty to run in place on a wheel, averaged over 9,000 image frames acquired at 30 fps. *Right*: Confocal fluorescence image reconstructed by the 3D Soma-print algorithm from the first round of postmortem HCR-FISH images, showing fluorescent labeling of mRNA encoding GCaMP8m and tdTomato. Dashed white ellipses enclose subregions with cells that can be readily tracked by eye across the image panels. Scale bar: 100 μm. **B)** Example traces of Ca^2+^ activity from neurons imaged in **A** via the GRIN lens. **C)** We used the sparsely labeled set of red (tdT^+^) fluorescent cells in **A** to evaluate the quality of cell pair matching with the Soma-print algorithm. Using visually determined matches from the sparsely labeled subset as the ground truth, the precision and recall values for matches determined with the algorithm were 90% ± 1% and 95 ± 2%, respectively (mean ± s.e.m. over n=3 image pairs). (See Fig. 1E for definitions of alignment precision and recall). **D)** Map of 333 cells matched with the 3D Soma-print algorithm out of a total of 414 cells from one *in vivo* imaging plane (80% sensitivity), color-coded by the cells’ registered axial locations in the *ex vivo* z-stack of images. Scale bar: 100 μm. **E)** Image overlay of cell pairs matched by the Soma-print algorithm, **D**, across the *in vivo* (green) and reconstructed *ex vivo* (magenta) cell maps for the data of **A**. Overlapping pixels in the two maps are shown in white. Scale bar: 100 μm. **F)** Distributions of Soma-print scores for pairs of best-matched cell pairs (orange) or 2^nd^-best-matched cell pairs (blue), computed for the image of **E**. Data are shown in the same format as in Fig. 2G. **G)** Schematic of the optomechanical protocol used to align the miniature two-photon microscope (TRANSVISTA) for imaging in a freely behaving animal. **(1)** We first aligned the microscope’s optical axis and the mouse’s implant while the animal was head-fixed. Using a 5-axis mechanical stage and a laser pointer, we used the light reflected off the cranial window to align the mouse relative to the illumination axis of the laser. We then used a 3-axis mechanical holder to co-align the mini-scope and its headpiece to this same axis. **(2)** After identifying a satisfactory field-of-view for two-photon imaging, we glued the lower baseplate of the headpiece to the mouse while it was head-fixed. **(3)** After the glue dried, we detached the microscope from the lower baseplate and returned the mouse to its home cage. Prior to subsequent imaging sessions, we reconnected the upper and lower baseplates to ensure a precise return to the same field-of-view. After *in vivo* imaging was completed, we followed the tissue processing and sectioning steps of **Fig. S1B** or **S1C**. **H)** Corresponding pair of example images of mouse motor cortical layer 2/3 pyramidal cells that virally expressed GCaMP8s. *Left*: Time-averaged image taken during active mouse behavior in an open field enclosure, averaged over 6,000 image frames taken at 9.2 fps (**Supplementary Video 1**). *Right*: Confocal fluorescence image from the first round of HCR-FISH imaging, showing nuclear DAPI staining and fluorescent labeling of mRNAs encoding GCaMP8s, SST, Cux2, and Slc17a7. The image is a sum of the image slices taken of 3 consecutive axial planes spaced 2 μm apart. Dashed white ellipses enclose subregions with neurons that can be readily tracked by eye across the image panels. Scale bar: 100 μm. **I)** *Top:* Example trace of the mouse’s running speed in the open field enclosure. *Bottom*: Ca^2+^ activity traces from 10 example neurons from panel **H**. **J)** Distributions of Soma-print scores for pairs of best-matched (orange) or 2^nd^-best-matched cell pairs (blue), for the data of **H**. Data are shown in the same format as in **F** and Fig. 2G. 211 of the 274 neurons (sensitivity: 77%) found in one plane *in vivo* were successfully matched to a cell found in the *ex vivo* image, each with a relative likelihood of an incorrect match of <0.05.

To evaluate the compatibility of TRU-FACT with these deep brain imaging preparations and to quantify the accuracy of cell registration, we used the same Drd1a-Cre × Ai14 mice as in our prior benchmarking studies performed in neocortex (**Fig. 1C–E**). Striatal spiny projection neurons (SPNs) of both the direct and indirect pathways of the basal ganglia expressed GCaMP8m virally (**Methods**). Only those of the direct pathway expressed the red tdTomato marker, since these SPNs express the D1-dopamine-receptor, whereas those of the indirect pathway do not^107^. As before (**Fig. 1C–E**), the static tdTomato marker was expressed in a subset of cells that served as alignment fiducials for quantifying the registration fidelity achieved separately with the GCaMP images.

During *in vivo* imaging, we recorded SPN Ca²⁺ activity patterns while mice were at liberty to run or walk in place on a wheel (**Fig. 5B**). Afterwards, we dissected each brain, applied the optomechanical protocol of **Fig. S1C**, used multi-round HCR-FISH to probe SPN gene expression, and applied the 3D Soma-print algorithm to align cells across the *in vivo* and postmortem datasets. Using the red cells to provide ground truth alignments against which we could evaluate the cell matches performed separately with the denser set of green cells, the precision and recall values of cell matches output by the 3D Soma-print algorithm for the GRIN lens image data were 90% ± 1% and 95 ± 2%, respectively (mean ± s.e.m.; n=3 image pairs; **Fig. 5C–F**). For the Ca²⁺ videos taken via a microprism, the absence of field curvature implied that the 2D Soma-print algorithm sufficed for cell matching, yielding precision and recall values of 85 ± 1% and 92 ± 2%, respectively (**Fig. S6D–H**).

In addition to imaging deep brain areas, another common usage of micro-optics in neuroscience involves miniaturized microscopes for imaging neural activity in freely behaving animals^109,112–116^. To illustrate TRU-FACT’s compatibility with this application, we used a miniature head-mounted two-photon microscope (TRANSVISTA)^117^ to image the movement-evoked neural Ca²⁺ activity of layer 2/3 motor cortical pyramidal cells during active exploration of an open field enclosure (**Fig. 5G–J**). (**Supplementary Video 1** shows the alignment of a Ca^2+^ video taken in a running mouse to the postmortem HCR-FISH image data, along with other alignments described in the paper). We tracked the mouse’s locomotor behavior with an overhead camera as we imaged the neurons’ Ca^2+^ dynamics (**Fig. 5I**). Among 274 layer 2/3 cells identified *in vivo*, the 2D Soma-print algorithm matched 211 of them to corresponding cells in *ex vivo* HCR-FISH image data (**Fig. 5J**).

Overall, TRU-FACT allows reliable alignments and RNA profiling of cells imaged via a GRIN microlens or a microprism in head-fixed mice, or with a miniature microscope during naturalistic behavior in an open enclosure. This will empower studies of deep brain areas and freely moving animals, enabling multimodal analyses of a range of subcortical regions and behavioral paradigms.

### Alignments involving high-plex spatial transcriptomic techniques

Combining *in vivo* functional imaging with high-plex spatial biology analyses is a key step toward achieving in-depth understanding of how gene expression variations relate to neural dynamics. Both custom and commercial platforms for large-scale, high-plex spatial transcriptomic analyses allow deep molecular profiling, but aligning the individual cells from the resulting datasets to those from *in vivo* Ca²⁺ imaging remains a technical challenge that has not been previously surmounted.

The main obstacles to such alignments stem from the use of thinly cut (∼10 μm) tissue sections, as generally required by commercial spatial biology platforms^13,66,118^. Such thin sections are highly prone to non-rigid deformations that can occur during sectioning and mounting—including wrinkling, folding, tearing, or shearing—making it hard to collect tissue slices in a way that readily permits alignments to images acquired *in vivo*. Further, a tissue slice thickness of ∼10 μm is on the same scale as the optical sectioning provided by the laser-scanning microscopes typically used for *in vivo* imaging. If the cut sections are not accurately aligned parallel to the *in vivo* imaging planes (**Fig. S2B)**, or if the field curvature of the *in vivo* images is comparable to or greater than the thickness of the sections, then the cells observed *in vivo* will be distributed across several or even many postmortem tissue slices. This, in turn, further increases the technical hurdles associated with tissue collection and alignment. Further, the need to analyze multiple tissue slices raises the costs of the high-plex spatial biology assays, which are typically expensive and rise with the number of tissue sections. Finally, cell matching is far more challenging with high-plex spatial transcriptomic data than with low-plex alternatives, due to the typical appearances of the high-plex images, in which individual RNA molecules appear as isolated dots. The resulting images of cells as collections of dots can make it much harder to identify cellular boundaries and tissue features in high-plex images, complicating alignments of two image sets with very different visible appearances and forms of contrast.

To address these challenges, we extended the TRU-FACT optomechanical and computational workflow to thinly cryosectioned tissue slices (**Fig. S1C**). Owing to the accurate, parallel alignment of the thinly cut sections to *in vivo* two-photon images taken with a conventional, flat-field microscope objective lens (**Fig. S2A**), usually only one or two tissue slices contained the entire Ca^2+^-imaging field-of-view (∼700 × 700 μm^2^), easing alignments of cells and their activity patterns to their high-plex molecular profiles. All of our high-plex transcriptomic studies used the Vizgen (MERSCOPE) commercial platform for MERFISH analyses^13^, but the results should be readily extendable to other high-plex platforms that use thin sections, such as the Xenium^66^ or NanoString^119,120^ platforms.

We first verified that 10-μm-thick slices could be sectioned in parallel and matched to the focal planes of conventional *in vivo* two-photon imaging. To do this, we visually aligned ten consecutive 10-μm-thick slices, as imaged with postmortem two-photon microscopy, to their corresponding *in vivo* two-photon imaging planes. We found that every tissue slice was well-matched to a single plane imaged *in vivo*, with easily registered cells distributed across the entire field-of-view (**Fig. 6A**).

**Figure 6.**
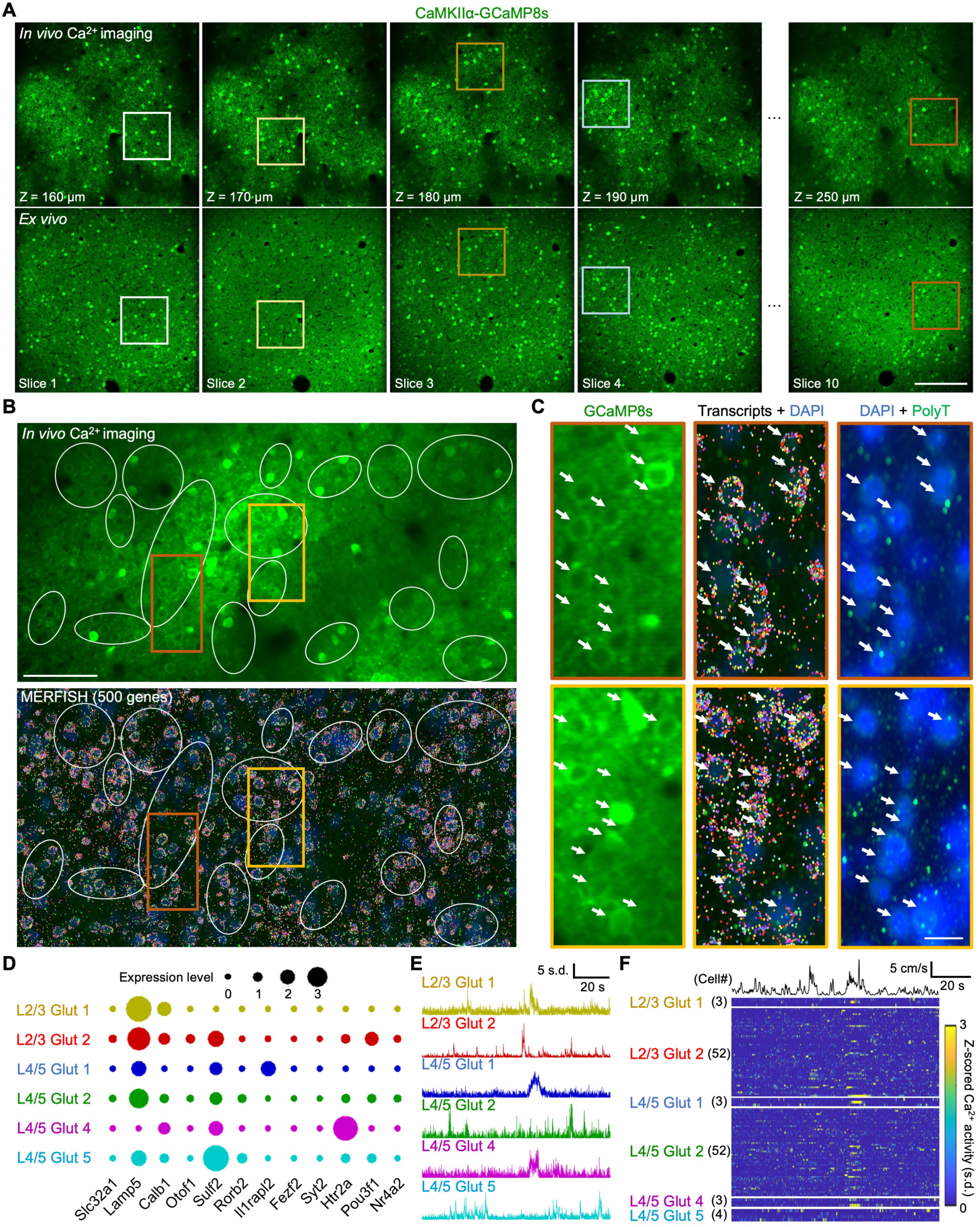
Alignment of *in vivo* neural Ca^2+^ imaging and high-plex RNA transcriptomics (MERFISH) data using a 500-gene panel. **A)** *Top:* Panel of two-photon *in vivo* Ca^2+^ images (averages of 300 frames taken at 30 fps) from mouse motor cortex, showing viral expression of GCaMP8s in pyramidal cells after local injection of AAV2/PHP.eB-CaMKII-jGCaMP8s. The depth below the pia is indicated in the lower left corner of each image. *Bottom,* Corresponding two-photon images acquired postmortem from successive 10-μm-thick tissue sections, using native GCaMP8s fluorescence. Colored squares enclose example areas in which cells can be readily matched by eye across the two sets of images. Scale bar: 200 μm. **B)** *Top*: Example two-photon *in vivo* Ca^2+^ image taken ∼280 μm beneath the surface of motor cortex, showing the viral expression of GCaMP8s in pyramidal cells. *Lower*: Corresponding MERFISH image, taken with a commercial MERFISH instrument (MERSCOPE, Vizgen) and the PanNeuro 500-gene panel. The *Vizualizer* software (Vizgen) assigned each detected RNA molecule a different color in the image, according to which of the 500 genes it encoded. DAPI labeling of cell nuclei is shown in blue. White ellipses enclose regions in which neurons can be readily matched by eye across the two images. Areas within colored rectangles are shown at greater magnification in **C**. Scale bar: 100 μm. **C)** Magnified views of areas enclosed in the red and yellow rectangles in **B**, showing images taken *in vivo* (*left column*), of RNA molecules and DAPI acquired with MERFISH (*middle column*), and of DAPI plus a green fluorescently labeled poly-thymine (PolyT) probe used to reveal DNA (*right column*). Arrows point to cells that are matched across images. Scale bar: 20 μm. **D)** Dot plots showing mean normalized levels of gene expression (**Methods**) for individual transcriptomic clusters of glutamatergic cells determined using a set of 15 cortical marker genes by mapping each cell’s gene expression profile onto the Allen Institute database with MapMyCells (RRID:SCR_024672). Cell clusters are labeled according to the terminology from the Allen Institute. **E)** Ca^2+^ activity traces for 6 example neurons, one from each cluster in **D**. **F)** Color raster plots of Ca^2+^ activity traces from neurons assigned through postmortem MERFISH to each of the glutamatergic neuron-types in **D**, shown beneath a trace of the mouse’s locomotor speed. The total numbers of cells of each type are indicated in the parentheses.

Next, to afford confidence that individual cells would be recognizable and RNA integrity would be well maintained throughout the MERFISH tissue processing and imaging protocol, we performed a standard MERSCOPE validation test using a house-keeping gene, *EEF2,* to examine RNA abundance and image features. The resulting single molecule FISH (smFISH) image showed that RNA integrity was maintained and that tissue features and cells could be well recognized and matched to those in the tissue slice as it was imaged prior to the MERSCOPE protocols (**Fig. S7A**). The aligned datasets showed consistent cell patterns and overall organization, confirming morphological fidelity throughout tissue processing.

Next, we applied TRU-FACT to align high-plex spatial transcriptomic data to two-photon Ca^2+^ videos of layer 2/3 motor cortical neurons in mice allowed to run in place on a wheel (**Fig. 6B, C**). After *in vivo* imaging, we prepared thin tissue slices and used a commercial MERFISH 500-gene PanNeuro panel to profile transcriptomic identities in the same cells monitored *in vivo*. Using the 2D Soma-print algorithm, 196 out of 272 cells found *in vivo* were registered to counterparts in a single 10-μm-thick slice (**Fig. S7B, C**). This allowed us to classify cells based on their transcriptomic markers and to examine how their Ca^2+^ dynamics related to the mouse’s running speed (**Fig 6D–F** and **S7D**).

### Functional activity profiles of layer 5 motor cortical neuron-types during skilled grasping

To study the relationships between neural cell-types, as determined by genetic and projection-based profiling, and their functional activity patterns, we applied the TRU-FACT pipeline to the motor cortex of head-fixed mice trained on a skilled pellet reaching and grasping task. Motor cortical activity is necessary for the performance of this dexterous multi-phase behavior, and prior studies have revealed some functional specializations during this behavior across the region’s diverse neuron classes^21,121–123^. For example, a population of layer 2/3 intratelencephalic (L2/3 IT) neurons appears to encode motor performance outcomes^123^. Further, layer 5 intratelencephalic (L5 IT) neurons appear to signal the movement amplitude, whereas a subset of layer 5 extratelencephalic (L5 ET) neurons projecting to the pons exhibit dynamics relating to movement direction^122^. However, it remains less clear how the activity patterns of finer-grained neural subtypes, as distinguished by gene expression profiles or long-range axonal projections, may differ during execution of this skilled task.

To examine this, we expressed GCaMP8m in neurons of the motor cortical caudal and rostral forelimb areas (CFA and RFA) using an AAV2/PHP.eB-CaMKII-jGCaMP8m virus. We also injected 6 distinct barcode viruses into 6 downstream regions (**Fig. 7A**). As the mice performed the pellet reaching task, we performed two-photon Ca^2+^ imaging of L5 cortical pyramidal neurons in both CFA and RFA at a depth of 550 µm below the pia. After *in vivo* Ca^2+^ imaging, we performed multi-round HCR-FISH on fixed tissue sections aligned to the imaging field-of-view (**Fig. S1B**), which allowed us to profile 12 cortical layer-specific genes and 6 axonal projections (**Fig. 7B, S8A**). Segmentation and matching of cells across the *in vivo* and *ex vivo* datasets allowed us to make direct correspondences between motor cortical cell-types and their recorded Ca^2+^ activity patterns (**Figs. 7C** and **S8B–E**).

**Figure 7.**
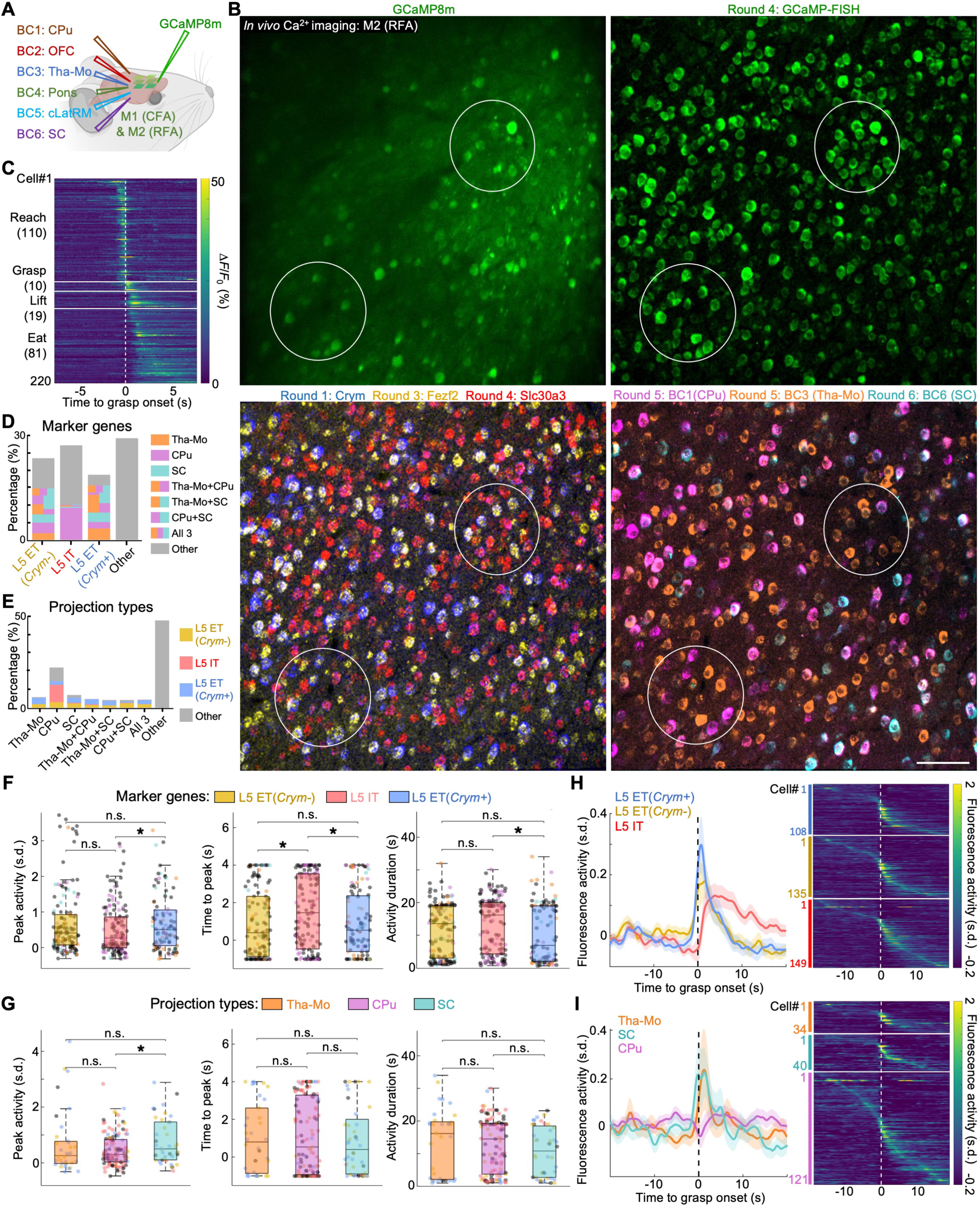
TRU-FACT reveals distinct Ca^2+^ activity patterns during skilled reaching behavior in layer 5 motor cortical neurons with different transcriptomic and projectomic signatures. **A)** Schematic of virus injections used to label motor cortical neurons with various long-range axonal projections. We injected 6 different retroAAVs into C57BL/6J mice to express 6 different barcodes (termed BC1–BC6), respectively, in dorsal striatum (CPu), orbital frontal cortex (OFC), motor thalamus (Tha-Mo), pons, contralateral lateral rostral medulla of the brainstem (cLatRM), and superior colliculus (SC). To virally express GCaMP8m in pyramidal cells in primary and secondary motor cortex (caudal and rostral forelimb areas, respectively), we made a series of local AAV injections in the motor cortex. About 2–3 weeks after virus injections, we trained the mice on the pellet reaching task (**Methods**); Ca^2+^ imaging sessions began 2–3 weeks after that. Example results for a subset of marker genes and axonal projections are shown in **B–H**; additional data from these experiments are shown in **Fig. S8**. **B)** Four co-aligned images. *Upper left*: Mean fluorescence intensity values (average of 9000 image frames acquired at 15 fps) from an *in vivo* two-photon Ca^2+^ video taken in neocortical layer 5 (550 μm depth) of the rostral forelimb area (RFA) of a mouse performing the pellet reaching task. We acquired the other 3 images with confocal microscopy for multi-round, HCR-FISH analyses of the same tissue plane. *Lower left*: Composite HCR-FISH image for 3 marker genes (*Fezf2*, *Slc30a3*, *Crym*) probed in HCR-FISH rounds 1–4. *Upper right*: Fluorescence image of an Alexa-488-labeled HCR-FISH GCaMP probe from HCR-FISH round 4. *Lower right*: Composite HCR-FISH image of the probes to 3 different barcodes that label axonal projections to CPu, Tha-Mo, or SC in HCR-FISH rounds 5 or 6. White ellipses enclose example cells that are readily matched by eye across the images. Scale bar: 100 μm. **C)** Color raster plots showing mean traces of trial-averaged Ca^2+^ activity for 220 example motor cortical pyramidal cells, identified across n=2 mice and temporally aligned to the onset of pellet grasping (white vertical dashed line). Each row shows data for an individual cell, averaged across 5–48 trials per cell. Neural traces are arranged vertically based on the time of their peak, trial-averaged Ca^2+^ activity. 110 of these neurons activated around the time of reach onset, prior to grasping. 10 neurons activated around the onset of grasping. 19 neurons activated around the time that the mouse lifted the pellet. 81 neurons exhibited more prolonged activity across the period when the mouse ate the pellet. **D)** Stacked bar plot showing percentages of motor cortical pyramidal neurons (584 total cells monitored in n=2 mice) with different axonal projection targets, for the following transcriptomic cell-types: Layer 5 intratelencephalic (IT) neurons, *Crym^+^ or Crym^−^* extratelencephalic (ET) neurons, or other neurons that were not assigned to one of these transcriptomic classes. The full height of each bar indicates the percentage of cells among the total neuron population with the designated transcriptomic class. Colored segments within each bar show the percentages of neurons with the indicated patterns of axonal projections. Subsets of cells with axonal projections to two or three target areas are denoted by multi-colored segments. **E)** Stacked bar plot in the same format as that in **D** for the same 584 neurons, showing the different transcriptomic classes of neurons with the designated patterns of axonal projections. **F, G)** Box-and-whisker plots of the (*left*) *z-*scored, peak [Ca^2+^]-related fluorescence activity, (*middle*) the time of peak Ca^2+^ activity relative to grasp onset, and (*right*) the duration of Ca^2+^ activity, quantified as the full width at half maximum (FWHM) of the grasp-aligned Ca^2+^ response, plotted for layer 5 motor cortical neurons of 3 different transcriptomic classes (**F**, intratelencephalic (IT), or *Crym^+^ or Crym^−^* extratelencephalic (ET) neurons) or 3 different projectomic classes (**G**, axonal targets in motor thalamus (Tha-Mo), caudate putamen (CPu), or superior colliculus (SC)). In each plot, the central black line denotes the median, the box encloses the 25th–75th percentiles, and whiskers extend to 1.5 times the interquartile distance. Data points denote values from individual cells. In **F**, colored and gray data points respectively denote cells that did or did not express one of the projection-barcode mRNAs. In **G**, colored and gray data points respectively denote cells that were or were not from one of the transcriptomic types in **F**. We evaluated z-scored [Ca^2+^]-related fluorescence activity parameters across cell classes using a permutation test and 100,000 different random permutations followed by a Dunn-Sidak correction for multiple comparisons (n.s. not significant, \**p*<0.05, \*\**p*<0.01, \*\*\**p*<0.001; the data of **F–I** are from n=108 L5 ET Crym^+^, 135 L5 ET Crym^−^, and 149 L5 IT neurons, and 34 Tha-Mo-, 40 SC-, and 121 CPu-projecting neurons across n=2 mice). **H, I)** *Left*, Mean ± s.e.m. trial-averaged Ca^2+^ activity traces (*z*-scored) relative to the onset of pellet grasping (*t*=0 s), averaged across each transcriptomic, **H**, or projectomic, **I**, class of layer 5 motor cortical neurons noted in **F** and **G**. *Right*: Color plots showing trial-averaged, *z*-scored Ca^2+^ activity levels for individual neurons, plotted relative to the time of grasp onset (*t*=0), with one row per cell, sorted by time-to-peak within each class. 5–48 grasping trials per neuron. Colored numbers indicate the numbers of individual cells in each transcriptomic or projectomic class.

We first categorized L5 neurons by their gene expression and long-range projection patterns. Gene expression analyses identified L5 IT (Slc30a3+) and two subsets of L5 ET neurons (Fezf2+/Crym+, Fezf2+/Crym-) (**Fig. 7D**). Our axonal projection-based analysis focused on three main projection targets, namely motor thalamus (Tha-Mo+), superior colliculus (SC+), and the caudate-putamen (CPu+), as well as combinations thereof (**Fig. 7E**). After combining the results of the genetic and projection-based analyses, we found that the joint classifications fit with prior results; L5 ET neurons projected to all three downstream targets, whereas L5 IT neurons mainly projected to CPu (**Fig. 7D, E**)^124^.

To probe the basic physiological distinctions between these cell populations, we first analyzed the waveforms of all detected Ca^2+^ transient events without reference to the mouse’s behavior. This revealed modest but statistically significant differences among genetically-defined subtypes regarding the decay time constant (τ) and amplitude of Ca^2+^ events (Kruskal-Wallis ANOVA; both p<0.001). Post hoc analyses showed that L5 IT (Slc30a3+) neurons had faster decay kinetics (τ = 1.47 ± 0.02 s; mean ± s.e.m.; n=1,168 events) than either L5 ET (Fezf2+/Crym-) (τ = 1.77 ± 0.02 s; p<0.001; n=1,519 events) or L5 ET (Fezf2+/Crym+) neurons (τ = 2.01 ± 0.03 s; p<0.001; n=1,259 events; Wilcoxon rank-sum tests with Bonferroni correction for multiple comparisons). L5 IT neurons also had Ca^2+^ events with slightly smaller amplitudes (0.63 ± 0.007 Δ*F*/*F*₀ ; mean ± s.e.m.) than either L5 ET (Crym-) (amplitude = 0.68 ± 0.007 Δ*F*/*F*₀ ; p<0.001) or L5 ET (Crym+) neurons (amplitude = 0.66 ± 0.007 Δ*F*/*F*₀ ; p<0.001). Similarly, analyses by axonal projection targets revealed significant differences in both Ca^2+^ event amplitudes and decay time constants across projection-defined classes (Kruskal-Wallis ANOVA, both p<0.001). Neurons projecting to Tha-Mo had Ca^2+^ events with significantly larger amplitudes (0.72 ± 0.01 Δ*F*/*F*₀ ; mean ± s.e.m.; n=519 events) than CPu-projecting (0.63 ± 0.007 Δ*F*/*F*₀ ; n=969 events; p<0.001) or SC-projecting neurons (0.66 ± 0.01 Δ*F*/*F*₀ ; n=396 events; p<0.001; Wilcoxon rank-sum tests with Bonferroni correction for multiple comparisons). SC-projecting neurons also had faster decay kinetics (τ = 1.57 ± 0.04 s; mean ± s.e.m.) than Tha-Mo-projecting (τ = 2.02 ± 0.04 s; p<0.001) or CPu-projecting cells (τ = 1.93 ± 0.03 s; p<0.001).

We next looked at the specific relationships between the Ca^2+^ dynamics of the various subtypes of L5 cells and the pellet reaching behavior^121^ (**Fig. 7C, F–I**). An analysis of Ca^2+^ responses aligned to the grasp phase of the pellet-reaching behavior revealed functionally distinct activity patterns between neural subtypes. We quantified the magnitude of peak Ca^2+^ activity, the time to peak activity relative to grasp onset, and the duration (FWHM) of the grasp-evoked Ca^2+^ activity. Among the genetically defined cell classes, L5 IT (Slc30a3+) neurons exhibited lower peak Ca^2+^ activity than L5 ET (Fezf2+/Crym+) neurons (0.51 ± 0.06 *vs*. 0.78 ± 0.1 z-score; mean ± s.e.m.; permutation test with Dunn-Sidak correction, p=0.03), but were statistically indistinguishable from L5 ET (Fezf2+/Crym-) cells (0.69 ± 0.08 z-score) in this regard. L5 IT (Slc30a3+) neurons also exhibited longer times to their Ca^2+^ activity peak than L5 ET (Fezf2+/Crym-) (1.49 ± 0.16 s *vs*. 0.91 ± 0.16 s; p=0.029) or L5 ET (Fezf2+/Crym+) neurons (0.90 ± 0.17 s; p=0.034). In addition, L5 IT (Slc30a3+) neurons exhibited broader grasp-related Ca^2+^ transients than L5 ET (Fezf2+/Crym+) neurons, as reflected by a longer FWHM (13.19 ± 0.7 s *vs*. 10.47 ± 0.9 s; mean ± s.e.m.; p=0.044), but were not significantly different from L5 ET (Fezf2+/Crym-) neurons in this regard.

Analyses based on cells’ projection targets showed that SC-projecting neurons exhibited peak grasp-aligned Ca^2+^ activity of greater amplitude than that of CPu-projecting neurons (0.77 ± 0.13 *vs*. 0.46 ± 0.05 z-score; mean ± s.e.m.; permutation test with Dunn-Sidak correction, p=0.026), whereas the peak Ca^2+^ activity of Tha-Mo-projecting neurons had an intermediate magnitude that did not differ significantly from the amplitudes of either of the two former groups. In contrast, the time-to-peak and FWHM of grasp-aligned Ca^2+^ activity were statistically indistinguishable across the projection-defined cell classes. Stacked heatmaps of grasp-aligned activity revealed that neurons projecting to the SC exhibited Ca^2+^ activity that peaked around the time of pellet grasping, whereas CPu-projecting neurons exhibited far more heterogeneity in the times of their peak Ca^2+^ responses. Overall, these observations suggest that L5 ET neurons may be more closely involved in motor execution, whereas the more varied timing of L5 IT and CPu-projecting neurons may reflect roles in other facets of the behavior, such as motor planning, feedback-processing, or reward anticipation.

Altogether, these findings demonstrate that genetically and projection-defined L5 neuron subtypes in the motor cortex exhibit distinct functional activity during skilled movement, highlighting the complex interplay between molecular identity, neural connectivity, and animal behavior.

## Discussion

Here we have described TRU-FACT, a generally applicable pipeline for aligning cells across *in vivo* and postmortem image datasets. The pipeline is compatible with thick (**Fig. 1–5, 7**) and thin (**Fig. 6**) tissue sections and intravital two-photon imaging with a mesoscope (**Fig. 4**), miniature microscope (**Fig. 5G–J**), or via a microprism (**Fig. S6D–H**) or GRIN microlens (**Fig. 5A–F**). We demonstrated the robustness of TRU-FACT to tissue distortions or optical aberrations, as well as biological applications using low- or high-plex platforms for spatial transcriptomics together with RNA barcodes for tracing neuronal projections. Illustrating that the pipeline makes cell alignment a routine endeavor, the data here are from 13 mice and 5 different *in vivo* imaging preparations, yielding a total of 10,522 cell matches. The key ingredients of TRU-FACT are an optomechanical approach to aligning the planes of tissue slicing to those imaged *in vivo*, a graph-based signature of individual cells that enables cell alignments to be impervious to differences in the appearances of images taken with different contrast modalities, and a statistical framework that assigns to each cell match a p-value and estimated likelihood that the match was performed correctly.

### Optomechanical workflow to facilitate accurate cell matching

Accurate registrations of individual cells across distinct imaging modalities are essential for integrating functional, anatomical, and molecular information at single-cell resolution. TRU-FACT advances this goal through an optomechanical strategy that aligns and maintains a physical record of the orientation of the *in vivo* imaging planes based on the form of the tissue surface touching the animal’s glass implant. The orientation of these imaging planes is then mechanically transferred postmortem to a parallel set of tissue cutting planes. A simple version of this strategy suitable for beginners involves casting a mold of the tissue form (**Fig. S1A**). More advanced versions are less time-consuming for experienced users and work well with thick or thin tissue sections, using an agarose mold (**Fig. S1B**) or a rigid embedding of the tissue in OCT (**Fig. S1C**), respectively. The use of any of these three versions eases the difficulty of imaging common sets of parallel planes *in vivo* and postmortem, improves the accuracy of cell matching, and makes the task of cell alignment, which used to be considered highly challenging, far more accessible to the average user (see **Methods** for guidance).

A critical but often overlooked physical source of cell alignment errors is the angular mismatch between the *in vivo* and postmortem imaging planes. Even small angular deviations can lead to substantial differences in image content (**Fig. S2B**). These image differences can be further exacerbated by optical aberrations and tissue deformations (**Fig. S2A**). Traditional computational methods for image reconstruction and registration are often ill-suited to handle the unpredictable deformations that can occur during postmortem tissue processing and mounting, especially in the face of optical aberrations that can vary substantially across the *in vivo* imaging field-of-view. We addressed this challenge by developing an optomechanical workflow that maintains angular alignments throughout all stages of *in vivo* and *ex vivo* imaging. By ensuring that postmortem tissue sections are cut and imaged strictly parallel to the original planes of *in vivo* imaging, the workflow reduces the differences between the images taken with different microscopes and contrast mechanisms and reduces the subsequent computational difficulties associated with cell alignment.

### Graph theoretic approach to cell matching

Although our optomechanical alignment strategy eases the difficulty of cell matching, it is not a complete solution to the challenge of matching cells across different imaging modalities. The core issue is that fluorescent labels make cells visible but do not necessarily provide identifiable signatures allowing unambiguous tracking of individual cells. TRU-FACT addresses this weakness by referring to each cell with a unique marker that can be faithfully tracked across datasets acquired with different microscopes and contrast mechanisms. This marker is the Soma-print, which is based not on a cell’s visual appearance but on its geometric relationships to neighboring cells.

The Soma-print is a multi-dimensional object that provides a hallmark of individual cells that is sufficiently distinctive to be robust to moderate levels of tissue distortion, optical aberration, or misalignment during tissue slicing. The Euclidean graph-based representation of the Soma-print is modality-agnostic, generalizable across diverse *in vivo* and *ex vivo* imaging conditions and contrast modalities, and computationally efficient as it only requires cell centroid positions. Empirical distributions of Soma-print similarity scores are well fit with Gaussian functions, facilitating the assignment to each cell match of both a p-value and the likelihood of a correct registration.

Just as human annotators can match cells by referencing nearby cells that have already been matched, the Soma-print algorithm uses a multi-round algorithm to progressively identify matched cells. After an initial round of cell matching, subsequent rounds selectively use cells that were previously matched in prior rounds as the basis for computing a new set of Soma-prints. This iterative design improves matching accuracy and makes the algorithm more immune to being fooled by candidate cell pairs that may partially overlap but nonetheless constitute mismatches.

Overall, the Soma-print algorithm constitutes a robust computational approach for representing and registering cells using distinctive, graph-theoretic signatures of individual cells. We focused here on using the Soma-print algorithm to register cells across *in vivo* and *ex vivo* image datasets, but the algorithm may be more broadly applicable to a range of image registration tasks in biological microscopy and biomedical imaging.

### Statistical framework for cell registration

Unlike many prior studies that relied on human visual scoring and assessments to determine cell matches, we developed a statistical framework to estimate the probability that each individual cell is correctly matched. We reasoned that the distribution of Soma-print similarity scores consists of two qualitatively distinct components, those from correctly and incorrectly registered cell pairs. Accordingly, we approximated the statistical distribution of best-matched scores as a mixture of two Gaussian functions, representing the two components. We used the distribution of Soma-print scores for 2nd-best matches across cells as a means of further characterizing the incorrect pairings. Notably, the distributions of 2nd-best scores were well fit with Gaussian functions and well approximated the portion of the distributions of the best scores that reflected incorrect matches. Through parametric fitting to these Gaussian-distributed components, the Soma-print algorithm evaluates the accuracy of cell registration and yields a p-value for each cell match. This empirical Bayes framework^89,90^, grounded in population-level statistics, provides objective and statistically principled measures of confidence in cell matching results at single-cell resolution, unbiased by human visual assessments.

We verified that the matches determined with our statistical framework closely agreed with those found by human annotators using sparse subsets of red-labeled cells for which visual assessments could be trusted. The automated alignments took minutes to perform, whereas the visual assessments took many hours of painstaking work and were only feasible at all owing to the optomechanical alignment of the imaging and cutting planes. Notably, our statistical framework is flexible, in that users can adjust the likelihood thresholds of accepted matches according to the stringency needed for specific studies. The framework may also generalize to other cell-matching metrics besides the Soma-print score, providing statistical rigor to other means of matching cells.

### Alignment of *in vivo* imaging to high-plex spatial transcriptomics

Combining *in vivo* functional imaging with large-scale spatial transcriptomic profiling is a potent strategy for uncovering how molecular diversity shapes cellular function in intact biological systems. Recent advances allow the detection of hundreds to thousands of genes at cellular resolution in intact tissue slices^13,17,66,125^, offering opportunities to define cells molecularly *in situ* in unprecedented depth.

Despite this potential, most prior efforts to align *in vivo* recorded cells to molecular profiles were limited to small gene panels, typically with <25 genes. Other studies used code-based versions of HCR-FISH or coppaFISH to probe 23- or 72-gene sets, respectively^61,63^. Further, multiple past studies focused on alignments of sparse, inhibitory neuron populations in thick (≥100 μm) tissue sections, due to the greater ease of aligning sparsely labeled cell sets. Alignments of densely labeled excitatory neurons in thin sections (∼10 μm) have remained particularly challenging^54,61^.

Here, we show that TRU-FACT enables the integration of *in vivo* Ca^2+^ imaging with high-plex spatial transcriptomic profiling, using MERFISH and a 500-gene panel for the study of densely labeled excitatory neurons in thin tissue sections. Owing to the precision of our optomechanical alignment procedures, we can re-identify most of the cells observed together *in vivo* within the same plane in one or two of the thin tissue sections. This helps to reduce the substantial costs of high-plex analyses, which typically grow with the number of sections. The compatibility of our pipeline to the use of either thin or thick tissue slices implies that it should readily adapt to additional high-plex spatial transcriptomic platforms as they continue to evolve rapidly.

### Connectivity tracing with RNA barcodes

Mapping neural connectivity and integrating these data with large-scale recordings and molecular profiling at cellular resolution are among the major technical challenges in neuroscience. Traditional tracing techniques are limited by color diversity and co-labeling constraints, which restricts the capability to track multiple projections for individual neurons or brain areas. RNA barcode-based tracing offers a scalable solution that integrates seamlessly with spatial transcriptomics and overcomes these limitations^94,126^. Although the use of RNA barcodes has been conceptually explored and used for standalone studies of tracing, barcode-based tracing has not been previously integrated into investigations of neural function^127–129^. Here, we incorporated barcoded retrograde viral tracers into the TRU-FACT pipeline to map long-range neural projections and integrate this information with functional recordings and transcriptomic profiling. Although our current implementation focuses on retrograde tracing using retroAAVs, our approach can be expanded to more refined connectivity mapping strategies^130,131^, including those for anterograde tracing or using additional viral vectors^132–135^. Ongoing improvements in viral design, tracing efficiency and specificity, and signal amplification will continue to advance multimodal circuit mapping and can be applied with TRU-FACT.

### Transcriptomic and projectomic characterizations of motor cortical activity

To highlight TRU-FACT’s ability to relate cells’ dynamics to their transcriptomic signatures and axonal projection patterns, we used the pipeline to study layer 5 motor cortical neurons in mice performing a skilled reaching task. Prior studies found that motor cortical neurons exhibit functional specialization during skilled movement, with distinct layer- and projection-defined populations encoding features such as movement amplitude, direction, and performance outcome^21,121–123^. In line with this work, we observed systematic differences in task-related activity across both genetically- and projection-defined neural populations.

L5 extratelencephalic *Fezf2*+ neurons had Ca^2+^ activity time-locked to the grasp phase of our motor assay, suggestive of a role in motor execution. L5 intratelencephalic *Slc30a3*+ cells had weaker and temporally broader Ca^2+^ responses, suggestive of roles beyond the immediate production of movements. CPu-projecting cells showed heterogeneous timing of their peak Ca^2+^ activity relative to grasp onset, suggesting this cell population might play a range of roles during the reaching behavior.

These findings illustrate how TRU-FACT allows the neural dynamics underlying the control of movement to be examined in a single experimental framework across cells with multiple distinct molecular markers and projection targets. This capability can be deployed systematically for more extensive characterizations of motor control under normal and pathological movement states, towards identifying cell-types and output pathways that are preferentially affected by movement disorders.

### Limitations of the study

The Soma-print algorithm requires that, prior to its application, the *in vivo* and *ex vivo* image datasets have been approximately aligned and individual cells have been segmented in each dataset. Thus, successful usage of TRU-FACT requires a reasonably good ‘pre-alignment’ of the two image datasets, plus dependable cell segmentation results from both the *in vivo* and postmortem datasets. In scenarios in which the labeling or visualization of cells is insufficient to allow reliable segmentation, cell alignment quality is likely to be impaired. Ongoing advances in cell labeling and visualization methods, pre-alignment and cell segmentation algorithms, microscopy instrumentation, and spatial biology techniques will likely further expand TRU-FACT’s applicability. Although TRU-FACT is designed to be applicable to all cell-types, the experiments presented here focused on the brain and did not include inhibitory neurons or glial cells. Moreover, we have not attempted to apply TRU-FACT with spatial biology techniques requiring the use of fresh frozen tissue. Further work will be needed to validate its performance across a greater diversity of cell-types, tissues, and tissue handling methods.

TRU-FACT enables reliable cell alignments for neural activity data acquired *in vivo* with two-photon microscopy, but its application to neural recordings performed with one-photon fluorescence Ca^2+^ imaging is likely to be more challenging, owing to the lack of optical sectioning. However, instruments that temporally multiplex the acquisition of one- and two-photon Ca^2+^ activity traces might serve as a bridge that allows users to perform one-photon Ca^2+^ imaging in freely moving mice and then later register the cells in the one-photon images to volumetric two-photon image stacks, thereby opening the door to the use of TRU-FACT^136^.

### Outlook

TRU-FACT provides a broadly applicable pipeline for integrating functional imaging with postmortem molecular profiling and long-range connectivity mapping for the same exact sets of cells. The pipeline’s core elements—the optomechanical alignment process, Soma-print cell alignment algorithm, and the statistical framework for evaluating the quality of cell matches—should be readily adaptable to a wide range of molecular profiling methods, including those involving *in situ* RNA sequencing^16,66,137^, spatial proteomics^67,138^, epigenomics^139,140^, or metabolomics^141^.

As new spatial biology platforms emerge, TRU-FACT should be capable of linking the *in vivo* and postmortem datasets that will arise from these new methods. Applying TRU-FACT with high-plex spatial transcriptomic methods that are compatible with thick tissue slices, such as 3D-MERFISH^68^ and STARmap^16^, may increase the total numbers of cells and genes that can be probed with our methods. This in turn may propel studies of activity-dependent gene expression. We also expect TRU-FACT will allow alignments between *in vivo* imaging studies of neural activity and electron- or optical microscopy-based analyses of synaptic connections and the neural connectome^142,143^. Further, the use of TRU-FACT to identify cell-types with specific functional properties should facilitate research and therapeutic efforts to manipulate these cell-types, such as with enhancer virus techniques^144–147^. Due to the versatility of its design and minimal assumptions, TRU-FACT seems poised to impact multiple areas of biomedicine. We anticipate future applications to studies of tissues beyond the brain, such as lymph nodes^148,149^, spinal cord^150^, plants^151,152^, and skin^153,154^, for which information about the dynamics, spatial organization, and molecular states of individual cells is likely to be important for understanding biological and disease mechanisms.

---

**Figure S1.**
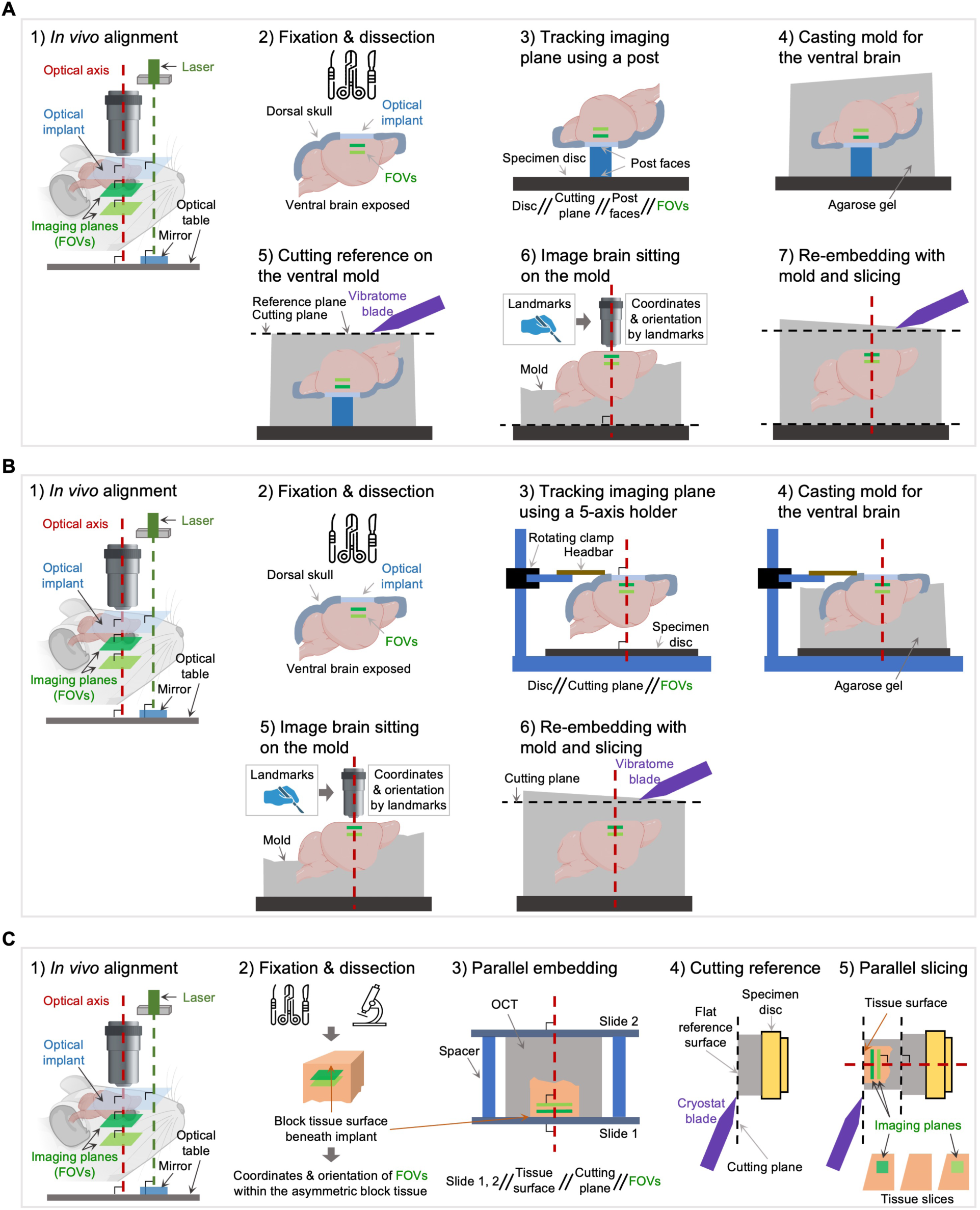
Three approaches to cut tissue slices aligned to *in vivo* imaging planes. We developed three approaches for cutting postmortem tissue sections that were parallel to the tissue planes imaged *in vivo.* Common to all three approaches is a strategy of transferring sets of parallel planes, using the glass surface from the *in vivo* imaging preparation (*i.e.*, the surface of the cranial window, microprism, or GRIN microendoscope). In the first two approaches, we also used a cast mold that captures the shape of the brain as an ancillary tool to facilitate alignment of the planes. The third approach is the one we used for mice imaged with a microendoscope or microprism, although this approach also works well in conjunction with a cranial window preparation. **A)** Alignment using an optical post. **(1)** During *in vivo* imaging, we aligned the axis of optical imaging to be perpendicular to the outward-facing surface of a glass implant, thereby establishing a volumetric set of imaging planes in tissue that all lie parallel to each other at a well-defined angle relative to the optical axis. The schematic illustrates this for the case of a cranial window. **(2)** After *in vivo* imaging, we perfused the animal to fix the brain tissue while the glass implant remained intact. After perfusion, we post-fixed the skull with the glass implant and then removed the ventral portion of the skull, leaving the dorsal part of the skull and the implant intact. **(3)** We glued a custom cylindrical glass post, of which the opposing surfaces were optically flat and mutually parallel, to the surface of the glass implant. We flipped the brain so that its ventral side was facing up, and we glued the other side of the post onto the specimen disc of a vibratome, ensuring that the *in vivo* imaging plane, the two faces of the cylindrical post, the disc and the cutting plane were all mutually parallel. **(4)** We embedded the brain and disc in 2.5% agarose gel. **(5)** Using the vibratome, we cut the gel surface to be parallel to the specimen disc. **(6)** We made a mold of the brain’s ventral surface by cutting the gel approximately in half, roughly parallel to the face of the disc. We detached the mold, brain, remaining parts of the skull, and the implant. We glued the flat surface of the resulting mold onto another specimen disc. We flipped the brain to be dorsal side up, created several landmarks by cutting the brain well away from the site of *in vivo* imaging, and placed the brain back into the mold. We used a two-photon microscope to measure the positions of the landmarks relative to the imaging field-of-view. **(7)** We embedded the brain and initial mold within another 2.5% agarose gel and proceeded to perform vibratome tissue sectioning. The resulting tissue slices were directly used for HCR-FISH studies (sections of 100–300-μm thickness) or further processed for MERFISH studies using the method of (**C**) to prepare 10-μm-thick slices. **B) Alignment by direct embedding of a whole post-fixed brain**. **(1, 2)** The first two steps were identical to those of the first approach, as described in (**A**). **(3)** We used a custom mechanical holder with 5 degrees of mechanical freedom to grasp the fixed mouse brain with its glass implant still intact. We placed the brain above a vibratome specimen disc, leaving some air space between the two. Using the 5 degrees of mechanical freedom, we aligned the optical axis to be perpendicular to the outward-facing surface of the glass implant, matching the orientation used during *in vivo* imaging. **(4)** We embedded the brain with its implant intact, together with the vibratome specimen disc, in a 2.5% agarose gel. **(5, 6)** We made a mold of the brain, as in step 6 of panel **A**, and cut tissue sections as in step 7 of panel **A**. Theoretically, we can use this approach to track *in vivo* planes that are at any angle and transfer them to be parallel with the tissue sectioning and *ex vivo* imaging. Additionally, we can also cut from the ventral brain side in step 6 after cutting the dorsal mold surface as a reference plane if the imaging area is deep in the brain. **C) Alignment using direct embedding of a brain tissue block**. The third and most general strategy is schematized here and is the same as that described in Fig. 1B.

**Figure S2.**
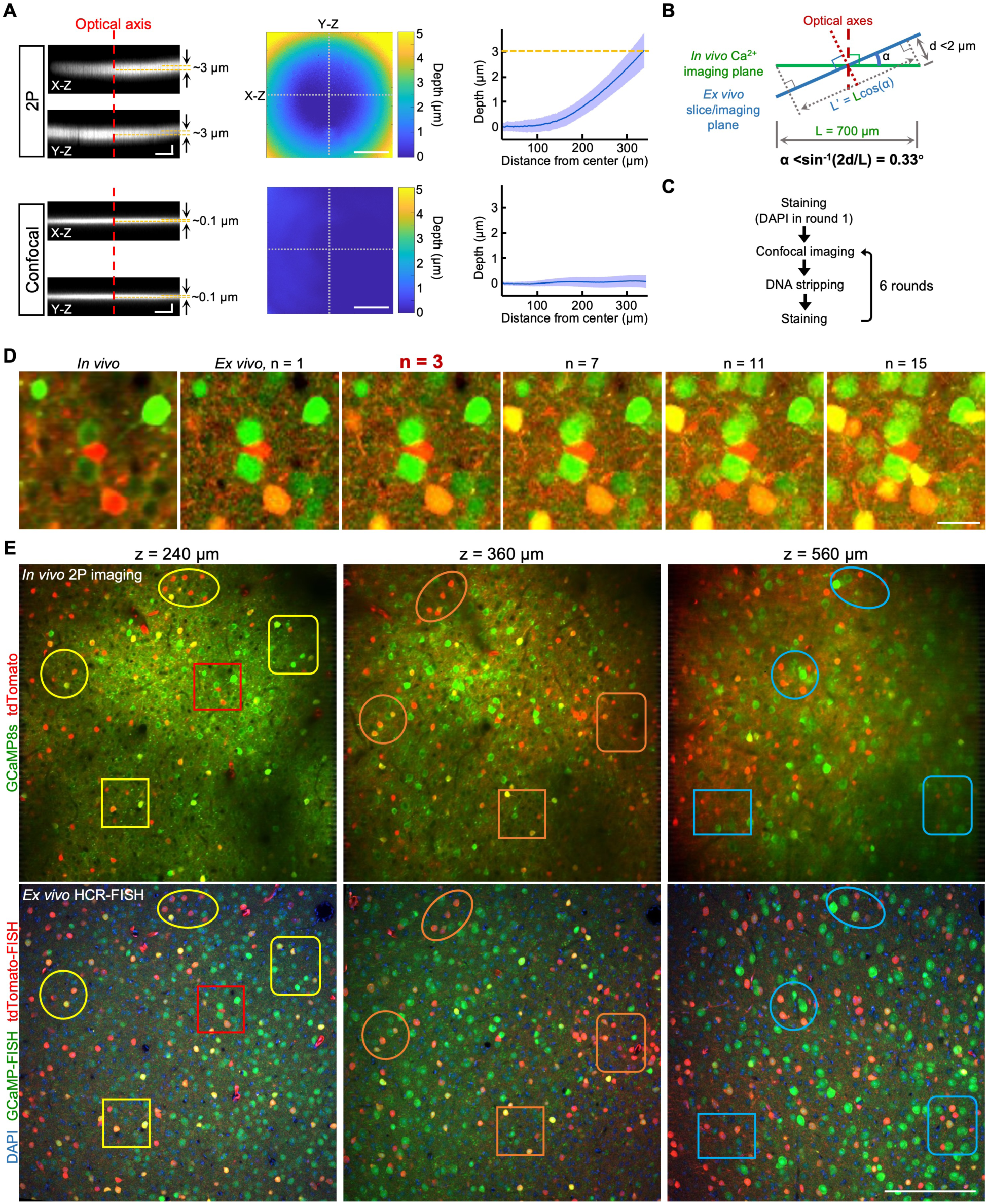
The focal manifold of the objective lens used for two-photon Ca^2+^ imaging was sufficiently flat for precise, volumetric alignments between the optical planes imaged *in vivo* and flat tissue sections imaged by confocal fluorescence microscopy. **A)** We evaluated the field curvatures of the objective lenses used for *in vivo* two-photon microscopy and confocal fluorescence imaging of the tissue sections cut for HCR-FISH. The data shown were obtained for a custom-built two-photon microscope (*upper row*) equipped with a 16× water immersion Nikon CFI75 LWD 0.8 NA objective lens with a 3.0-mm-working distance, and a Leica SP8 confocal fluorescence microscope (*bottom row*) equipped with a Leica HC Plan APO CS2 20× / 0.75 IMM objective lens (**Methods**). As expected, the plan apo objective lens had minimal field curvature. The objective lens used for two-photon microscopy had a level of field curvature that was measurable but still sufficiently minimal to allow precise alignments of the *in vivo* and *ex vivo* images. *Left*: Cross-sectional projections from volumetric image stacks acquired by using either the two-photon or the confocal microscope to image a fluorescent glass coverslip. With the confocal microscope, the image of the fluorescent glass surface exhibited ∼0.1 μm axial deviation from the center of image to the periphery. By comparison, using the two-photon microscope, the image of the fluorescent glass surface exhibited ∼3 μm axial deviation from the image center to the periphery. Scale bars: 100 μm (X and Y lateral dimensions) and 10 μm (Z, axial dimension). *Middle*: Color plots showing the axial displacement of the image of the slide as a function of the lateral displacement from the optical axis. For each (X, Y) position, we determined the axial displacement of the image of the slide by performing a Gaussian fit to the image brightness values as a function of Z and then taking the Z position corresponding to the peak of the Gaussian. Scale bar: 200 μm; *Right*: Mean ± s.d. axial displacement of the two-photon and confocal images of the fluorescent slide, plotted as a function of the radial displacement from the optical axis, averaged over all azimuthal angles. **B)** Diagram showing that for an imaging field of 700 μm width, the polar angular misalignment between the cut tissue sections (which should be parallel to the *ex vivo* imaging plane) and the *in vivo* imaging field should ideally be <0.33° to ensure an axial deviation of <2 μm at the edge of the field-of-view, which is finer than the axial FWHM (∼4.8 μm) of the point spread function of the two-photon microscope. **C)** Summary of the six-round HCR-FISH pipeline. We fluorescently labeled tissue slices using a multi-round HCR-FISH protocol^155^ (including a DAPI stain of nuclear DNA in round 1), imaged the slices on a confocal fluorescence microscope, and then stripped all single-stranded DNA probes from their mRNA targets before the next round of labeling and imaging. Owing to the DNA stripping step, the DAPI stain of nuclear DNA was only visible in round 1. To provide fiducial markers allowing the alignment of cells across rounds, in all rounds of HCR-FISH we used Alexa-488-labeled probes of GCaMP mRNA that we imaged in the green fluorescence channel. **D)** Panel of images showing how the visual similarity of an *in vivo* two-photon image (*leftmost image*) to postmortem confocal fluorescence images of the corresponding tissue section processed with HCR-FISH depended on the number of confocal image planes that were averaged together. The five *ex vivo* confocal images shown are maximum projections of n = 1, 3, 7, 11 or 15 image slices (2 μm axial spacing between successive slices; 100-μm-thick tissue section). Throughout this paper, we used projections of n = 3 confocal image slices to boost the resemblance of corresponding *in vivo* and *ex vivo* images, as this choice for how to process the confocal data generally yielded a good compromise between ensuring the inclusion of cells observed *in vivo* and avoiding the addition of extra cells that were from outside the *in vivo* image plane. The images shown provide magnified views of the region enclosed in the red square in the upper left image of **E**, which corresponds to Area 2 of and has the same fluorescence labels (GCaMP8s and tdTomato) as Fig. 1C. Scale bar: 20 μm. **E)** Example pairs of corresponding images taken by *in vivo* two-photon microscopy in live mice (*upper row*) or postmortem HCR-FISH and confocal fluorescence microscopy (*bottom row*), across different depths (240 μm, 360 μm, or 560 μm) beneath the cortical surface of Area 2 shown in Fig. 1C, using the same fluorescence labeling strategy with GCaMP8s and tdTomato. The two-photon images shown are time-averaged over 3,000 image frames taken at 30 fps. The *in vivo* Z-stack images were taken from the pial surface (Z = 0 μm) to layer 5 (Z = 600 μm). The confocal images shown are projections over n = 3 image slices (2 μm spacing between slices; 100-μm-thick tissue sections). The colored ellipses and rectangles enclose cells at each tissue depth that are readily matched by eye across the image pairs. Scale bar: 200 μm.

**Figure S3.**
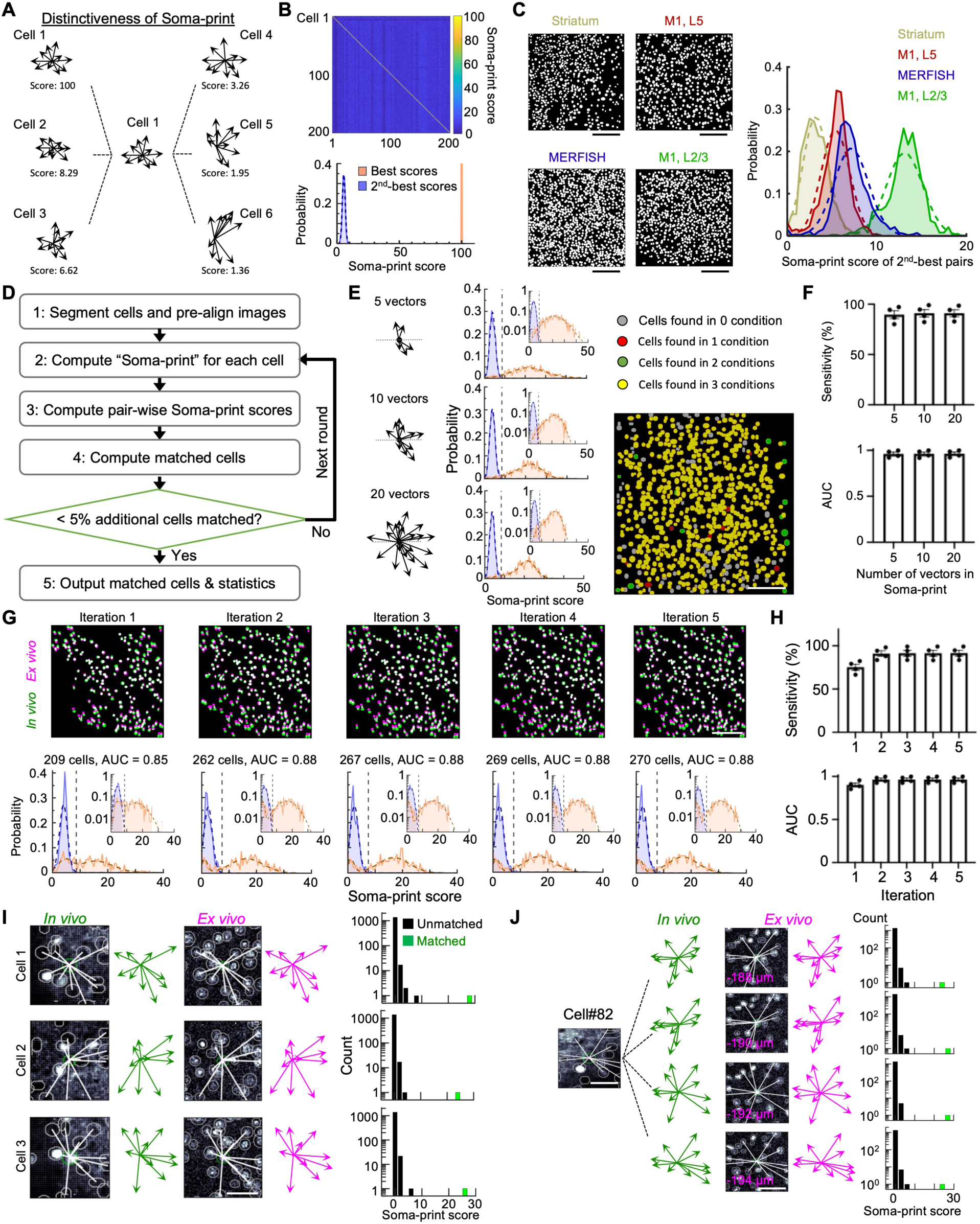
Soma-print specificity and optimization of the Soma-print algorithm. **A)** To illustrate how the Soma-print provides a distinctive marker of individual cells, we computed the Soma-print similarity scores (ranging in principle from 0–100) for 6 different pairs of motor cortical neurons, each comprising Cell 1 and one of Cells 1–6. Shown for each of these cells is its Soma-print, comprising a set of 10 vectors, with each vector extending from the cell body of the cell in question to the cell body of a nearby neuron. The Soma-print similarity score for Cell 1 and itself is 100 (*upper left*), whereas the Soma-print scores for Cell 1 and any of the 5 other cells shown is < 10. **B)** To formalize the observations of **A**, we performed a control analysis in which we computed Soma-print scores for pairs of cells from the same exact image (one of the images used in the analysis for Fig. 1E). *Upper*: Matrix of Soma-print similarity scores computed for each cell (numbered 1–200) on the *x*-axis to every other cell (*y-*axis). *Lower*: Distributions of Soma-print similarity scores, for the sets of best-matched (orange) and 2^nd^-best-matched (blue) cell pairs, based on the data in the upper panel. By construction, each cell’s best-match was to itself, yielding Soma-print scores of 100. Scores for pairs of different cells were generally all < 10, with a median value of ∼6. **C)** We explored how the distributions of 2^nd^-best Soma-print scores (computed with Soma-prints of 10 vectors each) varied across images in which the cells had different densities and spatial patterning. *Left*: Four example cell maps, as determined *ex vivo* through MERFISH studies of layer 2/3 of mouse motor cortex or through HCR-FISH studies of mouse dorsal medial striatum, layer 5 of the primary motor cortex (M1), or layer 2/3 of M1. Scale bars: 200 μm. *Right*: After computing Soma-print similarity scores across pairs of cells within each of the 4 images shown in the left panel, we plotted the distributions of 2^nd^-best similarity scores. Dashed lines show Gaussian fits to each of the distributions. These 2^nd^-best scores tended to be highest for the data from motor cortex layer 2/3 but were generally well below the range of score values for best-matched cell pairs (see panels **E, G**). **D)** Flowchart showing the steps of the iterative Soma-print algorithm. Step 1: *In vivo* and *ex vivo* images of the same field-of-view are pre-aligned to share a common scale and orientation and to generate cell maps. Step 2: The Soma-print of each cell is computed as in Fig. 2B. Step 3: Pairwise similarity scores are calculated as in Fig. 2C. Step 4: Pairs of matched cells are determined based on the set of similarity scores. Step 5: If the increase in the percentage of cells from the *in vivo* dataset that were successfully matched is ≥ 5%, then algorithm goes back to Step 2 for another round of computation. If this percentage is < 5%, then, in Step 6, the algorithm outputs the final set of matched cells, parametric fits, and statistics (*e.g.*, see Fig. 2G and **Fig. S4D**). Notably, in the first round of computation, the method identifies pairs of matched cells across *in vivo* and *ex vivo* images by requiring the pairs to have a likelihood ratio for an incorrect *vs*. a correct match of <0.05. In subsequent rounds, the algorithm re-computes each cell’s Soma-print by restricting the selection of neighboring cells to only those that were successfully matched in prior computational rounds. **E)** We explored whether cell pair assignments varied when each cell’s Soma-print was defined using different numbers of nearest neighbors. *Left*: Soma-prints for an example, individual motor cortical neurons from Fig. 1E, using either 5, 10, or 20 nearest neighbors. *Middle*: Distributions of Soma-print similarity scores for best-matched (orange) and 2^nd^-best-matched (blue) pairs of cells, along with corresponding parametric fits, plotted in the same format as Fig. 2G, using *n* = 5, 10, or 20. *Right*: A cell map based on one of the *in vivo* images from Fig. 1E, in which the color of each cell denotes whether it was successfully matched to a counterpart in the *ex vivo* HCR-FISH irrespective of the value of *n* (yellow cells), for two of the three *n*-values (green cells), for only one value of *n* (red cells), or for none of the three *n*-values (blue cells). The vast majority of cells are colored yellow, indicating that across the range *n* = 5–20, the number of vectors in the Soma-print is not a strong determinant of whether a cell is matched or not. Notably, only 0.2% ± 0.05% (mean ± s.e.m.; n = 3 images from Fig. 1E) of cells altered their matches as we varied *n* values. Scale bar: 200 μm. **F)** For the datasets of Fig. 1E, we plotted the mean ± s.e.m. sensitivity (*top*, the % of cells observed *in vivo* that were also found *ex vivo*) and the area under the ROC Curve (*lower*, the AUC, characterizing the distinguishability of the distributions of Soma-print scores for best-matched and 2nd-best-matched cell pairs), as a function of the number of vectors (*n =* 5, 10 or 20) used in the Soma-print (n=4 images). The results were largely indistinguishable, irrespective of whether we used Soma-prints with 5, 10, or 20 vectors. Black data points indicate results from individual images. **G)** We examined the outputs of the Soma-print algorithm across its iterative rounds of computations using a different image from the data of Fig. 1E than that used in panel **E** above. *Upper*: Overlaid maps of cells found *in vivo* (green) and *ex vivo* (magenta) after each of the first 5 rounds of computation. Scale bar: 200 μm. *Lower*: Distributions of Soma-print scores for best-matched (orange) and 2nd-best-matched cell pairs (blue) at the end of each iteration, with parametric fits to the distributions shown in dashed lines, plotted in the format of Fig. 2G. Above each panel are the number of cells matched across the two imaging modalities and the AUC value characterizing the distinguishability of the two distributions. **H)** Plots of the mean ± s.e.m. sensitivity (*upper*) and AUC values (*lower*, defined as in **F**) across the 5 iterations studied in **G** (n=4 images from Fig. 1E). Both the sensitivity and AUC values plateau around the third iteration. **I)** 3 example matched cells from 3 different axial planes, using the 3D Soma-print algorithm, showing that the local Soma-prints of individual cells from the same *in vivo* image were each well preserved across individual *ex vivo* images, despite a global misalignment that occurred between the *in vivo* image and the *ex vivo* images. Corresponding *in vivo* and *ex vivo* images of the cells, matched Soma-print vectors, and histograms (cell count) of all Soma-print scores are shown for each cell. The cell match yielding the highest Soma-print score is denoted in the histogram with a green bar. Scale bar: 40 μm. **J)** Using the 3D Soma-print algorithm, individual cells observed *in vivo* could generally be matched across multiple different image slices of the *ex vivo* confocal image stack. *Left*: Time-averaged projection image of an example motor cortical neuron (cell #82) from Fig. 2L, overlaid with (white) and adjacent to (green) depictions of the Soma-prints that yielded the best matches to images of the same cell in different slices of the corresponding confocal image stack. *Middle*: Images of the same cell in 4 different slices of the confocal stack, acquired with 2 μm axial spacing, overlaid with (white) and adjacent to (magenta) depictions of the Soma-prints that yielded the best matches to the cell’s *in vivo* image. Scale bar: 40 μm. *Right*: Histograms of Soma-print scores, with the *y*-axis shown on a logarithmic scale, for the best (green) and rejected (black) matches between the *in vivo* image of the neuron and candidate cell matches in each of the 4 confocal image slices.

**Figure S4.**
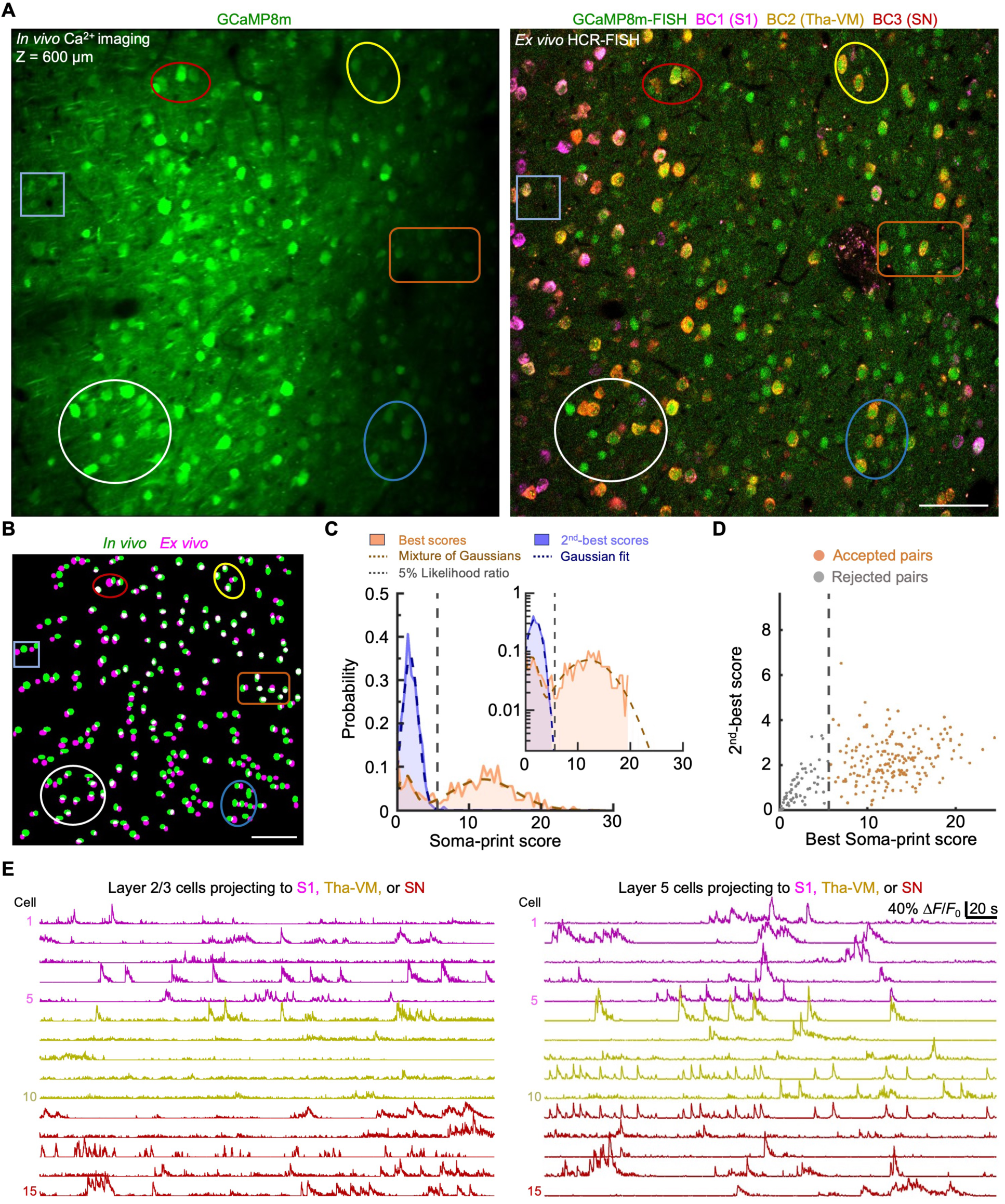
Alignment of Ca^2+^ imaging and RNA barcode data for layer 5 cortical neurons. **A)** Corresponding pair of *in vivo* (*left*; mean image across 3,000 frames taken at 30 fps) and HCR-FISH (*right*) images acquired from tissue 600 μm beneath the surface of motor cortical area M2, from the same mouse as in Fig. 3F, G. The *in vivo* image shows pyramidal cells virally expressing the GCaMP8m Ca^2+^-indicator. The HCR-FISH image shows fluorescently labeled RNA barcodes (BCs) that mark cells with axonal projections to primary somatosensory cortex (S1, BC1), motor thalamus (ventral medial thalamus, Tha-VM, BC2), and substantia nigra (SN, BC3), in addition to a probe targeting GCaMP mRNA (green). Colored ellipses and rectangles in **A** and **B** enclose subsets of cells that are readily matched by eye. Scale bar: 100 μm. **B)** Overlay (white) of the cell maps determined from the *in vivo* (green) and HCR-FISH (magenta) images in **A**. Scale bar: 100 μm. **C)** Distributions of Soma-print similarity scores for pairs of best-matched cell pairs (orange) or 2^nd^-best-matched cell pairs (blue), computed for the images of **A** and shown in the same format as Fig. 2G and Fig. 3I. Parametric fits (dashed curves) used a Gaussian function to fit the distribution of scores for 2^nd^-best matches or a mixture of two Gaussians to fit the distribution of scores for best-matched cell pairs. Using these fits, we computed the likelihood ratio that a given Soma-print score reflects a 2^nd^-best match *vs.* a best-match. In practice, we only accepted matches for which this likelihood ratio was <0.05 (vertical dashed line marks this 0.05 cutoff value). *Inset*: The same distributions and fits shown with a logarithmic scale on the *y-*axis. Overall, 187 of the 262 cells seen *in vivo* in layer 5 were successfully matched to one of the 456 cells found *ex vivo*. **D)** Scatter plot plotted in the same format as Fig. 3J, with each data point showing results for an individual cell in **A, B**. Each point’s *x-*coordinate value is the Soma-print similarity score for the cell’s best-matched pairing; the *y*-coordinate is the Soma-print score for the cell’s 2^nd^-best pairing. Vertical dashed line divides the cells into two subsets, those for which the likelihood ratio of an incorrect match *vs.* a correct match is greater (gray points) or less than (orange points) 0.05. **E)** Example Ca^2+^ activity traces for pyramidal cells of layer 2/3 (*left column*, cells from Fig. 3F) and layer 5 (*right column*, cells from **A**). The colors of the individual traces indicate neurons that have axonal projections in areas S1, Tha-VM, or SN.

**Figure S5.**
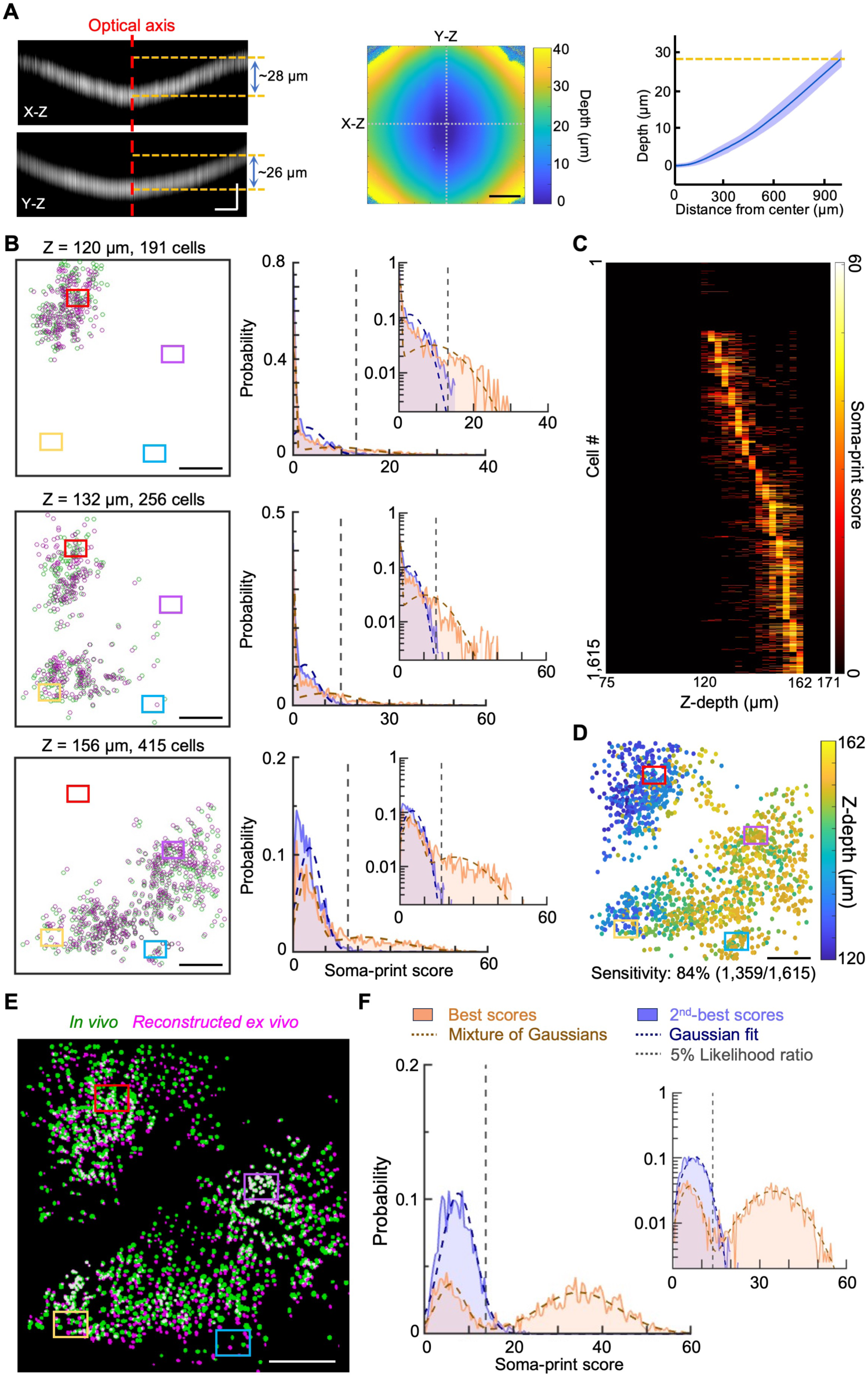
Field curvature for mesoscopic two-photon imaging and alignments to HCR-FISH confocal image stacks using the 3D Soma-print algorithm. **A)** We evaluated the field curvature of the objective lens^97^ used with our *in vivo* two-photon mesoscope^104^. Owing to the substantial curvature, alignments of cells observed *in vivo* to those found in postmortem confocal image stacks spanned across multiple slices of the confocal image stack. *Left*: Cross-sectional projections from volumetric image stacks acquired using the two-photon mesoscope to image a fluorescent glass coverslip. The image of the glass surface exhibited 26–28 μm axial deviation from the image center to the periphery. Scale bars: 200 μm (X and Y lateral dimensions) and 20 μm (Z, axial dimension). *Middle*: Color plots showing the axial displacement of the image of the slide as a function of the lateral displacement from the optical axis. For each (X, Y) position, we determined the axial displacement of the image of the slide by performing a Gaussian fit to the image brightness values as a function of Z and then taking the Z position corresponding to the peak of the Gaussian. Scale bar: 400 μm; *Right*: Mean ± s.d. axial displacement of the two-photon mesoscope image of the fluorescent slide, plotted as a function of the radial displacement from the optical axis, averaged over all azimuthal angles. **B)** Automated cell alignment using the 3D Soma-print algorithm to register cells from the *in vivo* mesoscope image of Fig. 4A to multiple image slices of the *ex vivo* confocal image stack. *Left:* Overlaid maps of matched pairs of cells found *in vivo* (green) and *ex vivo* (magenta) across 3 example image planes with 191, 256, and 415 matched cells, respectively. *Right*: Corresponding distributions of Soma-print similarity scores and parametric fits, plotted in the same format as Fig. 2G, for the cell pairs matched in each panel at left. Colored rectangles in **B, D** and **E** enclose the same cells and subregions as those in Fig. 4A**–C**. Scale bars: 400 μm. **C)** Color plot showing the best Soma-print scores, denoted in color, for all 1,615 *in vivo* cells for each of 33 axial planes in the corresponding confocal image stack (axial spacing of 3 μm, *i.e.,* finer than the ∼6 μm axial extent of the two-photon mesoscope’s point-spread function). Each cell observed *in vivo was* generally matched across multiple different image slices of the confocal stack. The image slice with the maximum Soma-print score was used to identify the cell’s axial position in the stack. **D)** Map of cells observed *in vivo,* showing 1,359 neurons that were matched to cells found in the corresponding *ex vivo* confocal image stack using the 3D Soma-print algorithm (1,615 total cells found *in vivo*, yielding a sensitivity of 84%). Cells are colored according to the axial plane in the stack yielding the match with the highest Soma-print similarity score, found as in **C**. Scale bar: 400 μm. **E)** Overlay (white) of cell maps obtained from *in vivo* (green, data of Fig. 4A) and HCR-FISH Round 1 using probes of GCaMP RNA (magenta, data of Fig. 4B). Cells found *in vivo* were paired to their counterparts in the confocal image stack across different image slices, hence we stitched the *ex vivo* cell map from patches assembled across multiple image slices by using the 3D Soma-print algorithm (**Methods**). Scale bar: 400 μm. **F)** Distributions of Soma-print similarity scores and corresponding parametric fits for pairs of best-matched and 2^nd^-best-matched cells, plotted in the same format as Fig. 2G for the data of **E**.

**Figure S6.**
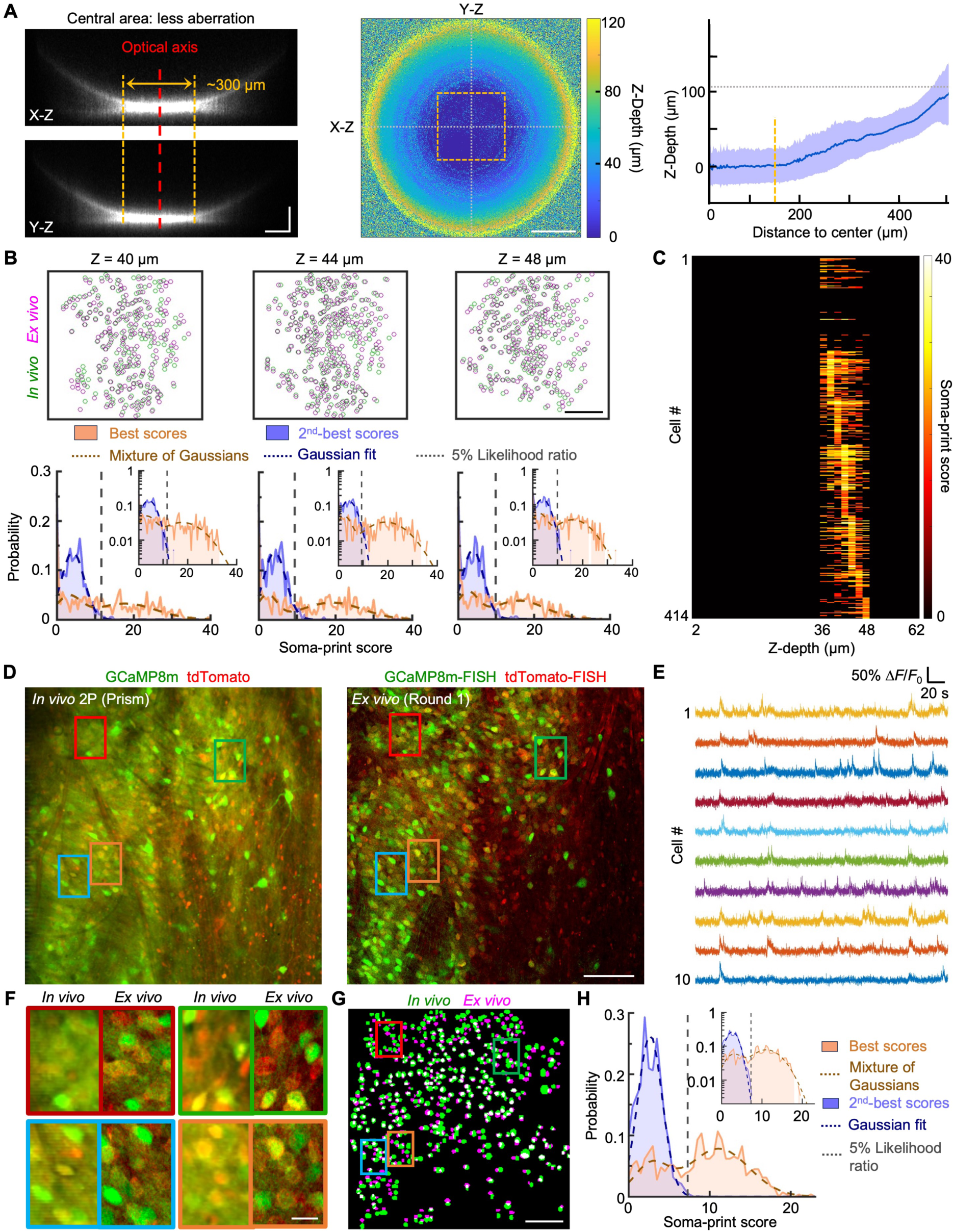
Field curvature of a GRIN microlens and alignments of Ca^2+^ videos, obtained in deep brain tissue via a GRIN microlens or a microprism, to postmortem RNA FISH images. **A)** We evaluated the field curvature of a GRIN lens (1 mm diameter, 4.38 mm length; 1050-006242, Inscopix) used for two-photon Ca^2+^ imaging in the dorso-medial striatum (**Methods**), showing a need to align neurons across multiple *ex vivo* confocal slices as in **Fig. S5**. *Left*: Cross-sectional projections from volumetric image stacks acquired using two-photon microscopy through a GRIN lens to image a fluorescent glass coverslip. The image of the glass surface has relatively low axial resolution but is also quite flat within a ∼150 μm radius from the optical axis. Scale bars: 100 μm (X and Y lateral dimensions) and 50 μm (Z, axial dimension). *Middle*: Color plots showing the axial displacement of the image of the slide as a function of the lateral displacement from the optical axis, determined as in **Figs. S2A** and **S5A**. For each (X, Y) position, we determined the axial displacement of the image of the slide by performing a Gaussian fit to the image brightness values as a function of Z and then taking the Z position corresponding to the peak of the Gaussian. Scale bar: 200 μm; *Right*: Mean ± s.d. axial displacement of the two-photon image of the fluorescent slide, plotted as a function of the radial displacement from the optical axis, averaged over all azimuthal angles. The vertical dashed line marks 150 μm radial displacement from the optical axis; we used the portion of the *in vivo* imaging field-of-view lying within radial displacements less than this. **B)** *Top*: Maps of cells obtained from *in vivo* (green) and *ex vivo* (magenta) datasets, showing that different subsets of cells observed *in vivo* through a GRIN lens are matched with counterparts found in several different axial planes in the *ex vivo* image stack (image planes were acquired with an axial spacing of 2 μm; shown are planes 4 μm apart). Scale bar: 100 μm. *Bottom*: Distributions of Soma-print scores for pairs of best-matched (orange) or 2^nd^-best-matched cell pairs (blue), along with parametric fits and semi-logarithmic versions (*insets*) of the plots, shown in the same format as in Fig. 5F for each of the image slices in the top row. **C)** Color plot for the same dataset as in **B**, showing the best Soma-print scores for all 414 cells found *in vivo,* when matched using the 2D Soma-print algorithm to the data from individual axial planes in the *ex vivo* image stack. The 3D algorithm then identifies for each cell the axial plane with the maximum Soma-print score. **D)** Corresponding *in vivo* and *ex vivo* images of spiny projection neurons from the dorsal medial striatum of a Drd1a-Cre ✕ Ai14 mouse. We labeled cells and acquired images in the same manner as in Fig. 5A, except that here the *in vivo* two-photon Ca^2+^ imaging was performed through a microprism implanted in the striatum. Rectangles enclose example subregions with neurons that can be readily tracked by eye across the image panels shown in **F**. Scale bar: 100 μm. **E)** Example traces of Ca^2+^ activity from neurons imaged in **D** via the microprism of an awake resting mouse. **F)** Magnified views of the regions enclosed by the color-corresponding rectangles in **D**. Within the image panel for each subregion, image frames are arranged from left to right to show the *in vivo* and HCR-FISH Round 1 data, respectively. Scale bar: 20 μm. **G)** Overlay of cell pairs matched by the Soma-print algorithm across *in vivo* (green) and reconstructed *ex vivo* (magenta) cell maps for the data of **D**, shown in the same format as in **B**. Scale bar: 100 μm. **H)** Distributions of Soma-print scores for pairs of best-matched cell or 2^nd^-best-matched cell pairs computed for the image of **D**. Data and parametric fits are shown in the same format as in **B.**

**Figure S7.**
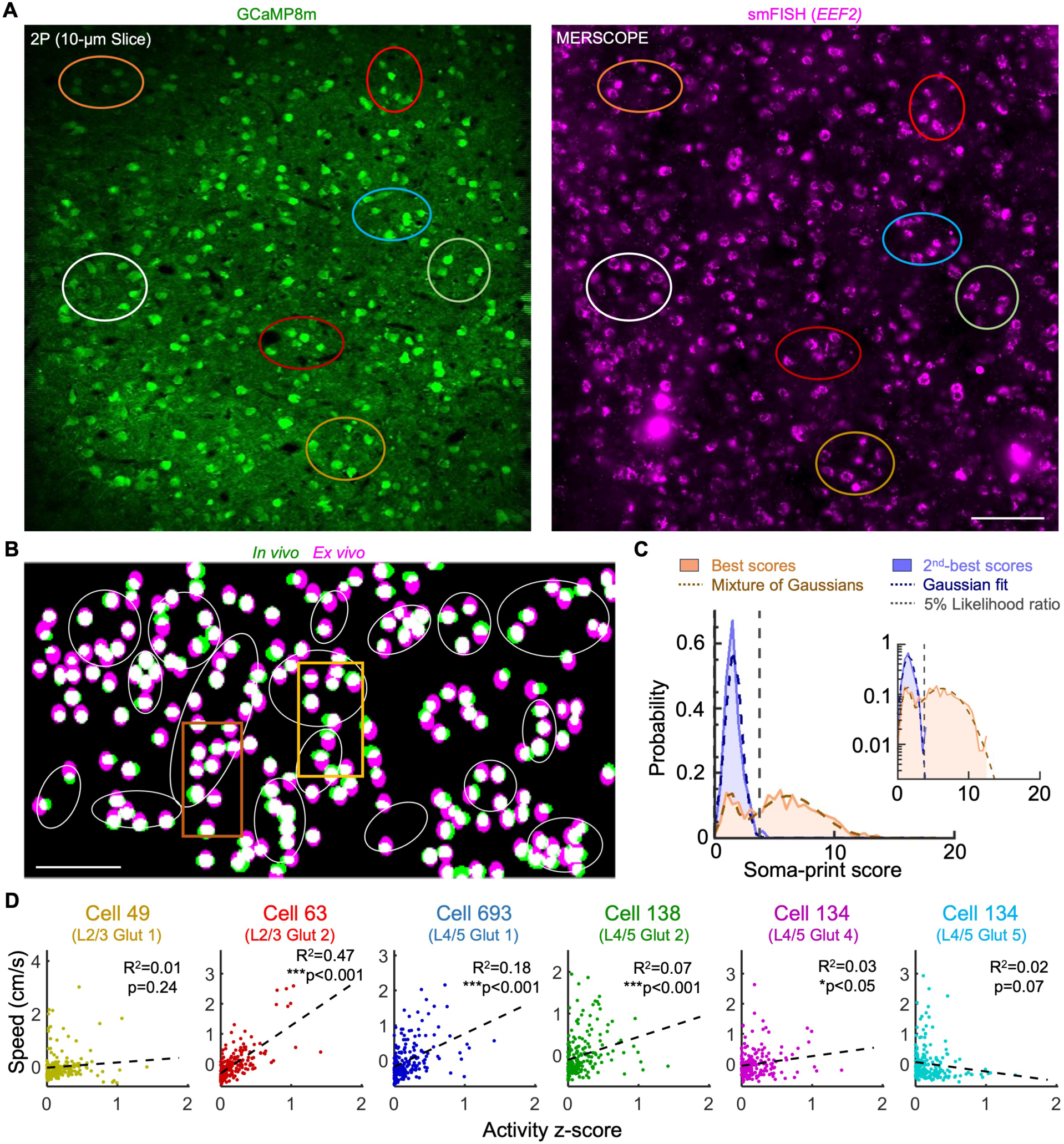
Validation of RNA quality using the MERSCOPE and alignment quantifications for MERFISH. **A)** Example data from a validation study in which we used the MERSCOPE (Vizgen) to check RNA abundance in tissue processed using our TRU-FACT procedures. *Left*: *Ex vivo* two-photon image of native GCaMP8m fluorescence from a 10-μm-thick tissue section from mouse motor cortex. *Right*: The corresponding single molecule FISH (smFISH) image from the MERSCOPE showing the detection of mRNA for the abundant, *EEF2* housekeeping gene in the same field-of-view as that in the left panel. The two images show good maintenance of detectable RNA and that cells’ morphologies were well preserved across our MERFISH tissue processing. Colored ellipses in **A** and **B** enclose subregions with cells that are readily matched by eye across the two images. Scale bar: 100 μm. **B)** Overlay (white) of matched *in vivo* (green) and *ex vivo* (magenta) cells (from Fig. 6B). White ellipses enclose the same regions as in Fig. 6B, in which neurons can be readily matched by eye across the two images. Areas within colored rectangles are shown at greater magnification in Fig. 6C. Scale bar: 100 μm. **C)** Distributions of Soma-print scores and corresponding parametric fits for pairs of best-matched and 2^nd^-best-matched cells, plotted in the same format as Fig. 2G for the data of **B**. Using the likelihood ratio cutoff of 0.05 (vertical dashed line), 196 out of the 272 cells observed *in vivo* were successfully matched to corresponding cells found postmortem (sensitivity: 72%). *Inset*: The same distributions and fits shown with a logarithmic scale on the *y-*axis. **D)** Linear regression plots of the mouse’s running speed versus the z-scored Ca^2+^ activity of 6 different example cells from each cluster in Fig. 6D**–F** (red: L2/3 Glut 2, blue: L4/5 Glut 2, yellow: L4/5 Glut 5, grey: others).

**Figure S8.**
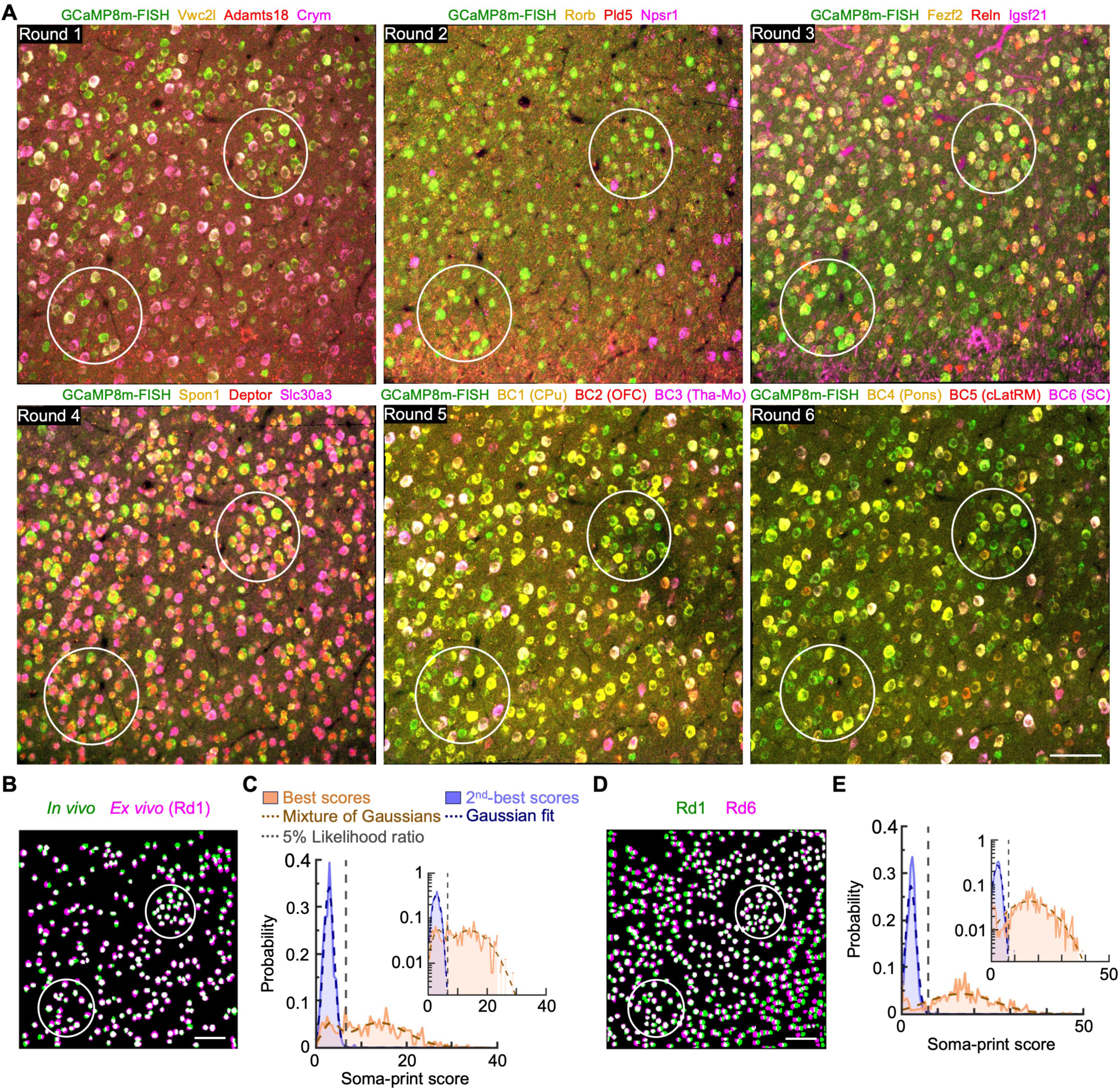
Registering 6 rounds of HCR-FISH and *in vivo* Ca^2+^ imaging data. **A)** Example confocal fluorescence images across 6 rounds of HCR-FISH, aligned to the corresponding *in vivo* Ca^2+^ imaging data (Fig. 7B shows the *in vivo* image). Rounds 1–4 of HCR-FISH used gene probes for the mRNA of GCaMP and 3 other genes per round, noted above each image in colored fonts. Rounds 5 and 6 involved gene probes for GCaMP and a total of 6 barcodes (termed BC1–6), expressed respectively in neurons with axonal projections in caudate putamen (CPu), orbitofrontal cortex (OFC), motor thalamus (Tha-Mo), the pons, contralateral lateral rostral medulla (cLatRM), and superior colliculus (SC). White ellipses enclose example cells matched across images. Scale bar: 100 μm. **B)** Overlay (white) of the cell maps determined by HCR-FISH using the GCaMP probe in Round 1 of **A** (magenta colored cells) or *in vivo* Ca^2+^ imaging data (green cells). Only cells that were matched across both modalities are shown. Scale bar: 100 μm. **C)** Distributions of Soma-print similarity scores and corresponding parametric fits for pairs of best-matched and 2^nd^-best-matched cells, plotted in the format of Fig. 2G, for the Round 1 images of panel **A** and Fig. 7B. Out of 328 neurons found *in vivo,* we successfully matched 273 to cells found *ex vivo* (sensitivity: 83%). **D)** Overlay (white) of the cell maps determined *ex vivo* using the GCaMP probe in Round 1 (green) or Round 6 (magenta) of HCR-FISH for the data of **A**. Only cells that were matched across both modalities are shown. Scale bar: 100 μm. **E)** Cells can be matched well across rounds of multi-round HCR-FISH, but not perfectly. The plots show distributions of Soma-print similarity scores and corresponding parametric fits for pairs of best-matched and 2^nd^-best-matched cells, plotted in the format of Fig. 2G, for the images from Rounds 1 and 6 of panel **A** and Fig. 7B. We matched 508 or 84% of 608 cells across the two rounds.

**Supplementary Video 1 | Example TRU-FACT alignments of *in vivo* neural Ca^2+^ activity and spatial transcriptomic data, using conventional, mesoscope, GRIN microlens-based, or miniature two-photon microscopy in awake mice, together with either HCR-FISH or MERFISH.**

The video shows five movie clips, showcasing the following five example alignments:

**1.** Somatosensory cortical neural Ca^2+^ activity imaged in an awake mouse with conventional two-photon microscopy, aligned to HCR-FISH data. The data are from the mouse of Fig. 1C.

**2.** Motor cortical neural Ca^2+^ activity imaged in an awake mouse with a two-photon mesoscope, aligned to HCR-FISH data. The data are from Fig. 4A**,C**.

**3.** Striatal neural Ca^2+^ activity imaged in an awake mouse via a GRIN microlens, aligned to HCR-FISH data. The data are from Fig. 5A.

**4.** Motor cortical neural Ca^2+^ activity imaged in an awake mouse with conventional two-photon microscopy, aligned to MERFISH data. The data are from Fig. 6B.

**5.** Motor cortical neural Ca^2+^ activity imaged in a freely moving mouse using a miniature two-photon microscope, aligned to HCR-FISH data. The data are from Fig. 5H. The side-by-side video clips of the concurrent Ca^2+^ activity patterns and mouse behavior are shown with a playback speed of ∼21.6× the original frame acquisition rate.

In each of the above examples, the white dashed ellipses enclose example regions in the *in vivo* and *ex vivo* images that can be readily matched by eye.

## Methods

### Mice

We used male and female C57BL/6J (Jackson Laboratory #000664), Ai14 (also known as Gt(ROSA)26Sortm14(CAG-tdTomato)Hze; Jackson Laboratory #007914), and Drd1a-Cre mice (also known as 129S6.FVB(B6)-Tg(Drd1a-cre)AGsc/KndlJ; Jackson Laboratory #02829). We generated double transgenic Drd1a-Cre/Ai14 animals by crossing homozygous Drd1a-Cre and Ai14 mice. Mice were 10–14 wks old at the start of experiments. Mice resided in a facility with a regular 12-h-light/12-h-dark cycle. The Stanford Administrative Panel on Laboratory Animal Care (APLAC) approved all animal experiments.

### Viruses

For studies of neural Ca^2+^ activity, we subcloned Addgene plasmids pGP-AAV-Syn-jGCaMP8m-WPRE (Addgene-plasmid #162375) and pGP-AAV-Syn-jGCaMP8s-WPRE (Addgene plasmid #162374), both produced by the HHMI GENIE Project. We used restriction enzyme BamHI (NEB R0136S) and HindIII-HF (NEB R3104L) to digest the plasmids, and with a Quick Ligation Kit (NEB M2200L) we ligated the jGCaMP8m and jGCaMP8s fragments, respectively, into a pAAV-CaMKII-BamHI-HindIII-WPRE backbone, which we had made previously^107^. The HHMI Janelia Research Campus Viral Vector Core facility packaged the AAV2/PHP.eB-CaMKII-jGCaMP8s and AAV2/PHP.eB-CaMKII-jGCaMP8m viruses.

To delineate neural projections, we generated 8 separate retroAAV barcode plasmids, each with a unique barcode. To create these barcodes, we used a random DNA sequence generator^156^ to produce a random 2000 bp sequence and then Primer3^157^ to select 8 probe sequences, each 100- or 200-bp in length, within the 2000 bp, optimized for annealing efficiency. To verify a lack of significant alignments with sequences in the mammalian genome, we evaluated candidate barcode sequences with BLAST.

We created barcode viruses by cloning a 448 bp promoter segment from the human Synapsin 1 gene on chrX:47,619,811-47,620,258 into an AAV2 inverted terminal repeat backbone^158^. Following this promoter, we incorporated the infrared protein miRFP670^159^ fused with Flag, HA, and Myc tags to allow detection by immunostaining^160^. After the open reading frame in each construct, we incorporated a unique barcode (100 bp for constructs 1-4 and 200 bp for constructs 5-8). To enhance expression levels, after each barcode we added a Woodchuck Hepatitis Virus Post-Transcriptional Regulatory Element (WPRE) and a Bovine Growth Hormone (Bgh) polyadenylation sequence. We sent the sequences to Addgene for synthesis and then GenScript commercially synthesized the viral constructs. We then cloned the constructs into bacterial expression plasmids and amplified the plasmids via maxi-preparation.

The HHMI Janelia Research Campus Viral Vector Core facility generated the retrograde adeno-associated viruses (rAAVs)^161^ to express each barcode in the brain. The resulting rAAV concentrations were as follows: Barcode #1 pAAV hSyn_iFP_Flag (4.67 × 10^12^ GC/mL); Barcode #2 pAAV hSyn_iFP_Flag (6.91 × 10^12^ GC/mL); Barcode #3 pAAV hSyn_iFP_Flag (8.28 × 10^12^ GC/mL); Barcode #4 pAAV hSyn_iFP_Flag (8.61 × 10^12^ GC/mL); Barcode #5 pAAV hSyn_iFP_Flag (1.15 × 10^13^ GC/mL); Barcode #6 pAAV hSyn_iFP_Flag (7.00 × 10^12^ GC/mL); Barcode #7 pAAV hSyn_iFP_Flag (5.20 × 10^12^ GC/mL); Barcode #8 pAAV hSyn_iFP_Flag (1.12 × 10^13^ GC/mL).

### HCR-FISH probes and amplifiers

We used multi-round HCR-FISH^155,162^ to differentiate between subtypes of neurons in the mouse motor cortex. We selected target genes based on published sequencing results^49^ and we picked 21 high-expressing genes with a differential expression pattern allowing the differentiation of subtypes of cells within a broader class of either layer 2/3 or layer 5 pyramidal neurons of the mouse motor cortex. With multi-round HCR-FISH, we were limited in each labeling round to 3–5 fluorescent labels, which could each be assigned to an individual gene or barcode. Within a given round, each label had to be assigned a unique amplifier sequence. To determine assignments of genes and their corresponding HCR amplifiers to specific rounds of labeling, we created groups of genes with similar expression levels in motor cortical neurons, based on the Allen Brain ISH map^163,164^, and assigned amplifiers to the individual genes or barcodes to be labeled within each round.

We sent the sequences of the target genes and barcodes to Molecular Instruments Inc. and bought ready-to-use HCR^TM^ RNA-FISH v3.0 compatible probe sets, customized for our selected genes and grouping. We used sets of 5 different HCR^TM^ v3.0 amplifiers (termed B1 to B5; Molecular Instruments Inc.), each conjugated to one of 4 different fluorescent dyes (Alexa fluorophores 488, 546, 594, and 647), allowing us to handle all possible gene target-to-fluorophore assignments. Marker genes and barcodes were assigned to the HCR amplifiers as follows:

B1: Vwc2l, Fezf2, Otof, Rorb, Erg, Barcode 1, Barcode 5;

B2: Spon1, Cux2, Igsf21, Npsr1, Cre, Ccdc3, Barcode 2, Barcode 6;

B3: Cpne5, Deptor, Adamts18, Pld5, Slc17a7, Barcode 3, Barcode 7, tdTomato;

B4: Crym, Slc30a3, Reln, Stard8, Met, Barcode 4, Barcode 8;

B5: Rspo1, GCaMP.

We also used a B7-488 amplifier-fluorophore pair to label the RNA sequence for GCaMP, and a B3-Cy7 amplifier-fluorophore pair to allow barcode labeling with an infrared fluorophore. (It should be noted that Molecular Instruments Inc. discontinued v3.0 RNA-FISH probe sets and amplifiers in May, 2025). To optimize FISH labeling, we adjusted the concentration of each RNA probe within the 4–40 nM range, based on the targeted gene’s level of RNA expression in our tissue samples.

### Surgical procedures

For cortical windows, we performed virus injections (first jGCaMP8- and then barcode-expressing viruses) from anterior to posterior and then made a craniotomy all in the same surgery. For deep brain imaging studies, we performed the craniotomy and implantation of the GRIN lens or microprism about 1 wk after virus injection.

Prior to surgery, we subcutaneously administered mice carprofen (5 mg/kg) and dexamethasone (2 mg/kg). We anesthetized mice with isoflurane (4–5% induction, 1–2% maintenance, both in O_2_) in a stereotactic frame. We maintained body temperature at 36.5 °C with a heating pad. We applied xylocaine (2–5 mg/mL) at the surgical site for topical anesthesia.

To express jGCaMP8 virally, we infused 500 nL of virus solution (2.0 × 10^12^ GC/mL) at each injection site (2–4 sites per brain area). Coordinates of those injection sites in cortex (mm from bregma: AP, ML, DV from pia) were as follows: **Fig. 1C**, M1: +1.0, 1.2, –0.3 and –0.7; S1: –1.0, 1.5, –0.3 and –0.7; **Fig. 3D**, M2: +2.0, 1.0, –0.3 and –0.7; **Fig. 4A**, M1: +0.6 and +1.6, 1.0, –0.3 and –0.7; M2: +0.6 and +1.6, 1.9, –0.3 and –0.7; **Fig. 6A, B**, M1: +0.5, 1.7, –0.3 and –0.7; **Fig. 7A, B**, M2 (RFA): +2.0, 1.0, –0.3 and –0.7; M1 (CFA): +0.5, 1.7, –0.3 and –0.7.

To validate the barcode design and test whether multiple barcodes can be expressed in one single cell, we injected mixed retroAAVs containing the eight artificial barcodes into the primary motor cortex (M1: +0.5, 1.7, –0.7) of the mouse, with each barcode diluted to the same concentration (0.9 × 10^12^ GC/mL). For viral expression of each barcode, we diluted all barcode retroAAVs to 4–5 × 10^12^ GC/mL before injection and infused 200–300 nL of virus per site (1 site for each brain area) during the same surgery as for injection of jGCaMP8-expressing virus. We injected these retroAAVs in various brain areas, as noted in the individual panels of **Figs. 3**, **7** (mm from bregma: AP, ML, DV): S1: –0.1, 1.8, –0.6 from pia; Tha-VM: –1.1, 0.8, –4.1; SN: –3.1, 1.5, –4.3; CPu: +1.0, 2.0, –3.5; OFC: +2.6, 1.8, –2.8; Tha-Mo: –1.0, 1.1, –3.5; Pons: –4.2, 0.4, –5.4; cLatRM: –5.5, –1.5, –4.7; SC: –3.4, 0.8, –1.8.

To perform the craniotomy, we first removed the skin atop the cranium and removed all soft tissues from the skull surface with a scalpel and scissors. Next, we used a 0.7-mm-diameter micro drill burr (#19007-07, Fine Science Tools) to perform a craniotomy above the virus injection site by thinning the skull. We used a glass capillary (Sutter Instrument, BF100-58-10), fashioned with a puller (Sutter Instrument, Model P-2000), to penetrate the brain. We loaded virus solution into the capillary and attached it to a Picospritzer-III (Parker Hannifin, 052-0500-900). We injected the solution into the brain at the rate of ∼50 nL/min. We waited 5 min before retracting the capillary.

In mice receiving a glass cranial window, we created a square craniotomy from AP 3.0 to –2.0 mm (relative to bregma), approximately matching the size of the implanted coverglass (5 mm wide). We dried the edges of the window implant with surgical eye spears (Butler Schein Animal Health, 1556455) and glued the window to the skull with ultraviolet (UV) light-cured adhesive (Loctite, 4305) while protecting the mouse’s eyes with aluminum foil to prevent UV-light damage. We affixed a stainless-steel annular head plate (10–12 mm inner diameter) to the cranium, for head-fixation during brain imaging, and filled the gap between the head plate and skull with dental cement (Fisher Scientific, 10000787).

For Drd1a-Cre × Ai14 mice receiving an implanted gradient refractive index (GRIN) microlens (1 mm diameter, 4.38 mm length; 1050-006242, Inscopix) or microprism (1.5 × 1.5 × 3.0 mm; OS PN160712BK01, OptoSigma) in the dorsal striatum, we followed similar procedures as those above except that we used a Nanoject II (Drummond) to inject 500 nL of AAV2/PHP.eB-CaMKII-jGCaMP8m virus into the dorsal striatum (AP +0.5, ML 1.8, DV –3.0 to –2.4 for GRIN lens experiments; AP +0.5, ML 1.8 and 2.4, DV –3.0 to –2.4 for microprism experiments).

In mice receiving an implanted GRIN lens (**Figs. 5A, S6A–E**), we installed the GRIN lens in the striatum as described previously^107^. In brief, we stereotaxically implanted a cannula above the dorsal striatum (centered at AP +0.5, ML 2.0) about 1 wk after virus injection. We prefabricated the cannula by fixing a custom-made Schott Glass coverslip (2-mm-diameter, 0.1-mm-thickness; TLC International) to the tip of a 3.5-mm long, extra-thin 18G stainless steel tube (Ziggy’s Tubes and Wires) using an optical adhesive (Norland #81). We ground off any excess glass with a polishing wheel (Ultratec). Using a 27G blunt needle, we aspirated cortical tissue down to DV –2.0 mm beneath the dura and implanted the exterior glass face of the cannula at DV –2.3 mm at a 10° lateral angle from the sagittal plane to target the dorsal medial striatum. We applied UV-light-cured adhesive (Loctite 4305) to seal the gap between the optical tube and the skull and attached a custom-designed headplate to the skull. Dental cement was applied to fix the optical tube and headbar to the cranium. 4 weeks after the cannula implantation, we assessed Ca^2+^ indicator expression in the dorsal striatum by inserting a GRIN lens (1 mm diameter, 4.38 mm length; 1050-006242, Inscopix) into the cannula and used a custom-built two-photon microscope equipped with a 0.8 NA microscope objective lens (CFI75 LWD 16× W, Nikon) to image the striatum while the mouse was head-fixed and placed on a running wheel. In mice that exhibited sufficiently uniform indicator expression, we secured the GRIN lens within the guide tube with the UV-light-cured adhesive.

In mice receiving an implanted microprism (**Fig. S6D–H**), we performed the implantation according to a published protocol (Hjort et al., 2024), albeit with some modifications. In brief, a 1.5 × 1.5 × 3.0 mm microprism was custom-assembled to a 3-mm-diameter #1 glass coverslip (Warner Instruments, CS-3R 64-0720) using optical adhesive (Norland #81). After virus injection, we used a 0.5-mm-diameter drill burr (#19007-05, Fine Science Tools) to create a 1.7 mm × 1.7 mm square craniotomy with the four corners positioned at AP +2.1 and +0.4, ML +1.3 and +3.0. We removed the center bone piece, cleaned the bone debris, and detached the dura from the edges of craniotomy. We used absorbable gelatin sponge (SURGIFOAM) wetted with Ringer’s solution to control bleeding and clean the brain surface. We held the attached microprism (1.5 × 1.5 × 3.0 mm; OS PN160712BK01, OptoSigma) using a 20G blunt needle attached to a vacuum, lowered the microprism to DV –2.8 mm at a maximum rate of 500 μm/min, and affixed it in place with UV-light-cured adhesive (Loctite 4305). We attached a custom-designed headbar to the skull and applied dental cement to fix the headplate and microprism. 4 wks after implantation, we assessed Ca^2+^ indicator expression with a custom-built two-photon microscope equipped with a 0.45 NA objective lens (CFI Plan lD 10× / WD 4.0 mm, Nikon).

After surgery, we transferred the mouse to a recovery cage and placed it on a heating pad until it awoke. We then returned the mouse to its home cage and provided food and water on the cage floor without use of a food hopper for 3–5 d. We administered carprofen (5 mg/kg) and dexamethasone (2 mg/kg) subcutaneously for up to 3 d after surgery to reduce post-surgical discomfort. For the first 7 d after surgery, we checked all mice daily for signs of distress. Mice typically recovered for 14–21 d before behavioral training sessions began.

### Behavioral assays

We employed two different behavioral assays for head-fixed mice, wheel running and pellet reaching. For the former assay, we head-fixed mice and placed them on a custom 3D-printed running wheel (a cylinder of 4.3 cm diameter and 10 cm length) equipped with a rotary encoder. For the latter assay, we placed mice in a cup-shaped holder that allowed them to sit on their hindlimbs and rest their forelimbs on a rod in front of a custom-designed automatic pellet presentation and dispenser system. A rotating wheel delivered sugar pellets one at a time in front of the mouse, accompanied by a 400 ms, 4 kHz tone. We first trained mice to eat pellets by mouth for 4 d (1 h training session per day). On the 5th day, we changed the position of the mice so they could no longer reach the pellet by mouth. Over a period of 2–7 d, the mice began using their right forelimb to retrieve the pellet and bring it to their mouth. A physical barrier prevented the mouse from reaching the food pellet with the left forelimb. We trained mice daily for another ∼4 d to retrieve the pellet with the right forelimb (1 h session per day), in the absence of Ca^2+^ imaging, until their pellet grasping performance was stable for 3 d. For each mouse, we then performed 3–4 sessions of joint Ca^2+^ imaging and pellet-reaching, with sessions typically spaced 2 d apart.

During the period that a mouse was involved in pellet reaching experiments, we restricted its food to maintain its body weight at ∼90% of its baseline value. For both wheel running and pellet reaching behavioral studies, we positioned a camera (The Imaging Source, DMK37BUX273) and infrared illuminator (Tendelux, AI4) in front of the mice to record their behavior. A custom system using National Instruments hardware (myRIO-1900) and software (LabVIEW 2019) synchronized the acquisition of two-photon Ca^2+^ videos with the behavioral readouts from the rotary encoder and camera. For each image frame recorded by the two-photon microscope (30 fps image acquisition), 3 image frames were acquired by the behavioral camera (90 fps image acquisition). In studies with the running wheel, the rotary encoder provided one measurement for each image frame taken by the two-photon microscope, yielding a 30-Hz-recording from the running wheel.

### Two-photon Ca^2+^ imaging

*In vivo* imaging sessions began 4–5 wks after virus injection, using either a two-photon mesoscope, a head-mounted miniature two-photon microscope (mini-scope), or a conventional two-photon microscope.

For studies involving the two-photon mesoscope (**Fig. 4, S5**), we performed large-scale Ca²⁺ imaging with an upgraded version of our published mesoscope^104,165^. This instrument, now equipped with faster electronics and a superior photodetection system, uses either one beam or a time-multiplexed set of beams from a tunable 80 MHz Ti:sapphire laser (MaiTai eHP DeepSee; Spectra Physics) to scan a 4 mm² area of brain tissue. For the studies in this paper, we performed imaging at 7.5 fps with one beam (30 mW; 940 nm wavelength for imaging GCaMP fluorescence) at a depth of 150 µm in tissue and took images of 2048 × 2048 pixels, with the width of each image pixel corresponding to a 1 µm span in tissue. In total, we acquired 5,000 image frames in about 11.1 min.

For studies involving Ca^2+^ imaging in freely behaving mice (**Fig. 5G–J**), we used a miniature two-photon microscope (TRANSVISTA SUPERNOVA-100; headpiece FIRM-U; 3× objective, 0.7 numerical aperture, working distance 1.0 mm; Transcend Vivoscope Biotech Co. Ltd, Beijing, China)^117^. This microscope uses an ultrashort-pulsed laser (920 nm; Axon 920-2 TPC, Coherent Corp.) and has a built-in blue LED for wide-field one-photon imaging of the brain surface. For the studies in this paper, we acquired two-photon images at 9.21 fps using 60 mW of laser illumination power. We imaged neural Ca^2+^ activity at cortical tissue depths of 140 µm, 160 µm, and 180 µm, using the device’s electronically tunable lens to adjust the focal plane. The two-photon images had 512 × 440 pixels (FOV size: 467 µm × 401 µm). In each session, we acquired a total of 3,000 or 6,000 image frames in 5.4 min or 10.8 min for each plane. We used the infrared camera and lighting provided by the SUPERNOVA-100 system to take videos of the mouse’s behavior (30 fps, 1600 × 1200 pixels) in a rectangular open field enclosure (40 cm × 30 cm).

Except for *in vivo* imaging done with the two-photon mesoscope or mini-scope, for all other Ca²⁺ imaging we used a custom-built two-photon microscope with both one- and two-photon imaging capabilities. This microscope was equipped with an ultrashort-pulsed laser (MKS Spectra-Physics, Insight X3, 680–1,300 nm tunable), an electro-optical modulator (Conoptics, EOM 350-80-LA-02 with a 302RM driver), and two galvanometric mirrors, one linear and one resonant. The latter galvanometer had a resonant frequency at 8 kHz (Novanta Photonics, 6215H, CRS8K), allowing 30-fps-image-acquisition (512 × 512 pixels) under the control of ScanImage 5.6.1 software (Vidrio Technologies). We customized the software to include additional modules written in MATLAB (MATLAB 2019a, MathWorks) for automated acquisition of image tiles and stacks and fine alignments between *in vivo* and *ex vivo* imaging sessions with the same brain. A dichroic mirror (Semrock, FF555-Di03-25x36, 555 nm edge) separated fluorescence emissions into green and red detection channels, each of which had a bandpass fluorescence filter (Semrock, FF02-525/40-25 and FF01-607/70-25, respectively). A pair of photomultiplier tubes (PMT), controlled by custom software written in LabView 2019 (National Instruments), detected fluorescence photons. An electronically gated PMT (Hamamatsu, H11706P-40) captured green fluorescence signals, and a standard PMT (Hamamatsu, H10770PA-40) captured red fluorescence.

For studies performed with either the conventional two-photon microscope or the two-photon mesoscope, we positioned head-fixed mice under the objective lens. The mesoscope studies used a water-immersion objective lens optimized for large-scale two-photon imaging^97^ (1.0 numerical aperture (NA) for fluorescence collection, 2.5 mm working distance; Jenoptik). For mice with an implanted cranial window or GRIN microlens (1 mm diameter, 4.38 mm length; 1050-006242, Inscopix), the objective lens of the two-photon microscope was a 16× water immersion lens (Nikon CFI75 LWD 0.8 NA, 3.0-mm-working distance). For mice with an implanted microprism (1.5 × 1.5 × 3.0 mm; OS PN160712BK01, OptoSigma), the objective lens was a Nikon 10× CFI Plan lD 0.45 NA objective with a 4.0-mm-working distance.

To reliably image the same individual neurons across multiple *in vivo* and *ex vivo* imaging sessions, we performed a series of alignment steps, starting with coarse alignments of the mouse head’s position and orientation, and then proceeding to finer alignment steps intended to attain cellular level registration across sessions. At the start of each imaging session, we oriented the mouse’s head in a consistent direction and placed it on a custom-assembled 4-axis mechanical stage with 2 translational (using a motorized XY stage for lateral positioning, Sutter, MP-285) and 2 rotational degrees of freedom (using two-stacked tip-tilt goniometers with 40 deg range, B54-60LN, Suruga Seiki). The stage resided on a lab jack, which allowed coarse axial positioning, and the microscope objective lens was held by a piezoelectric stage (PI, Q-545.240 with a E-873.1AT controller), which allowed fine axial adjustments of the focal plane.

To align the surface of the mouse’s glass implant to be perpendicular to the optical axis of the microscope, we used the 4-axis stage and light from a laser pointer (CPS532-C2, 0.9 mW, Thorlabs) that was set perpendicular to the optical table and parallel to the microscope’s optical axis (**Fig. S1A–C**). We used the 4-axis stage to align the mouse such that the reflection of the laser pointer illumination off the mouse’s glass implant returned directly back to the laser pointer, thereby ensuring that the mouse’s glass implant was orthogonal to the microscope’s optical axis. We then used the microscope’s one-photon fluorescence imaging capability to obtain a live image stream providing an overview (1.7 × 1.7 mm^2^) of the glass implant and the brain tissue beneath it. While looking at a stored, corresponding overview image from the mouse’s first imaging session, we translated the mouse (or its fixed head) laterally until the current overview image visually matched the stored image.

For fine lateral alignments of the mouse (or its fixed head) and fine axial adjustments of the image focal plane, we then switched from one- to two-photon imaging (0.7 × 0.7 mm^2^), again using a live image stream and a stored image from the first imaging session. We made fine adjustments to the lateral position of the mouse (or its fixed head) using the 4-axis stage and to the image focal plane using the piezoelectric stage, until the current image frame of the two-photon image stream visually matched the stored image.

Once these alignments were done, if the imaging session involved a live mouse, we then proceeded to Ca^2+^ imaging of neural activity (30 fps, 512 × 512 pixels). We used a laser illumination power at the specimen of ∼20–50 mW for imaging at tissue depths of 200–350 μm from the glass implant, and ∼80–100 mW for imaging at depths of 500–620 μm. For joint Ca^2+^ imaging and MERFISH studies, we took a series of 10-min recordings of Ca^2+^ activity as a head-fixed mouse ran on a wheel under the microscope objective lens, with each 10-min-recording acquired at one of 10 different tissue depths ranging from 200–350 μm beneath the brain surface, spaced 15 μm apart. For mice that performed the pellet reaching task (see *Behavioral assays* above), we performed a pair of 10-min-recordings of Ca^2+^ activity in cortical layer 5 at a depth of ∼550 μm beneath the pia. In cases when the imaging session involved an isolated mouse head, instead of imaging Ca^2+^ activity we proceeded to the optomechanical preparations done prior to brain tissue slicing (see next section).

For mice imaged with the two-photon mini-scope, prior to aligning the mini-scope we took a one-photon fluorescence image (∼10 mm^2^) of the brain tissue surface. We then aligned the microscope’s optical axis to be perpendicular to the mouse’s cranial window while it was head-fixed, using the reflection of light from a laser pointer (CPS635R, 1.2 mW, Thorlabs) off the glass cranial window. To do this, we used a custom 5-axis stage to adjust the position and orientation of the head-fixed mouse and another custom 3-axis holder to adjust the yaw, pitch, and roll coordinates of the mini-scope (step 1, **Fig. 5G**). While the mouse was head-fixed, we acquired a volumetric two-photon image stack that included the brain surface, allowing us to map the two-photon image data to the corresponding location in the one-photon image taken earlier. Once we had identified a satisfactory field-of-view for two-photon imaging, we raised the mouse’s head to the headpiece of the mini-scope and glued (MMA Structural Adhesive 1320H, Huitian) its lower baseplate to the mouse’s implanted headbar (step 2, **Fig. 5G**). Once the glue had dried, the mini-scope headpiece could be detached and reattached multiple times to the lower baseplate, allowing stable imaging of the same fields-of-view simply by mating the upper baseplate of the mini-scope to the lower baseplate that remained on the mouse (step 3, **Fig. 5G**). After all *in vivo* imaging sessions were completed, tissue processing and sectioning proceeded as shown in **Fig. S1B** or **S1C**.

### Measurements of field-of-view curvature across optical focal planes

To perform measurements of field curvature for either the two-photon microscope (512 pixels × 512 pixels × 161 slices with 0.5 μm spacing; **Fig. S2A**), the confocal microscope used for HCR-FISH image acquisition (512 pixels × 512 pixels × 81 slices with 0.5 μm spacing; **Fig. S2A**), the two-photon mesoscope (2048 pixels × 2048 pixels × 191 slices with 1 μm spacing; **Fig. S5A**), and GRIN lens imaging on the conventional two-photon microscope (512 pixels × 512 pixels × 191 slices with 1 μm spacing; **Fig. S6A**), we took volumetric image stacks of a fluorescent glass slide FluorCal (Valley Scientific, FC-OCS-EC) placed perpendicular to the optical axis on the specimen stage.

For the image stacks taken on the confocal or two-photon microscopes, with or without the GRIN lens, we cropped each stack by removing the outermost 6 pixels from each lateral side of the stack, yielding image slices of 500 × 500 pixels. We kept the image slices taken on the two-photon mesoscope at their original size of 2048 × 2048 pixels. Using these stacks, we then determined the image slice at which the fluorescence intensity was brightest for each lateral displacement relative to the optical axis. To do this, for each set of pixels that all had identical lateral coordinates but different axial coordinates, we fit a Gaussian function to the fluorescence intensity values as a function of axial depth. We took the axial coordinate value determined by this fit to have the peak fluorescence, and we compared how these axial coordinates varied across the imaging plane laterally. We averaged these axial coordinate values across a circular region of 20 μm radius centered on the optical axis as the reference depth. For all lateral coordinates, we tabulated the mean axial coordinate of peak fluorescence intensity, averaged over all pixels at the same radial distance from the optical axis.

### Optomechanical preparations for brain tissue slicing

At the end of each mouse’s last Ca^2+^ imaging session, we used our custom two-photon microscope or the commercial miniature two-photon microscope to acquire volumetric two-photon image stacks (2–5 μm axial spacing between image slices) that included the *in vivo* imaging field-of-views.

For mice with a glass cranial window, we then took laterally tiled sets of one- and two-photon images; we skipped this step for mice imaged with an implanted microprism or GRIN microlens or with the miniature two-photon microscope. We took a laterally tiled set of one-photon images at a tissue depth at which the brain surface’s blood vessels were most clearly visible. We took a laterally tiled set of two-photon fluorescence images at the tissue depths where the *in vivo* imaging field-of-views were sampled. These paired sets of tiled images provided visual references for the location at which we had acquired the volumetric image stack. For mice with an implanted microprism or GRIN microlens, the glass implant itself provided such a reference.

Next, we removed the mouse from the microscope stage and transcardially perfused it first with phosphate buffered saline (PBS) (10× PBS solution (Thermo Fisher, AM9625) diluted with RNase-free water (Thermo Fisher, 10977023) and then with 4% paraformaldehyde (PFA) (Electron Microscopy Sciences, 1224SK)). For brains to be analyzed by HCR-FISH, we removed the entire head and stored it in a solution of 4% PFA for 12–72 h at 4°C. For brains to be analyzed by MERFISH, we stored the head in this solution for 1–24 h at 4°C. During perfusion and subsequent handling of the brain, to protect the RNA from degradation we wore face masks and gloves and added RNase inhibitor (New England BioLabs, M0314L, 0.2 units/μL) to the PBS. Prior to perfusion of mice to be studied with MERFISH, we sprayed an aerosolized RNase inhibitor (RNaseZap, Thermo Fisher, AM9780) onto the tools and gloves to be used to touch the brain tissue. After fixation of the mouse’s entire head, with the glass implant still installed, we proceeded to brain tissue slicing.

Below, we describe 3 different approaches for slice preparation. The first method is arguably the most straightforward to implement and relies on a cylindrical post with optically flat faces to maintain parallel planes during imaging and tissue slicing. The second and third methods do not involve an optical post but instead require customized holders of the brain tissue. Common to the first two approaches was a key step that involved making a mold to capture the shape of the brain’s ventral side. The insight here was that the brain could be placed back into this mold with sufficient accuracy that the brain’s dorsal surface resumed its original orientation.

Across all of these approaches, two main strategies governed the alignment and preparation of brain tissue slices. First, to maintain sets of parallel planes during imaging and slicing, we used flat reference planes that we created in a gel or OCT embedding holding the brain, parallel to the focal planes imaged in the live mouse. Similarly to the *in vivo* imaging sessions, we imaged the tissue slices aligned orthogonally to the optical axis of the confocal microscope or the MERSCOPE. Second, to relocate a field-of-view and track its orientation in the postmortem tissue block or slice, at each step of tissue processing we took new images of the asymmetrically cut tissue blocks and noted updated sets of landmarks (reference points in tissue) to maintain an unbroken record of the orientation and location of the *in vivo* imaging field-of-view.

#### Alignment using an optical post

In this approach (**Fig. S1A**), we used a custom cylindrical glass post (4.0 mm diameter × 6.0 mm long, BMV Optical) with opposing faces that were optically flat and mutually parallel. We dissected the postfixed head of the mouse, removing the ventral portion of the skull while keeping intact the brain, cranial window, and dorsal portion of the skull. We glued one face of the glass post to the top surface of the cranial window, making both faces of the post parallel to the tissue planes imaged *in vivo*. We flipped the brain, so that its ventral side faced up, and glued the post’s opposite face to the specimen disc of a vibratome (Leica VT1000S), making this disc also parallel to the planes imaged *in vivo.* Next, we embedded the brain and disc in a 2.5% agarose gel (Lonza, SeaPlaque, 50100) in PBS (Thermo Fisher, AM9625; diluted with RNase free water, Thermo Fisher, 10977023). After putting the disc into the vibratome parallel to the cutting plane, we cut the gel block parallel to the specimen disc from the brain’s ventral side and thus also parallel to the planes imaged *in vivo*.

To create a mold of the ventral portion of the brain, we used a razor blade to manually cut the gel block into approximately two equal portions, with the cut approximately parallel to the face of the disc. We removed the ventral portion of the cut gel, exposing the ventral portion of the brain. We then detached the intact brain from the headbar, cranial window, and remaining portion of the skull. To preserve information about the brain’s orientation, we made 2 pairs of characteristic, asymmetric cuts in the brain tissue as landmarks, well outside the field-of-view imaged *in vivo.* These cuts ensured that the tissue slices to be made had a readily recognizable parallelepiped shape, which allowed us to easily distinguish the top and bottom sides of the slice and to relocate the cells of interest. After making these cuts, we placed the brain into the ventral gel mold and glued the flat surface of the mold to another vibratome disc. We took photographs of the cut brain in the mold, measured the distances from the edges of the brain block to the field-of-view imaged *in vivo*, and confirmed that we could re-find the cells of interest under the microscope. We made another 2.5% agarose gel mold surrounding the dorsal portion of the brain and the exposed portions of the first gel. We placed the second disc into the vibratome and made tissue sections (100–300 μm thick for HCR-FISH studies and 600–1,000 μm thick for MERFISH studies), with the slices cut parallel to original planes imaged *in vivo* by the flat surface just created.

For MERFISH studies, we then took the brain block, dehydrated it sequentially in solutions of 10%, 20%, and then 30% sucrose (Sigma-Aldrich, S0389) in RNase-free PBS at 4°C^166^, and embedded it in OCT (Fisher HealthCare, 4585). The OCT embedding retained the flat parallel planes of the brain block, as we made the embedding by placing the brain block on one microscope slide and then sandwiching the brain block using another microscope slide. To ensure that the sides of the glass slide sandwich had identical spacing, we created two sets of spacers by cutting several other glass slides in half, stacking them into two spacers of exactly equivalent height, and gluing them to each other and to the bottom slide of the sandwich with superglue (SUPER GLUE, 3EHP1). We confirmed the parallelism of the two glass slide sandwiches by checking that the reflections of the laser illumination (as described above) off the two spaced slides were merged. We made the OCT embedding by pouring liquid into the sandwich with a 4-side plastic barrier surrounding the tissue block and holding the liquid OCT in place and then freezing the assembly in a –20°C freezer. After freezing, we detached the embedded tissue block from the slide sandwich, removed the plastic barrier, and placed the tissue block on a flat reference surface created by cutting a blank OCT block attached to the specimen disc of a cryostat (Tanner Scientific, TN50) equipped with a MX35 Premier+ blade (Epredia, part #3052835) to make 10-μm-thick slices. We cut tissue slices and placed them on a specialized MERFISH slide (see MERFISH section below).

#### Alignment using direct embedding of a whole post-fixed brain

This method (**Fig. S1B**) allowed us to perform tissue slicing parallel to the planes of *in vivo* imaging without use of an optically flat glass cylindrical post. But, unlike the slicing method described above, the method here requires a custom mechanical holder for the mouse’s headplate and an accompanying, custom 5-axis mechanical stage to assist in the creation of the mold of the brain. A main benefit of this approach is its compatibility with many different mouse preparations; this generality is achieved by using the 5-axis stage to re-align the postfixed brain under the microscope to the same orientation used for *in vivo* imaging, prior to making the mold of the brain.

We first removed the ventral portion of the cranium from the post-fixed head of the mouse, as in the first method above, and inserted the headbar into the custom holder. We placed the specimen disc of the vibratome onto the flat surface of the custom 5-axis stage that was perpendicular to the optical axis, and used the custom holder to hold the brain above the disc, leaving some air space below the brain for casting the gel mold. We then used the 5-axis stage to adjust the orientation of the mouse head to be the same as it was during *in vivo* imaging. For example, in mice with an implanted cranial window, we used the same visible laser pointer (CPS532-C2, 0.9 mW, Thorlabs) as for *in vivo* imaging alignment and similarly adjusted the orientation of the mouse head so that the cranial window was perpendicular to the imaging microscope’s optical axis. Additionally, we could use the 5-axis stage and the microscope to re-find the same field-of-view imaged *in vivo.* Once the mouse head had resumed the proper orientation, we cast the gel mold surrounding the entire brain and headplate, on top of the specimen disc. We then proceeded as described above to cut the gel into two portions, remove the headbar, make characteristic asymmetric cuts in the brain, replace the brain into the mold, document the field-of-views’ positions, re-embed it in a second mold, place the disc into the vibratome, and make brain tissue slices for HCR-FISH studies. The resulting tissue slices can also be further processed (as described above) to prepare 10-μm-thick slices for MERFISH studies.

#### Alignment using direct embedding of a brain tissue block

In this approach (**Fig. 1B, S1C**), we manually dissected a brain tissue block containing the field-of-view imaged *in vivo*. We then trimmed the block asymmetrically, as described above, with one flat surface of the trimmed block comprising tissue that had directly contacted the flat optical implant, whether it was a glass cranial window, microprism, or GRIN microlens. We performed another round of fixing in 4% PFA for 30–120 min and then sequential dehydration in sucrose as described above. After dehydration, we took the brain block and placed its flat tissue surface onto a glass slide and embedded the brain block in OCT with another glass slide sandwich (as described above). After freezing the sandwich at –20°C, as described above, we extracted the embedded brain block, placed it on a flat reference surface created by cutting a blank OCT block in the cryostat, and used the same blade to make tissue slices (10 μm thick for MERFISH studies; 50–100 μm thick for HCR-FISH).

#### Selection of the appropriate optomechanical alignment method

We present three variants of our optomechanical alignment process (**Fig. S1A–C**). Unlike the other two methods, alignment using an optical post (**Fig. S1A**) requires no customized tools and is relatively insensitive to operator skill. We therefore recommend this approach for beginners. Users will have to purchase an optical post of the correct size, and there is some additional labor associated with gluing and detaching the optical post from the glass cranial implant.

Alignment involving a direct embedding of the whole post-fixed brain (**Fig. S1B**) requires a custom-made holder and is a bit more challenging but faster to execute than the beginners’ method. Compared to the third method (**Fig. S1C**) that uses OCT and a cryostat for cutting thin slices, the first two alignment methods, which both use agarose and a vibratome for cutting thick sections, are generally more prone to misalignments of the tissue cutting plane owing to the relative softness of an agarose embedding as compared to frozen OCT. We recommend choosing the direct embedding approach (**Fig. S1B**) when the experiment or data processing can tolerate a modest level of tissue distortion or when a cryostat is unavailable.

Alignment using direct embedding of a tissue block in OCT (**Fig. S1C**) and cutting slices with a cryostat is the most generally applicable of the three optomechanical variants, but it also requires prior training and experience with cryosectioning. In our experience, this method is the best choice for experiments involving an implanted microprism or GRIN microlens or those in which thin tissue sections (10 μm) are needed for the chosen spatial biology technique (*i.e.*, most high-plex methods).

### HCR-FISH labeling and confocal fluorescence imaging

After following one of the three alignment procedures described above, we performed multi-round HCR-FISH^155,162^ studies with the tissue slices. In each round, we sought to measure the relative expression levels of a total of 3 genes for which the RNA molecules were labeled with corresponding fluorescent Alexa dyes, respectively excited with 546 nm, 594 nm, and 647 nm illumination. Each round also included a GCaMP gene probe labeled with Alexa-488, chosen deliberately to allow us to aggregate into one detection channel the fluorescence photons from both GCaMP and the Alexa-488 dye label of GCaMP mRNA. After each round of labeling, we measured gene expression patterns by confocal microscopy. After each round of imaging, we removed the HCR-FISH signals by incubating the slides in DNAse I overnight and then started the next round. We used a 6-gene panel for L2/3 slices and a 12-gene panel for L5 slices, which we examined in 2 and 4 rounds of HCR-FISH labeling, respectively.

Our procedures for multi-round HCR followed published protocols but with slight modifications^155,162^. In brief, we incubated PFA-fixed, floating mouse brain tissue slices in 70% ethanol for 1–7 days. We then transferred the floating slices to 8% sodium dodecyl sulfate (SDS) solution in 2× saline sodium citrate (SSC) and incubated them for 30 min at room temperature with slight shaking. We then washed the slices in 2× SSC for 5 min and repeated this washing step three times. We then replaced the medium with HCR hybridization buffer (Molecular Instruments), incubated the slices in 37 °C for 10 min, added the probes at their optimized concentrations in the hybridization buffer, and incubated the slices overnight. The next day, we washed the hybridized slices with HCR washing buffer (Molecular Instruments) at 37 °C for 10 min, and repeated this washing step three times. We briefly washed the slices in 2× SSC and then replaced the medium with HCR amplification buffer (Molecular Instruments). In parallel to these steps for labeling the slices, we prepared the HCR amplifiers by heat-shocking each of them at 95 °C for 90 s and then snap-cooling them at room temperature for 30 min. After having prepared the amplifiers in this way, we added them to the buffer holding the tissue slices and incubated the slices overnight (12–18 h) with constant shaking. On the third day, we washed the slices with 2× SSC for 10 min, thrice repeating the washing. For the first round of labeling, we also incubated the slices with DAPI for 5 min and then washed them with 2× SSC for 10 min. We then placed the labeled slices onto a glass slide, mounted them with EasyIndex (refractive index of 1.52; LifeCanvas Technologies, #EI-100-1.52) and a coverslip (#1 thickness), and proceeded to image the mounted slices with confocal microscopy.

We used a Leica SP8 confocal microscope equipped with white light laser illumination to acquire tiled image stacks of the tissue sections. For fluorescence excitation, we scanned the slices with co-aligned beams from a 405-nm-emitting laser diode and the white-light-illumination laser (spectrally filtered to suit the fluorophores used). We acquired images at a pixel bit depth of 16 bits. For the microscope objective lens, we used an HC PL APO CS2 20× / 0.75 IMM lens (Leica) with a tunable correction collar that we adjusted for oil immersion (refractive index of 1.518). We acquired image slices of 1024 × 1024 pixels with a line-scanning speed of 400 Hz and averaged over a pair of line scans for each line in the digital image, with laser illumination powers at the specimen plane of approximately 17 μW for 405-nm-illumination, 16 μW for 488-nm-illumination, 125 μW for 546-nm-illumination, 207 μW for 594-nm-illumination, and 159 μW for 647-nm-illumination. Image slices in each stack were axially spaced by 2 μm. We collected fluorescence across 3 rounds of scanning, with up to two colors of fluorescence detection per round. Excitation wavelengths and emission bands for the three rounds were as follows. Round 1: 405 nm excitation, 430–490 nm emission; 647 nm excitation, 660–750 nm emission. Round 2: 488 nm excitation, 510–570 nm emission; 594 nm excitation, 610–660 nm emission. Round 3: 546 nm excitation, 555–590 nm emission. For image digitization, we did not apply an intensity offset for any of the color channels. We axially downsampled image stacks by averaging over sets of 3 successive image slices, yielding a net axial spacing of 4 μm between image slices in the downsampled stack. For near-infrared fluorescence confocal imaging of Cy7-conjugated HCR amplifiers, we used a Leica Stellaris8 DIVE confocal microscope and the same objective lens and settings as noted above, with the exception of the excitation wavelength of 730 nm and the emission band of 735–803 nm.

After confocal imaging was completed, we prepared for the next round of HCR labeling. We removed the coverslip covering the slices and washed the slices twice in 2× SSC. We then replaced the medium with DNAse I buffer (10×, Invitrogen, Catalog #AM8170G, diluted to 1× with RNAse free water) and washed the slices for 5 min. We then incubated the slices with DNAse I (Thermo Scientific, Catalog #EN0521, diluted 50× with DNAse I buffer) for 1 h at 37 °C. To cleave all remaining HCR probes and amplifiers, we replaced the buffer again with freshly prepared DNAse I and incubated the slices overnight at 37 °C. We further incubated the slices with 67% formamide (Thermo Scientific, Catalog #17899) in 2× SSC for 1 hour at 37 °C. We then washed the slices with 2× SSC, and repeated the above procedures for the next round of hybridization reaction and confocal imaging.

### MERFISH imaging

We mounted 10-μm-thick slices onto MERSCOPE slides (MERSCOPE Slide Box, Vizgen, 10500001), dried them in a cryostat for 20–30 min, washed the slides 3 times with RNase-free PBS at room temperature, and dried them for 60–90 min according to Vizgen protocols (91600002, 91700118). As the tissue slides dried, we imaged them with the two-photon microscope to visually compare the resulting fluorescence images, based on GCaMP’s native fluorescence, to a saved image of the mean, time-averaged Ca^2+^-related fluorescence intensity from the *in vivo* imaging session. We displayed both images side-by-side on the same computer monitor in the two-photon imaging room. By visually identifying patterns of cells’ spatial arrangements in the tissue slice that closely resembled those in the *in vivo* image, we determined which tissue slice or slices and the FOV’s lateral position within the slices on the MERSCOPE slides best matched the cells imaged while the mouse was alive.

After overnight permeabilization of the slides in 70% ethanol, we gave them to one of two different imaging facilities equipped with a commercial MERSCOPE (Vizgen) system. We sent the slides by overnight shipping (Vizgen protocol 1.5.092023), immersed in 70% ethanol and surrounded with ice packs, to the UCSD Center for Epigenomics. Alternatively, we provided samples immersed in 70% ethanol at 4 °C to the Stanford Protein and Nucleic Acid facility for MERFISH imaging with 500 genes. In both cases, we provided the MERSCOPE 500 Gene Imaging Kit (Vizgen, 10400006), MERSCOPE Sample Prep Kit (10400012), MERSCOPE Sample Verification Kit (Mouse, 10400008), and MERSCOPE PanNeuro Cell Type Panel 500 Gene (Mouse, 10400121) to the imaging facility, which followed Vizgen protocol (91600002) for sample preparation and MERFISH imaging.

### Analyses of pellet reaching behavioral data

We analyzed all food pellet reaching behavioral videos using DeepLabCut^167^ pose-estimation software to track the positions of the mouse’s right paw, mouth, and the food pellet. To identify time intervals within the behavioral videos that likely contained an attempt to reach and grasp the pellet, we computationally identified image frames in which the horizontal distance between the right paw and the pellet was less than 5 mm. We then visually inspected these and temporally adjacent image frames and determined specific frames corresponding to the onset and offset times of the reaching, grasping, pellet lifting and eating phases of the behavior.

### Analyses of neural Ca^2+^ activity

To extract neurons and their activity traces from Ca^2+^ videos, we used the EXTRACT cell extraction algorithm^168,169^. All further analyses of and figures showing neural activity used traces from EXTRACT.

To analyze Ca^2+^ videos acquired in mice that performed running or pellet-reaching behaviors, we first pre-processed the videos by temporally downsampling them from 30 fps to 15 fps. To computationally correct for lateral displacements of the brain during Ca^2+^ imaging, we aligned the image frames of the downsampled video with the NoRMCorre image registration software package^170^. To do this, we performed rigid image alignments and set the upsampling factor in NoRMCorre for subpixel registration to 30 and the maximum allowable pixel displacement to 30. Using the aligned video output from NoRMCorre, we identified individual neurons and their Ca^2+^ activity traces with EXTRACT^168,169^. To initialize EXTRACT, in some cases we provided it with a set of cells’ spatial filters that we generated with Cellpose^171^ cell segmentation software, as applied to a single image computed as a pixel-wise, temporally averaged version of the entire motion-corrected Ca^2+^ video. In other cases, we augmented the final set of active cells identified by EXTRACT with an additional set of quiescent cells that were only identified by Cellpose. Within EXTRACT, we set the mean cell radius to be 20 μm and applied a non-negativity constraint to the output Δ*F*(*t*)/*F*₀ traces of Ca^2+^ activity.

To analyze neural Ca^2+^ activity traces from mice that ran on a wheel, we computed *z-*scored versions of each cell’s Ca^2+^ activity trace as provided by EXTRACT, using each cell’s mean value of Δ*F*/*F*₀ computed over the entire movie. To relate the Ca^2+^ activity of individual cells to running speed, we determined mean values of the cell’s *z-*scored activity, averaged across 2.66 s. We also computed mean values of the mouse’s running speed over the same set of 2.66-s time bins. We then used the MATLAB function *fitlm()* to perform a linear regression between the resulting mean speed and mean *z-*scored activity values (**Fig. S7D**).

Analyses of neural Ca^2+^ activity from mice that performed the pellet reaching task aimed to identify cells that were activated during different phases of the behavior. We aligned neurons’ Ca^2+^ activity traces to the onset times of reaching, grasping, lifting, and eating, as determined from the mouse behavioral videos (see above). We computed trial-averaged Δ*F*(*t*)/*F*₀ traces, temporally aligned to the onset of each of these behavioral events (**Fig. 7C**). For these analyses, individual neurons in each image plane were tracked and computationally registered across recordings within and across imaging sessions (typically 3–4 sessions per mouse). We performed analyses of neuronal tuning properties by aggregating each neuron’s set of activity traces across all recordings and sessions.

To identify individual neurons that were tuned to specific behavioral phases, we evaluated mean Δ*F*/*F*₀ values averaged over the interval between the onset and offset times of each behavioral phase. We took the sets of these mean Δ*F*/*F*₀ values, determined for all behavioral trials, and performed one-tailed Wilcoxon rank-sum statistical tests to assess whether the mean Ca^2+^ activity during one behavioral phase was significantly greater than during another behavioral phase. Neurons were only identified as tuned if they had mean Ca^2+^ activity during one phase that was significantly greater (p < 0.05) than that during all other phases. Otherwise, neurons were identified as untuned.

### HCR-FISH data analyses

We exported the confocal fluorescence images from a custom Leica format into TIFF format using Leica LAS software. Using a customized Icy toolbox^172^, we aligned the fluorescence images from different rounds of HCR labeling to the *in vivo* frame-averaged image. We used Fiji (ImageJ) to rotate and crop the confocal images to the corresponding field-of-view imaged *in vivo.* To determine the spatial filters of cells in the confocal images, we used Cellpose^171^ to segment the cells, deleting identified cells that were obviously spurious and adding cells that the algorithm had obviously missed. For the datasets of **Figs. 1C–E** and **2** used as ground-truth, we used Fiji to visually proofread the resulting spatial filters and register them in the green channel side-by-side to the spatial filters determined for the *in vivo* imaging datasets with Cellpose and/or EXTRACT^168^. To quantify the alignment fidelity, we then visually checked whether or not each pair of registered cells both expressed the static red fluorescent marker.

### MERFISH data analyses

We processed MERFISH imaging data by using the Vizgen Post-processing Tool (VPT, Vizgen) for cell segmentation and Vizualizer software (Vizgen) for data visualization and export. The VPT pipeline integrated Cellpose^171^ for drawing cell boundaries based on DAPI and PolyA signals, yielding a cell-by-gene matrix that held the expression levels of each gene for each cell in its matrix entries. VPT generated an updated .vzg file with the segmentation results from Cellpose, which we opened in Vizualizer. Through a process of side-by-side visual inspection, in which we compared the MERFISH image (displayed in Vizualizer) and a corresponding *in vivo* image (displayed in Fiji), we used Vizualizer software to crop a region-of-interest within the MERFISH image corresponding to the tissue region imaged *in vivo.* To transform the centroids of the cells segmented by Cellpose to this cropped MERFISH image, we used Vizualizer to create an 8-sided polygon in the uncropped MERFISH image and we exported the vertices of this polygon to the cropped image. We used the MATLAB function *cpselect()* to situate the cell centroids within the cropped MERFISH image by applying an affine transformation between the uncropped and cropped MERFISH images using the vertices of the 8-sided polygon as control points for the transformation.

To determine the subtypes of individual motor cortical neurons, we assessed their transcriptomic signatures using the MapMyCells portal (RRID:SCR_024672, Hierarchical mapping) to map the cells’ gene expression profiles onto the Allen Brain Institute whole mouse brain database. After uploading the cell-by-gene matrix to the portal, we received its cell-type and subtype assignments. After identifying neural subtypes, we used gene expression levels, as determined by MERFISH, to create dot plots (**Fig. 6D**), in which each dot denotes the mean, log-normalized expression level of a specific gene, averaged over all cells of an individual neuron-type.

### Cell registration

To register pairs of matched cells across two different cell maps, we developed the Soma-print algorithm. The basic idea motivating the algorithm is that registering cell identities across maps is likely to be more robust when performed in a high-dimensional space capturing the geometric relationships between an individual cell and multiple of its nearest neighbors. To create such a high-dimensional space, we considered the space defined by a set of two-dimensional vectors, with each vector originating at a specific cell and extending to one of its multiple different neighboring cells. We termed this set of vectors a cell’s ‘Soma-print’ (akin to a fingerprint), as it is likely to be highly distinctive from the Soma-prints of other cells. We identified matching cell pairs across two different cell maps as those cell pairs with a high degree of concordance between their sets of such vectors, as quantified via a suitable metric. By determining the empirical distributions of this metric for best-matched and 2^nd^-best-matched pairs of cells, we estimated the relative likelihoods that cell pairings were incorrectly *vs.* correctly registered.

To enact this approach, we: (i) segmented cells in both the *in vivo* and *ex vivo* datasets to generate cell maps; (ii) performed a pre-alignment step involving an affine transformation to align the corresponding *in vivo* and *ex vivo* images; and (iii) applied the Soma-print algorithm to register cells across the aligned images. We developed two different versions of the Soma-print algorithm, the 2D and 3D Soma-print versions, which we deployed according to whether or not the *in vivo* imaging dataset sampled cells from more than one axial plane in tissue. In other words, in cases when the *in vivo* imaging field-of-view had substantial field curvature or extended depth of field, we used the 3D Soma-print variant for cell registration.

#### Cell segmentation

We generated cell maps by identifying regions-of-interest (ROIs) for individual cells in corresponding pairs of images, by either manually drawing ellipses around the cell bodies, or with two published algorithms providing cell maps, EXTRACT^168^ and Cellpose^171,173^, which respectively segment cells based on their Ca^2+^ activity patterns and morphology.

#### Pre-alignment

We first used our written records and photographs of image landmarks and the asymmetrically cut tissue block (see above) to perform an approximate, visual alignment of the corresponding pairs of *in vivo* and *ex vivo* images. (N.B: For the analyses of **Figs. 4** and **S5**, we performed this initial visual alignment prior to cell segmentation, owing to the large field-of-view for *in vivo* Ca^2+^-imaging). To refine this approximate visual alignment, we used the MATLAB function *fitgeotrans()* to perform an affine transformation that used prominent spatial landmarks, such as blood vessels and cells selected using the MATLAB function *cpselect()*, to register the image pairs.

#### 2D-Soma-print algorithm

We wrote the Soma-print algorithm to identify best-matched cell pairs in an iterative manner. This involved a unique initial round of computation, followed by subsequent rounds of cell pair identification and culling. We generally performed 3 total rounds of computation, all based on the cells’ Soma-print vector sets.

Using the registered pairs of *in vivo* and *ex vivo* images provided by the pre-alignment process, we computed a Soma-print for each cell in each of the two images by determining an *m-*fold set of two-dimensional vectors directed from the centroid of each cell’s soma to the centroids of *m* of the cell’s neighboring cells, where *m* was typically chosen to be 15 for *in vivo* images and 15–30 for *ex vivo* images. The reason we sometimes chose greater values of *m* for *ex vivo* images than for *in vivo* Ca^2+^ images was that the former generally revealed a much greater number of cells, up to about 2-fold more, than the corresponding *in vivo* Ca^2+^ images. To evaluate candidate cell pairs, in an initial round of computation, we selected *n* pairs of vectors drawn from the larger set of *m* vectors chosen for each cell, with the number of pairs typically set to *n=*10. This step was designed to address the fact that many of the cells in the *ex vivo* cell maps were unlikely to appear in the corresponding *in vivo* cell maps. To perform the selection, for each candidate cell pair, we chose the *n* vector pairs with the greatest similarity, determined using the Euclidean distance as a similarity metric. (Notably, we found with experience that the final sets of matched cell pairs provided by the Soma-print algorithm were not highly sensitive to the choice of *n* (**Fig. S3E, F**)). We took the mean value, 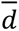, of the Euclidean distance for all *n* vector pairs and converted this into a score ranging from 0–100, using 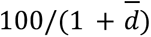. We measured 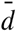 in units of image pixels, implying that the values of Soma-print similarity scores cannot be meaningfully compared across imaging workflows using distinct pixel sizes. We then multiplied each score by a weighting factor that ranged from 0–1 and depended on the spatial separation, *x*, between the two cells, computed as 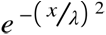, where λ was one tenth of the span of the entire field-of-view. We termed the value, *S*, of the weighted score the ‘Soma-print similarity score’ for the two cells. The weighting factor made it less likely that two cells situated far away in the image field would ultimately be identified as matching.

Given the Soma-print scores for all candidate cell pairs across the two cell maps, we computed two different statistical distributions, namely the distributions of scores for all best-matched and all 2^nd^-best-matched cell pairs, as determined for all cells from the *in vivo* cell map. The latter distribution characterizing mismatched pairs typically approximated a normal distribution, so we parametrically fit it to a Gaussian function. The former distribution typically approximated a sum of two normal distributions, which we interpreted as arising from correctly matched and incorrectly matched cell pairs, respectively. To this distribution we parametrically fit a Gaussian mixture model comprising a sum of two Gaussians with net unity area. Using the two different parametric fits, for each candidate cell pair we computed the likelihood ratio that the cell pair was correctly matched, using the ratio of likelihoods that the cell pair’s Soma-print score reflected a correct match (as computed using the Gaussian mixture model), versus an incorrect match (computed using the Gaussian fit to the score distribution for the 2^nd^-best-matched cell pairs). We determined the set of matched cell pairs from the first round of computation as those pairs for which the likelihood ratio of an incorrect match was <0.05. The p-value of each match is determined as the probability that the parametrically fit score distribution for 2^nd^-best-matched cell pairs would yield a score at least as great as the observed value. Following an empirical Bayes approach^89,90^, the *a posteriori* probability that a given cell match is incorrect is computed using Bayes’ rule based on the likelihood ratio of an incorrect *vs*. a correct match and the relative amplitudes of the two Gaussians in the mixture model.

In the algorithm’s second round, we recomputed the Soma-print for each cell in the *in vivo* map, this time by using the *n* nearest cells in the *in vivo* map that had been found in the first round of computation to have correct matches in the *ex vivo* map. The motivation for this step was that the restriction to well-matched cells would likely yield a more trustworthy Soma-print. Using this updated set of Soma-prints for cells in the *in vivo map*, we re-computed the Soma-print scores across all possible pairs of cells, with one cell in each pair taken from the *in vivo* map and one from the *ex vivo* map. To do this, for each cell in the *in vivo* map, we first computed the Soma-print for every cell in the *ex vivo* map using the *ex vivo* centroid positions of the *n* cells corresponding to those used to create the Soma-print of the *in vivo* cell. After computing Soma-print scores in this way for all possible pairs of cells across the two maps, we recomputed the likelihood ratios as described above and determined well-matched cell pairs as those for which the relative likelihood of an incorrect match was <0.05.

In the subsequent rounds of computation, we iterated the calculations, using the methods described above for round two, and thereby determined a near-final set of well-matched cells. We terminated the iterations when no more than 5% additional cells were newly matched relative to the previous iteration. Empirically, we found that using additional rounds of computation beyond three did not meaningfully alter the results for most datasets we tested (**Fig. S3H**).

After the last round of iteration computation, we curated the near-final list of matched cell pairs to determine the final set of matched cell pairs. To do this, for each cell pair on the list, we re-computed the likelihood ratio of an incorrect match, this time using the Gaussian mixture model to determine the likelihood of a correct versus an incorrect match, unlike the computations performed earlier in which we used the distribution of 2nd-best-matched Soma-print scores to determine the likelihood of an incorrect match. The reason for making this switch for the final determinations of matched cell pairs is that it is the more conservative of the two methods. The final set of matched cell pairs contained all pairs for which the likelihood ratio of an incorrect match was <0.05. (In rare cases in which there were few unmatched cells, the Gaussian mixture model became statistically indistinguishable from a unimodal Gaussian function, as determined using the Bayesian Information Criterion (BIC). In these cases, we reverted to the initial method of computing the likelihood of an incorrect match using the distribution of 2nd-best-matched scores).

Runtimes for each round of computation typically ranged from a few minutes to tens of minutes when using a conventional desktop computer (Intel(R) processor, Core(TM) i9-10900X CPU, 3.7 GHz clock speed, 256 GB of RAM).

#### 3D Soma-print algorithm

For mice in which we acquired large-scale *in vivo* imaging datasets (**Figs. 4, S5**) or performed imaging through a GRIN microlens (**Figs. 5, S6**), owing to the substantial field curvature of the optical focal plane, the resulting *in vivo* image frames could not be aligned well to a single image plane from the stack of processed optical sections (see ***Confocal microscopy*** section above regarding the image processing) acquired from a tissue block *ex vivo*. Even in the absence of such field curvature, other cases in which the *in vivo* images spanned multiple *ex vivo* image planes were those in which human errors in tissue handling led to suboptimal optomechanical alignments (**Fig. 2L–N**). To address all these cases, we developed the 3D Soma-print approach to register *in vivo* images to multiplane image stacks acquired *ex vivo*.

We took pre-aligned pairs of *in vivo* and *ex vivo* images and used the basic 2D-Soma-print methods described above to compute Soma-print scores for all pairs of cells across the *in vivo* image and each of the *ex vivo* image slices. (For GRIN lens datasets, owing to the substantial field curvature and worse axial resolution of the *in vivo* image, instead of using individual image slices, we used maximum axial projections across 3 or 4 adjacent *ex vivo* image slices). Most cells found *in vivo* appeared in more than one *ex vivo* image slice (or projection), yielding 2D-Soma-print matches across more than one image slice. From the set of all such matches determined with the 2D-Soma-print method, we determined the best match in 3D by taking the image slice and *ex vivo* cell yielding the maximum Soma-print score across all the candidate matched cells from all image slices.

We then used the *ex vivo* image stack to reconstruct a 2D-manifold that best approximated the field-of-view seen in the *in vivo* imaging data (**Figs. 2O**, **4B**, **4C**, **5A**). To do this, we used the MATLAB function, *tpaps()*, to fit a thin-plate smoothing spline, *z_fit_* = *f*(*x*,*y*), to the axial location of all matched cells as a function of their lateral (*x*, *y*) coordinates. We then computed an axial, maximum projection image using the 3 or 4 axial planes in the *ex vivo* image stack that were closest to the fit manifold, *z_fit_* = *f*(*x*, *y*). For a final set of statistical characterizations, we applied the 2D-Soma-print algorithm to this maximum projection image and the *in vivo* image. We verified that precision and recall values for cell matches obtained from this final 2D-Soma-print analysis were not meaningfully different from those obtained by taking the best match in 3D, as described in the prior paragraph.

### Analyses of Ca^2+^ activity in motor cortical neurons with defined RNA or projectomic signatures

To compare Ca²⁺ activity patterns across different transcriptomic and projectomic classes of neurons, we analyzed data from mice performing a pellet-reaching task. We first obtained neurons’ spatial masks from the Ca²⁺ movies using EXTRACT^168^ and from *ex vivo* HCR-FISH images using Cellpose^171^. We then registered neurons using the Soma-print algorithm. For each registered cell, we quantified expression for each gene or barcode label by averaging HCR-FISH fluorescence intensities within the neuron’s Cellpose mask. We focused on gene markers and projection pathways previously indicated to distinguish between known subclasses of layer 5 motor cortical neurons. We assigned each neuron to a marker-positive (+) or marker-negative (–) group using empirically selected intensity thresholds that reproduced the orthogonal subclass patterns in published reference datasets^21^.

After identifying cells with specific gene markers or projection labels, we analyzed each cell’s behaviorally related Ca^2+^ dynamics using custom MATLAB scripts (MATLAB 2023a, MathWorks). We z-scored each neuron’s raw fluorescence trace within the recording session by subtracting the mean fluorescence level and dividing by the s.d. of the baseline fluctuations. We smoothed the individual traces using a sliding filter with a 1-s-window and computed trial-averaged traces for each neuron, temporally aligned to grasp onset, with each trial corresponding to a single pellet-reaching attempt.

Using the resulting trial-averaged, z-scored traces, we quantified the response dynamics of individual neurons by computing their: 1) peak z-scored Ca^2+^ activity within the interval [–1 s, 4 s] relative to grasp onset; 2) mean time to achieve the Ca^2+^ activity peak relative to the time of grasp onset; 3) FWHM of the Ca^2+^ transient evoked at grasp onset, determined as the interval duration over which the z-scored trace exceeded half the amplitude between the pre-event baseline, determined across the interval [–6 s, –1 s] relative to grasp onset, and the peak value (**Figs. 7F, G**).

To evaluate differences in neurons’ behaviorally evoked Ca^2+^ dynamics between cells that did or did not exhibit a given gene or projectomic signature, we used a permutation test to evaluate the significance of a specific facet of z-scored Ca^2+^ activity, *viz.*, the peak amplitude, time-to-peak, or FWHM values calculated as described above. To generate a statistical null distribution, we randomly shuffled the cell labels across all cells in the dataset. We then computed the absolute mean difference for the quantity of interest between its value in the real data and in 100,000 different shuffled datasets. We determined the p-value of the permutation test by computing the proportion of shuffled datasets for which the absolute mean difference across the ‘positive’ and ‘negative’ groups met or exceeded the observed difference in the real data. To account for multiple pairwise comparisons, we applied a Dunn-Sidak correction to the relevant p-values.

To characterize the kinetics of individual Ca^2+^ transients, we first detected discrete Ca^2+^ events from the fluorescence traces (Δ*F*/*F*_0_) using a threshold-based method, requiring that the trace stay >2 s.d. above the mean baseline level for at least 200 ms. To ensure that our analyses focused on well-isolated Ca^2+^ transients, we excluded any Ca^2+^ event occurring within 1 s of a preceding event in the same neuron. For each valid Ca^2+^ event, we examined the trace between the interval [–1 s, 5 s] relative to the time of the detected Ca^2+^ event, and we fit the decay phase (from the peak to the end of the 6-s-interval) with a single-exponential function (𝐴 𝑒^−𝑡/τ^), using a nonlinear least-squares regression to estimate the amplitude (𝐴) and decay time constant (τ). We used the results of the individual fits in further analyses only if the coefficient of determination (𝑅^2^) exceeded 0.7. We grouped the resulting kinetic parameters by cell-type, and assessed statistical significance between groups using a Kruskal-Wallis ANOVA, followed by post hoc Wilcoxon rank-sum tests with a Bonferroni correction for multiple pairwise comparisons.

#### Statistics

We used MATLAB (MATLAB versions 2023 and 2024; Mathworks) to perform linear regression and nonparametric ANOVA analyses, as well as permutation and rank sum tests for comparisons between groups. Tests were two-tailed except as noted above.

## Acknowledgments

We gratefully acknowledge research funding to M.J.S. from HHMI, the Carol & Gene Ludwig Family Foundation, the Simons Foundation, the Stanford Knight Initiative for Brain Resilience, and a Vannevar Bush Faculty Fellowship from the U.S. Department of Defense, a Jane Coffin Childs postdoctoral fellowship to X.C.S., and an American Academy of Neurology Neuroscience Research Training Scholarship and a Chan Zuckerberg Biohub Physician Scientist Fellowship to G.C. Confocal imaging was done at the Stanford Cell Sciences Imaging Core Facility (RRID:SCR_017787) and the Wu Tsai Neuroscience Microscopy Service. We thank TRANSVISTA and Coherent Corp. for providing instrumentation enabling our experiments with the miniature two-photon microscope. We thank Jizhou Li, Ritchie Chen, and Jin Wang for technical discussions and Scott Sternson for inspirational conversation near the start of the project.

